# Accuracy, Robustness and Scalability of Dimensionality Reduction Methods for Single Cell RNAseq Analysis

**DOI:** 10.1101/641142

**Authors:** Shiquan Sun, Jiaqiang Zhu, Ying Ma, Xiang Zhou

## Abstract

**Background:** Dimensionality reduction (DR) is an indispensable analytic component for many areas of single cell RNA sequencing (scRNAseq) data analysis. Proper DR can allow for effective noise removal and facilitate many downstream analyses that include cell clustering and lineage reconstruction. Unfortunately, despite the critical importance of DR in scRNAseq analysis and the vast number of DR methods developed for scRNAseq studies, however, few comprehensive comparison studies have been performed to evaluate the effectiveness of different DR methods in scRNAseq.

**Results:** Here, we aim to fill this critical knowledge gap by providing a comparative evaluation of a variety of commonly used DR methods for scRNAseq studies. Specifically, we compared 18 different DR methods on 30 publicly available scRNAseq data sets that cover a range of sequencing techniques and sample sizes. We evaluated the performance of different DR methods for neighborhood preserving in terms of their ability to recover features of the original expression matrix, and for cell clustering and lineage reconstruction in terms of their accuracy and robustness. We also evaluated the computational scalability of different DR methods by recording their computational cost.

**Conclusions:** Based on the comprehensive evaluation results, we provide important guidelines for choosing DR methods for scRNAseq data analysis. We also provide all analysis scripts used in the present study at www.xzlab.org/reproduce.html. Together, we hope that our results will serve as an important practical reference for practitioners to choose DR methods in the field of scRNAseq analysis.

## INTRODUCTION

Single-cell RNA sequencing (scRNAseq) is a rapidly growing and widely applying technology [1–3]. By measuring gene expression at single cell level, scRNAseq provides an unprecedented opportunity to investigate the cellular heterogeneity of complex tissues [4–8]. However, despite the popularity of scRNAseq, analyzing scRNAseq data remains a challenging task. Specifically, due to the low capture efficiency and low sequencing depth per cell in scRNAseq data, gene expression measurements obtained from scRNAseq are noisy: collected scRNAseq gene measurements are often in the form of low expression counts, and in studies not based on unique molecular identifiers, are also paired with an excessive number of zeros known as dropouts [9]. Subsequently, dimensionality reduction (DR) methods that transform the original high-dimensional noisy expression matrix into a low-dimensional subspace with enriched signals become an important data processing step for scRNAseq analysis [10]. Proper DR can allow for effective noise removal, facilitate data visualization, and enable efficient and effective downstream analysis of scRNAseq [11].

DR is indispensable for many types of scRNAseq analysis. Because of the importance of DR in scRNAseq analysis, many DR methods have been developed and are routinely used in scRNAseq software tools that include, but not limited to, cell clustering tools [12, 13] and lineage reconstruction tools [14]. Indeed, most commonly used scRNAseq clustering methods rely on DR as the first analytic step [15]. For example, Seurat applies clustering algorithms directly on a low dimensional space inferred from principal component analysis (PCA) [16]. CIDR improves clustering by improving PCA through imputation [17]. SC3 combines different ways of PCA for consensus clustering [18]. Besides PCA, other DR techniques are also commonly used for cell clustering. For example, nonnegative matrix factorization (NMF) is used in SOUP [19]. Partial least squares is used in scPLS [20]. Diffusion map is used in destiny [21]. Multidimensional scaling (MSD) is used in ascend [22]. Variational inference autoencoder is used in scVI [23]. In addition to cell clustering, most cell lineage reconstruction and developmental trajectory inference algorithms also rely on DR [14]. For example, TSCAN builds cell lineages using minimum spanning tree based on a low dimensional PCA space [24]. Waterfall performs *k*-means clustering in the PCA space to eventually produce linear trajectories [25]. SLICER uses locally linear embedding (LLE) to project the set of cells into a lower dimension space for reconstructing complex cellular trajectories [26]. Monocle employs either independent components analysis (ICA) or uniform manifold approximation and projection (UMAP) for DR before building the trajectory [27, 28]. Wishbone combines PCA and diffusion maps to allow for bifurcation trajectories [29].

Besides the generic DR methods mentioned in the above paragraph, many DR methods have also been developed recently that are specifically targeted for modeling scRNAseq data. These scRNAseq specific DR methods can account for either the count nature of scRNAseq data and/or the dropout events commonly encountered in scRNAseq studies. For example, ZIFA relies on a zero-inflation normal model to model dropout events [30]. pCMF models both dropout events and the mean-variance dependence resulting from the count nature of scRNAseq data [31]. ZINB-WaVE incorporates additional gene-level and sample-level covariates for more accurate DR [32]. Finally, several deep learning-based DR methods have recently been developed to enable scalable and effective computation in large-scale scRNAseq data, including data that are collected by 10X Genomics techniques [33] and/or from large consortium studies such as Human Cell Atlas (HCA) [34, 35]. Common deep learning-based DR methods for scRNAseq include Dhaka [36], scScope [37], VASC [38], scvis [39], and DCA [40], to name a few.

With all these different DR methods for scRNAseq data analysis, one naturally wonders which DR method one would prefer for different types of scRNAseq analysis. Unfortunately, despite the popularity of scRNAseq technique, the critical importance of DR in scRNAseq analysis, and the vast number of DR methods developed for scRNAseq studies, few comprehensive comparison studies have been performed to evaluate the effectiveness of different DR methods for practical applications. Here, we aim to fill this critical knowledge gap by providing a comprehensive comparative evaluation of a variety of commonly used DR methods for scRNAseq studies. Specifically, we compared 18 different DR methods on 30 publicly available scRNAseq data sets that cover a range of sequencing techniques and sample sizes [12, 14, 41]. We evaluated the performance of different DR methods for neighborhood preserving in terms of their ability to recover features of the original expression matrix, and for cell clustering and lineage reconstruction in terms of their accuracy and robustness using different metrics. We also evaluated the computational scalability of different DR methods by recording their computational time. Together, we hope our results can serve as an important guideline for practitioners to choose DR methods in the field of scRNAseq analysis.

## RESULTS

We evaluated the performance of 18 DR methods (Table 1; Figure S1) on 30 publicly available scRNAseq data sets (Tables S1-S2) and 2 simulated data sets. Details of these data sets are provided in Methods and Materials. Briefly, these data sets cover a wide variety of sequencing techniques that include Smart-Seq2 (8 data sets), Smart-Seq (5 data sets), 10X genomics (6 data sets), inDrop (1 data set), RamDA-seq (1 data set), sci-RNA-seq3 (1 data set), SMARTer (5 data sets) and others (3 data sets). In addition, these data sets cover a range of sample sizes from a couple of hundred cells to over tens of thousands of cells. In each data set, we evaluated the ability of different DR methods in preserving the original feature of the expression matrix, and, more importantly, their effectiveness for two important single cell analytic tasks: cell clustering and lineage inference. In particular, we used 14 real data sets together with 2 simulated data sets for DR method comparison in terms of cell clustering performance. We used the another a set of 14 real data sets for DR method comparison in terms of trajectory inference. We used yet two additional large-scale scRNAseq data sets to examine the effectiveness and scalability of different DR methods there. In addition, we measured the computing stability of different DR methods and recorded their computation time. An overview of the comparison workflow is shown in Figure 1. Because common tSNE software can only extract a small number low-dimensional components [42–44], we only included tSNE results based on two low-dimensional components extracted from the recently developed fast *FIt-SNE* R package [44] in all figures. All data and analysis scripts for reproducing the results in the paper is available at www.xzlab.org/reproduce.html or https://github.com/xzhoulab/DRComparison.

**Figure 1.**
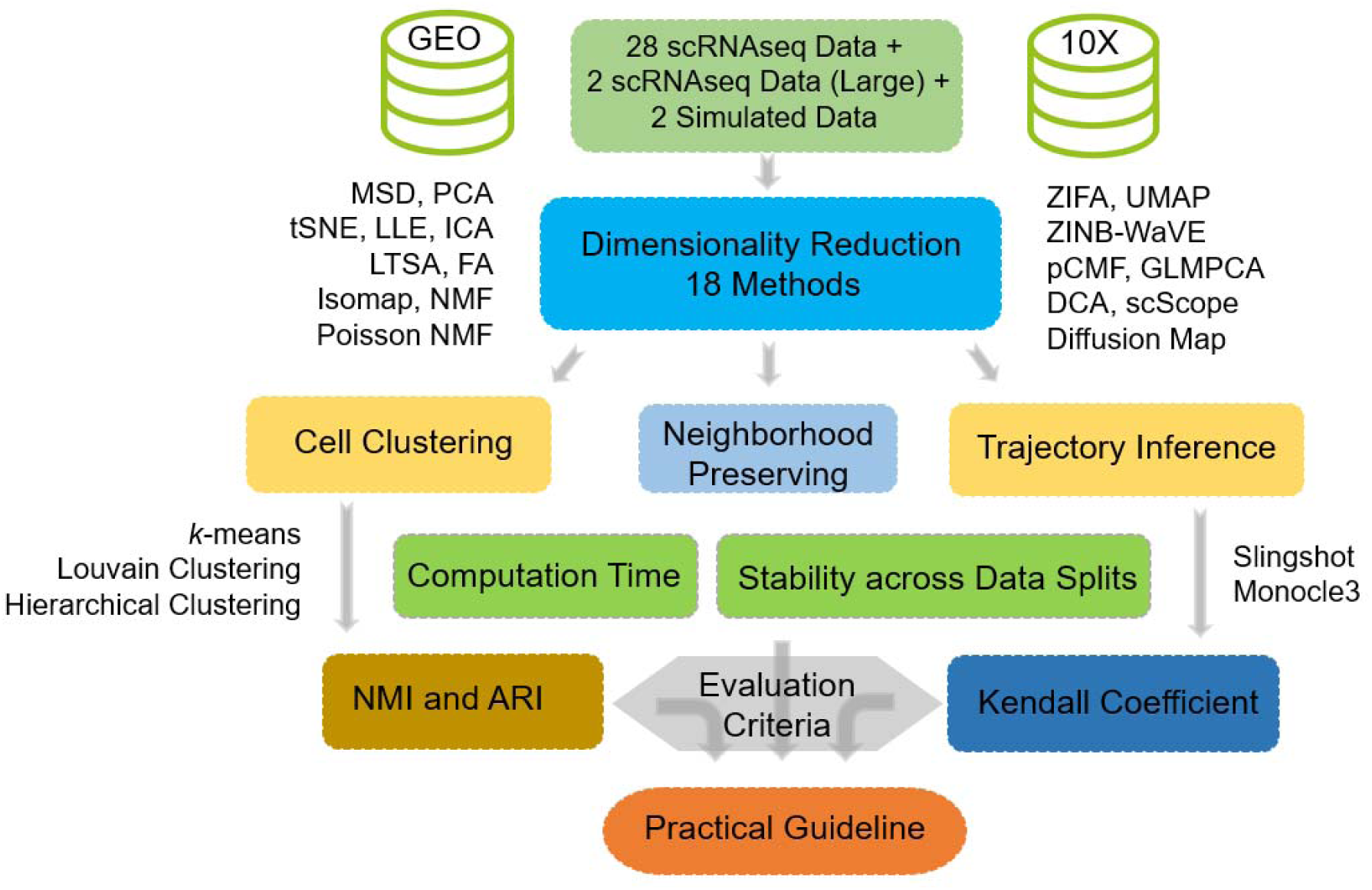
Overview of the evaluation workflow for dimensionality reduction methods. We obtained a total of 30 publicly available scRNAseq data from GEO and 10x Genomics website. We also simulated two addition simulation data sets. For each of the 32 data sets in turn, we applied 18 dimensionality reduction (DR) methods to extract the low-dimensional components. Afterwards, we evaluated the performance of DR methods by evaluating how effective the low-dimensional components extracted from DR methods are for downstream analysis. We did so by evaluating the two commonly applied downstream analysis: clustering analysis and lineage reconstruction analysis. In the analysis, we varied the number of low-dimensional components extracted from these DR methods. The performance of each DR method is qualified by Jaccard index for neighborhood preserving, normalized mutual information (NMI) and adjusted rand index (ARI) for cell clustering analysis, and Kendall correlation coefficient for trajectory inference. We also recorded the stability of each DR method across data splits and recorded the computation time for each DR method. Through the comprehensive evaluation, we eventually provide practical guidelines for practitioners to choose DR methods for scRNAseq data analysis.

**Table 1.**
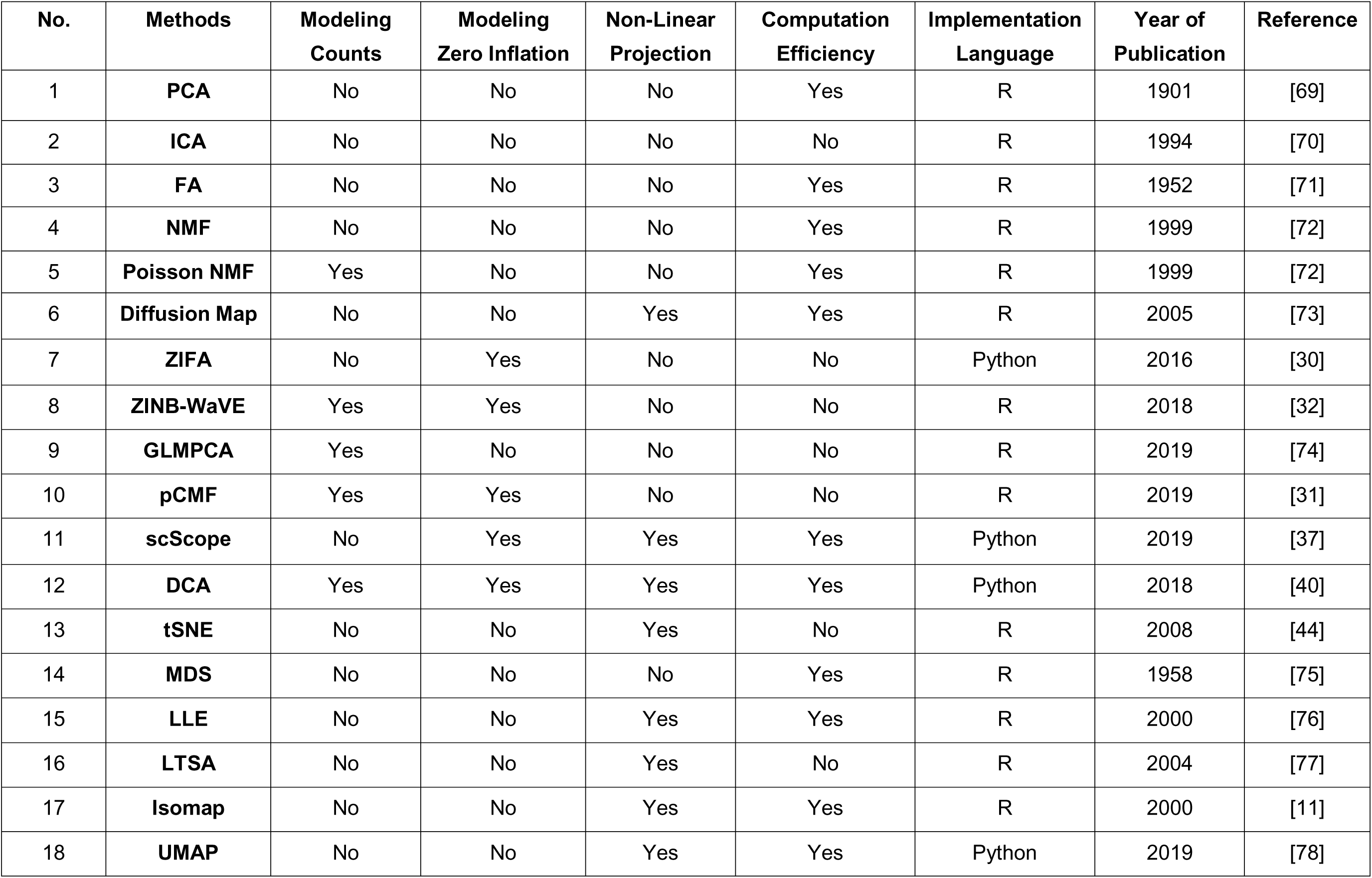
List of compared dimensionality reduction methods. We list standard modeling properties for each of compared dimensionality reduction methods. These properties include whether it models count data (3^rd^ column), whether it accounts for zero inflation (4^th^ column), whether it is a linear DR method (5^th^ column), its computation efficiency (6^th^ column), implementation language (7^th^ column), year of publication (8^th^ column), and reference (9^th^ column). FA: factor analysis; PCA: principal component analysis; ICA: independent component analysis; NMF: nonnegative matrix factorization; Poisson NMF: Kullback-Leibler divergence-based NMF; ZIFA: zero-inflated factor analysis; ZINB-WaVE: zero-inflated negative binomial based wanted variation extraction; pCMF: probabilistic count matrix factorization; DCA: deep count autoencoder network; GLMPCA: generalized linear model principal component analysis; Diffusion Map; MDS: multidimensional scaling; LLE: locally linear embedding, LTSA: local tangent space alignment; Isomap; UMAP: uniform manifold approximation and projection; tSNE: t-distributed stochastic neighbor embedding.

### Performance of DR methods for neighborhood preserving

We first evaluated the performance of different DR methods in terms of preserving the original features of the gene expression matrix. To do so, we applied different DR methods to each of 30 scRNAseq data sets (28 real data and 2 simulated data; excluding the two large-scale data due to computing concerns) and evaluated the performance of these DR methods based on neighborhood preserving. Neighborhood preserving measures how the local neighborhood structure in the reduced dimensional space resembles that in the original space by computing a Jaccard index [45] (details in Methods and Materials). In the analysis, for each DR method and each scRNAseq data set, we applied the DR method to extract a fixed number of low-dimensional components (e.g. these are the principal components in the case of PCA). We varied the number of low-dimensional components to examine their influence on local neighborhood preserving. Specifically, for each of 16 cell clustering data sets, we varied the number of low dimensional components to be either 2, 6, 14, or 20 when the data contains less than or equal to 300 cells, and we varied the number of low dimensional components to be either 0.5%, 1%, 2%, or 3% of the total number of cells when the data contains more than 300 cells. For each of the 14 trajectory inference data sets, we varied the number of low-dimensional components to be either 2, 6, 14, or 20 regardless of the number of cells. Finally, we also varied the number of neighborhood cells used in the Jaccard index to be either 10, 20, or 30. The evaluation results based on the Jaccard index of neighborhood preserving are summarized in Figures S2-S14.

In the cell clustering data sets, we found that pCMF achieves the best performance of neighborhood preserving across all data sets and across all included low-dimensional components (Figures S2-S7). For example, with 30 neighborhood cells and 0.5% of low-dimensional components, pCMF achieves a Jaccard index of 0.25. Its performance is followed by Poisson NMF (0.16), ZINB-WaVE (0.16), Diffusion Map (0.16), MDS (0.15), and tSNE (0.14). While the remaining two methods, scScope (0.1) and LTSA (0.06), do not fare well. Increasing number of neighborhood cells increases the absolute value of Jaccard index but does not influence the relative performance of DR methods (Figure S7). In addition, the relative performance of most DR methods remains largely similarly whether we focus on data sets with unique molecular identifiers (UMI) or data sets without UMI (Figure S8). However, we do notice two exceptions: the performance of pCMF decreases with increasing number of low-dimensional components in UMI data but increases in non-UMI data; the performance of scScope is higher in UMI data than its performance in non-UMI data. In the trajectory inference data sets, pCMF again achieves the best performance of neighborhood preserving across all data sets and across all included low-dimensional components (Figures S9-S14). Its performance is followed closely by scScope and Poisson NMF. For example, with 30 neighborhood cells and 20 low-dimensional components, the Jaccard index of pCMF, Poisson NMF, and scScope across all data sets are 0.3, 0.28, and 0.26, respectively. Their performance is followed by ZINB-WaVE (0.19), FA (0.18), ZIFA (0.18), GLMPCA (0.18), and MDS (0.18). In contrast, LTSA also does not fare well across all included low-dimensional components (Figure S14). Again, increasing number of neighborhood cells increases the absolute value of Jaccard index but does not influence the relative performance among DR methods (Figures S9-S14).

We note that the measurement we used in this subsection, neighborhood preserving, is purely for measuring DR performance in terms of preserving the original gene expression matrix and may not be relevant for single cell analytic tasks that are the main focus of the present study: a DR method that preserves the original gene expression matrix may not be effective in extracting useful biological information from the expression matrix that is essential for key downstream single cell applications. Preserving the original gene expression matrix is rarely the sole purpose of DR methods for single cell applications: indeed, the original gene expression matrix (which is the best-preserved matrix of itself) is rarely, if ever, used directly in any downstream single cell applications including clustering and lineage inference, even though it is computationally easy to do so. Therefore, we will focus our main comparison in two important downstream single cell applications listed below.

### Performance of DR methods for cell clustering

As our main comparison, we first evaluated the performance of different DR methods for cell clustering applications. To do so, we obtained 14 publicly available scRNAseq data sets and simulated two additional scRNAseq data sets using the *Splatter* package (Table S1). Each of the 14 real scRNAseq data sets contains known cell clustering information while each of the 2 simulated data sets contains 4 or 8 known cell types. For each DR method and each data set, we applied DR to extract a fixed number of low-dimensional components (e.g., these are the principal components in the case of PCA). We again varied the number of low-dimensional components as in the previous section to examine their influence on cell clustering analysis. We then applied either the hierarchical clustering method, the *k*-means clustering method, or Louvain clustering method [46] to obtain the inferred cluster labels. We used both normalized mutual information (NMI) and adjusted rand index (ARI) values for comparing the true cell labels and inferred cell labels obtained by clustering methods based on the low-dimensional components.

#### Cell clustering with different clustering methods

The evaluation results on DR methods based on clustering analysis using the *k*-means clustering algorithm are summarized in Figure 2 (for NMI criterion) and Figure S15 (for ARI criterion). Because the results based on either of the two criteria are similar, we will mainly explain the results based on the NMI criteria in Figure 2. For easy visualization, we also display the results averaged across data sets in Figure S16. A few patterns are noticeable. First, as one would expect, clustering accuracy depends on the number of low-dimensional components that are used for clustering. Specifically, accuracy is relatively low when the number of included low-dimensional components is very small (e.g. 2 or 0.5%) and generally increases with the number of included components. In addition, accuracy usually saturates once a sufficient number of components is included, though the saturation number of components can vary across data sets and across methods. For example, the average NMI across all data sets and across all methods are 0.61, 0.66, 0.67 and 0.67 for increasingly large number of components, respectively. Second, when conditional on using a low number of components, scRNAseq specific DR method ZINB-WaVE and generic DR methods ICA and MDS often outperform the other methods. For example, with the lowest number of components, the average NMI across all data sets for MDS, ICA and ZINB-WaVE are 0.82, 0.77 and 0.76, respectively (Figure S16A). The performance of MDS, ICA and ZINB-WaVE is followed by LLE (0.75), Diffusion Map (0.71), ZIFA (0.69), PCA (0.68), FA (0.68), tSNE (0.68), NMF (0.59) and DCA (0.57). While the remaining four methods, Poisson NMF (0.42), pCMF (0.41), scScope (0.26), and LTSA (0.12), do not fare well with a low number of components. Third, with increasing number of low-dimensional components, generic methods such as FA, ICA, MDS and PCA are often comparable with scRNAseq specific methods such as ZINB-WaVE. For example, with the highest number of low-dimensional components, the average NMI across all data sets for FA, ICA, PCA, ZINB-WaVE, LLE and MDS are 0.85, 0.84, 0.83, 0.83, 0.82 and 0.82, respectively. Their performance is followed by ZIFA (0.79), NMF (0.73), and DCA (0.69). The same four methods, pCMF (0.55), Poisson NMF (0.31), scScope (0.31), and LTSA (0.06) again do not fare well with a large number of low-dimensional components (Figure S16A). The comparable results of generic DR methods with scRNAseq specific DR methods with a high number of low-dimensional components are also consistent some of the previous observations; for example, the original ZINB-WaVE paper observed that PCA can generally yield comparable results with scRNAseq specific DR methods in real data [32].

**Figure 2.**
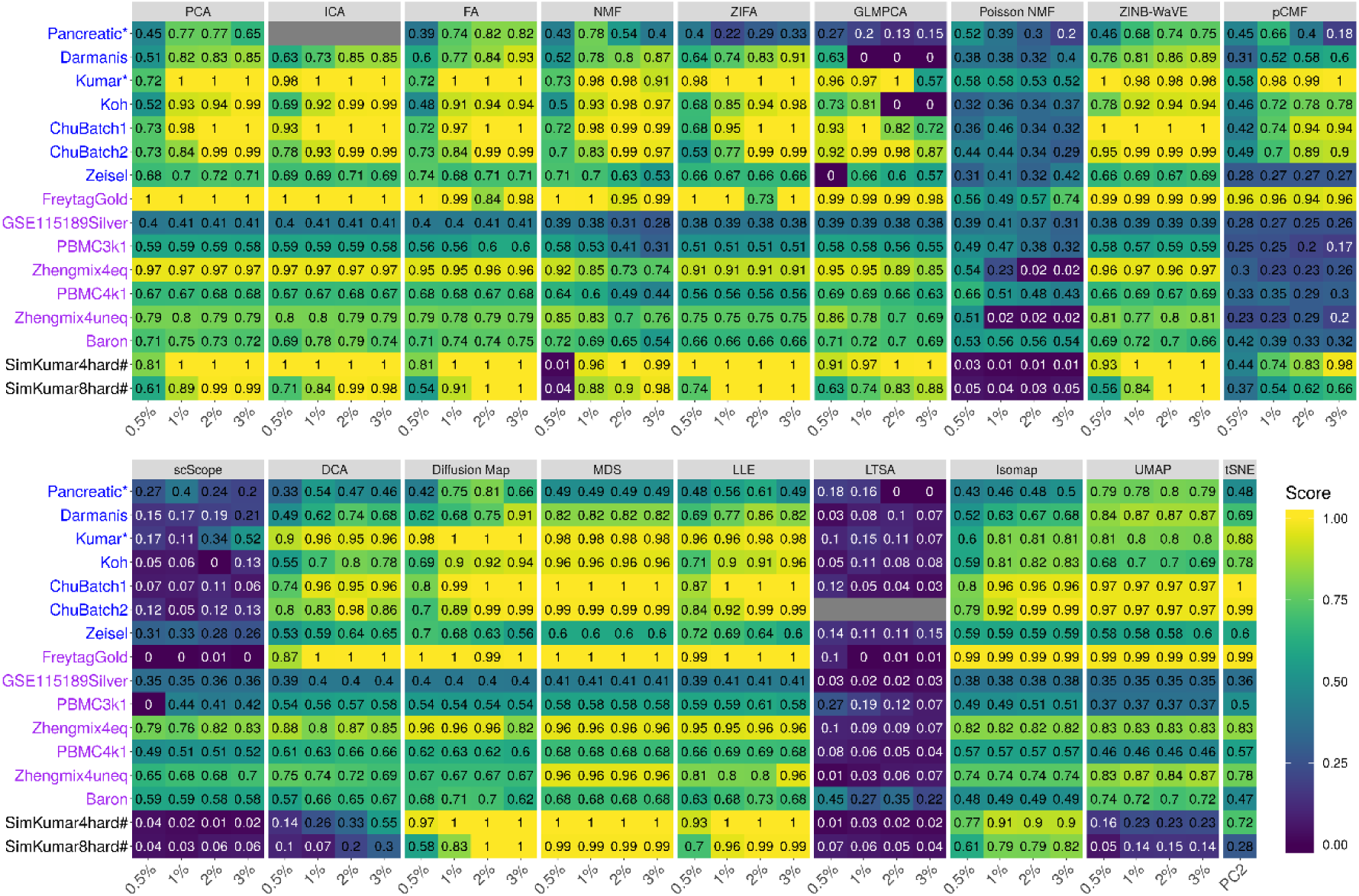
DR method performance evaluated by *k*-means clustering based on NMI in downstream cell clustering analysis. We compared 18 DR methods (columns), including factor analysis (FA), principal component analysis (PCA), independent component analysis (ICA), Diffusion Map, nonnegative matrix factorization (NMF), Poisson NMF, zero-inflated factor analysis (ZIFA), zero-inflated negative binomial based wanted variation extraction (ZINB-WaVE), probabilistic count matrix factorization (pCMF), deep count autoencoder network (DCA), scScope, generalized linear model principal component analysis (GLMPCA), multidimensional scaling (MDS), locally linear embedding (LLE), local tangent space alignment (LTSA), Isomap, uniform manifold approximation and projection (UMAP), and t-distributed stochastic neighbor embedding (tSNE). We evaluated their performance on 14 real scRNAseq data sets (UMI-based data are labeled as purple; non-UMI based data are labeled as blue) and 2 simulated data sets (rows). The simulated data based on Kumar data is labeled with #. The performance of each DR method is measured by normalized mutual information (NMI). For each data set, we compared the four different number of low-dimensional components. The four numbers equal to 0.5%, 1%, 2%, and 3% of the total number of cells in big data and equal to 2, 6, 14, and 20 in small data (which are labeled with *). For convenience, we only listed 0.5%, 1%, 2%, and 3% on x-axis. No results for ICA are shown in the table (grey fills) because ICA cannot handle the large number of features in that data. No results for LTSA are shown (grey fills) because error occurred when we applied the clustering method on LTSA extracted low-dimensional components there. Note that, for tSNE, we only extracted two low-dimensional components due to the limitation of the tSNE software.

Besides the *k*-means clustering algorithm, we also used the hierarchical clustering algorithm to evaluate the performance of different DR methods (Figures S17-S19). In this comparison, we had to exclude one DR method, scScope, as hierarchical clustering does not work on the extracted low-dimensional components from scScope. Consistent with the *k*-means clustering results, we found that the clustering accuracy measured by hierarchical clustering is relatively low when the number of low-dimensional components is very small (e.g. 2 or 0.5%), but generally increases with the number of included components. In addition, consistent with the *k*-means clustering results, we found that generic DR methods often yield results comparable to or better than scRNAseq specific DR methods (Figures S17-S19). In particular, with a low number of low-dimensional components, MDS achieves the best performance (Figure S19). With a moderate or high number of low-dimensional components, two generic DR methods, FA and NMF, often outperform various other DR methods across a range of settings. For example, when the number of low-dimensional components is moderate (6 or 1%), both FA and NMF achieve an average NMI value of 0.80 across data sets (Figure S19A). In this case, their performance is followed by PCA (0.72), Poisson NMF (0.71), ZINB-WaVE (0.71), Diffusion Map (0.70), LLE (0.70), ICA (0.69), ZIFA (0.68), pCMF (0.65), and DCA (0.63). tSNE (0.31) does not fare well, either because it only extracts two-dimensional components or because it does not pair well with hierarchical clustering. We note, however, that the clustering results obtained by hierarchical clustering are often slightly worse than that obtained by *k*-means clustering across settings (e.g., Figure S16 vs Figure S19), consistent with the fact that many scRNAseq clustering methods use *k*-means as a key ingredient [18, 25].

Finally, besides the *k*-means and hierarchical clustering methods, we also performed clustering analysis based on a community detection algorithm Louvain clustering method [46]. Unlike the *k*-means and hierarchical clustering methods, Louvain method does not require a pre-defined number of clusters and can infer the number of clusters in an automatic fashion. Following software recommendation [28, 46], we set the *k*-nearest neighbor parameter in Louvain method to be 50 for graph building in the analysis. We measured DR performance again by either average NMI (Figure S20) or ARI (Figure S21). Consistent with the *k*-means clustering results, we found that the clustering accuracy measured by Louvain method is relatively low when the number of low-dimensional components is very small (e.g. 2 or 0.5%), but generally increases with the number of included components. With a low number of low-dimensional components, ZINB-WaVE (0.72) achieves the best performance (Figures S20-S22). With a moderate or high number of low-dimensional components, two generic DR methods, FA and MDS, often outperform various other DR methods across a range of settings (Figures S20-S22). For example, when the number of low-dimensional components is high (6 or 1%), FA achieves an average NMI value of 0.77 across data sets (Figure S22A). In this case, its performance is followed by NMF (0.76), MDS (0.75), GLMPCA (0.74), LLE (0.74), PCA (0.73), ICA (0.73), ZIFA (0.72), and ZINB-WaVE (0.72). Again consistent with the *k*-means clustering results, scScope (0.32) and LTSA (0.21) do not fare well. We also note that the clustering results obtained by Louvain method are often slightly worse than that obtained by *k*-means clustering and slightly better than that obtained by hierarchical clustering across settings (e.g., Figure S16 vs Figure S19 vs Figure S22).

#### Normalization does not influence the performance of DR methods

While some DR methods (e.g. Poisson NMF, ZINB-WaVE, pCMF and DCA) directly model count data, many DR methods (e.g. PCA, ICA, FA, NMF, MDS, LLE, LTSA, Isomap, Diffusion Map, UMAP, and tSNE) require normalized data. The performance of DR methods that use normalized data may depend on how data are normalized. Therefore, we investigated how different normalization approaches impact on the performance of the aforementioned DR methods that use normalized data. We examined two alternative data transformation approaches, log2 CPM (count per million; 11 DR methods) and z-score (10 DR methods), in addition to the log2 count we used in the previous results (transformation details are provided in Methods and Materials). The evaluation results are summarized in Figures S23-S30 and are generally insensitive to the transformation approach deployed. For example, with the *k*-means clustering algorithm, when the number of low-dimensional components is small (1%), PCA achieves an NMI value of 0.82, 0.82 and 0.81, for log2 count transformation, log2 CPM transformation, and z-score transformation, respectively (Figures S16A, S26A, and S30A). Similar results hold for the hierarchical clustering algorithm (Figures S16B, S26B, and S30B) and Louvain clustering method (Figures S16C, S26C, and S30C). Therefore, different data transformation approaches do not appear to substantially influence the performance of DR methods.

#### Performance of DR methods in UMI vs non-UMI based data sets

scRNAseq data generated from UMI-based technologies (e.g., 10X genomics) are often of large scale, come with almost no amplification bias, do not display apparent dropout events, and can be accounted for by over-dispersed Poisson distributions. In contrast, data generated from non UMI-based techniques (e.g., Smart-Seq2) are often of small scale, have high capture rate, and come with excessive dropout events. Subsequently, the unwanted variation from these two types of dataset can be quite different. To investigate how different DR methods perform in these two different types of data sets, we grouped 14 cell clustering data sets into a UMI-based group (7 data sets) and a non UMI-based group (7 data sets). In the UMI-based data sets, we found that many DR methods perform reasonably well and their performance is relatively stable across a range of included low-dimensional components (Figure S31A). For example, with the lowest number of low-dimensional components, the average NMI of PCA, ICA, FA, NMF, GLMPCA, ZINB-WaVE, and MDS are 0.73, 0.73, 0.73, 0.73, 0.74, and 0.75, respectively. Their performance remains similar with increasing number of low-dimensional components. However, a few DR methods, including Poisson NMF, pCMF, scScope, and LTSA, all have extremely low performance across settings. In the non UMI-based data sets, the same set of DR methods perform reasonably well though their performance can vary with respect to the number of low-dimensional components (Figure S31B). For example, with a low number of low-dimensional components, five DR methods, MDS, UMAP, ZINB-WaVE, ICA, and tSNE, perform reasonably well. The average NMI of these methods are 0.83, 0.81, 0.80, 0.78, and 0.77, respectively. With increasing number of low-dimensional components, four additional DR methods, PCA, ICA, FA, and ZINB-WaVE, also start to catchup. However, a similar set of DR methods, including GLMPCA, Poisson NMF, scScope, LTSA, and occasionally pCMF, also do not perform well in these non-UMI data sets.

#### Visualization of clustering results

We visualized the cell clustering results in two example data sets: the Kumar data which is non-UMI based and the PBMC3k data which is UMI based. The Kumar data consists of mouse embryonic stem cells cultured in three different media while the PBMC3k data consists of 11 blood cell types (data details in the Supplementary Information). Here, we extracted 20 low-dimensional components in the Kumar data and 32 low low-dimensional components in the PBMC3k data with different DR methods. We then performed tSNE analysis on these low-dimensional components to extract the two tSNE components for visualization (Figures S32-S33). Importantly, we found that the tSNE visualization results are not always consistent with clustering performance for different DR methods. For example, in the Kumar data, the low dimensional space constructed by FA, pCMF and MDS often yield clear clustering visualization with distinguish clusters (Figure S32), consistent with their good performance in clustering (Figure 2). However, the low dimensional space constructed by PCA, ICA, and ZIFA often do not yield clear clustering visualization (Figure S32), even though these methods all achieve high cell clustering performance (Figure 2). Similarly, in the PBMC3k data set, FA and MDS perform well in clustering visualization (Figure S33), which is consistent with their good performance in clustering analysis (Figure 2). However, PCA and ICA do not fare well in clustering visualization (Figure S33), even though both of them achieve high clustering performance (Figure 2). The inconsistency between cluster visualization and clustering performance highlights the difference in the analytic goal of these two analyses: cluster visualization emphasizes on extracting as much information as possible using only the top two-dimensional components, while clustering analysis often requires a much larger number of low-dimensional components to achieve accurate performance. Subsequently, DR methods for data visualization may not fare well for cell clustering, and DR methods for cell clustering may not fare well for data visualization [20].

#### Rare cell type identification

So far, we have focused on clustering performance in terms of assigning all cells to cell types without distinguishing whether the cells belong to a rare population or a non-rare population. Identifying rare cell populations can be of significant interest in certain applications and performance of rare cell type identification may not always be in line with general clustering performance [47, 48]. Here, we examine the effectiveness of different DR methods in facilitating the detection of rare cell populations. To do so, we focused on the PBMC3k data from 10x Genomics [33]. The PBMC3k data were measured on 3,205 cells with 11 cell types. We considered CD34+ cell type (17 cells) as the rare cell population. We paired the rare cell population with either CD19+ B cells (406 cells) or CD4+/CD25 T Reg cells (198) cells to construct two data sets with different rare cell proportions. We name these two data sets PBMC3k1Rare1 and PBMC3k1Rare2, respectively. We then applied different DR methods to each data and used *F*-measure to measure the performance of rare-cell type detection following [49, 50] (details in Methods and Materials). The results are summarized in Figures S34-S35.

Overall, we found that Isomap achieves the best performance for rare cell type detection across a range of low-dimensional components in both data sets with different rare cell type proportions. As expected, the ability to detect rare cell population increases with increasing rare cell proportions. In the PBMC3k1Rare1 data, the *F*-measure by Isomap with four different number of low-dimensional components (0.5%, 1%, 2%, and 3%) are 0.74, 0.79, 0.79, and 0.79, respectively (Figure S34). The performance of Isomap is followed by ZIFA (0.74, 0.74, 0.74, and 0.74) and GLMPCA (0.74, 0.74, 0.73, and 0.74). In the PBMC3k1Rare2 data, the F-measure by Isomap with four different number of low-dimensional components (0.5%, 1%, 2%, and 3%) are 0.79, 0.79, 0.79, and 0.79, respectively (Figure S35). The performance of Isomap is also followed by ZIFA (0.74, 0.74, 0.74, and 0.74) and GLMPCA (0.74, 0.74, 0.74, and 0.74). Among the remaining methods, Poisson NMF, pCMF, scScope, and LTSA do not fare well for rare cell type detection. We note that many DR methods in conjunction with Louvain clustering method often yield an *F*-measure of zero when the rare cell type proportion is low (Figure S34C; PBMC3kRare1, 4.0% CD34+ cells) and only become reasonable with increasingly large rare cell type proportions (Figure S35C; PBMC3kRare2, 7.9% CD34+ cells). The poor performance of the Louvain clustering method for rare cell type detection is likely because its automatic way of determining cell cluster number does not fare well in the presence of uneven/un-balanced cell type proportions.

#### Stability analysis across data splits

Finally, we investigated the stability and robustness of different DR methods. To do so, we randomly split the *Kumar* data into two subsets with an equal number of cells for each cell type in the two subsets. We applied each DR method to the two subsets and measured the clustering performance in each subset separately. We repeated the procedure 10 times to capture the potential stochasticity during the data split. We visualized the clustering performance of different DR methods in the two subsets separately. Such visualization allows us to check the effectiveness of DR methods with respective to reduced sample size in the subset, as well as the stability/variability of DR methods across different split replicates (Figure S36). The results show that six DR methods, PCA, ICA, FA, ZINB-WaVE, MDS, and UMAP often achieve both accurate clustering performance and highly stable and consistent results across the subsets. The accurate and stable performance of ICA, ZINB-WaVE, MDS, and UMAP is notable even with a relatively small number of low-dimensional components. For example, with very small number of low-dimensional components, ICA, ZINB-WaVE, MDS, and UMAP achieve an average NMI value of 0.98 across the two subsets, with virtually no performance variability across data splits (Figure S36).

Overall, the results suggest that, in terms of downstream clustering analysis accuracy and stability, PCA, FA, NMF, and ICA are preferable across a range of data sets examined here. In addition, scRNAseq specific DR methods such as ZINB-WaVE, GLMPCA, and UMAP are also preferable if one is interested in extracting a small number of low-dimensional components, while generic methods such as PCA or FA are also preferred when one is interested in extracting a large number of low-dimensional components.

### Performance of DR methods for trajectory inference

We evaluated the performance of different DR methods for lineage inference applications (details in Methods and Materials). To do so, we obtained 14 publicly available scRNAseq data sets, each of which contains known lineage information (Table S2). The known lineage in all these data are linear, without bifurcation or multifurcation patterns. For each data set, we applied one DR method at a time to extract a fixed number of low-dimensional components. In the process, we varied the number of low-dimensional components from 2, 6, 14 to 20 to examine their influence for downstream analysis. With the extracted low-dimensional components, we applied two commonly used trajectory inference methods: *Slingshot* [51] and *Monocle3* [28, 52]. Slingshot is a clustering dependent trajectory inference method, which requires additional cell label information. We therefore first used either *k*-means clustering algorithm, hierarchical clustering or Louvain method to obtain cell type labels, where the number of cell types in the clustering was set to be the known truth. Afterwards, we supplied the low-dimensional components and cell type labels to the Slingshot to infer the lineage. Monocle3 is a clustering free trajectory inference method, which only requires low-dimensional components and trajectory starting state as inputs. We set the trajectory starting state as the known truth for Monocle3. Following [51], we evaluated the performance of DR methods by Kendall correlation coefficient (details in Methods and Materials) that compares the true lineage and inferred lineage obtained based on the low-dimensional components. In this comparison, we also excluded one DR method, scScope, which is not compatible with *Slingshot*. The lineage inference results for the remaining DR methods are summarized in Figures 3 and S37-S54.

**Figure 3.**
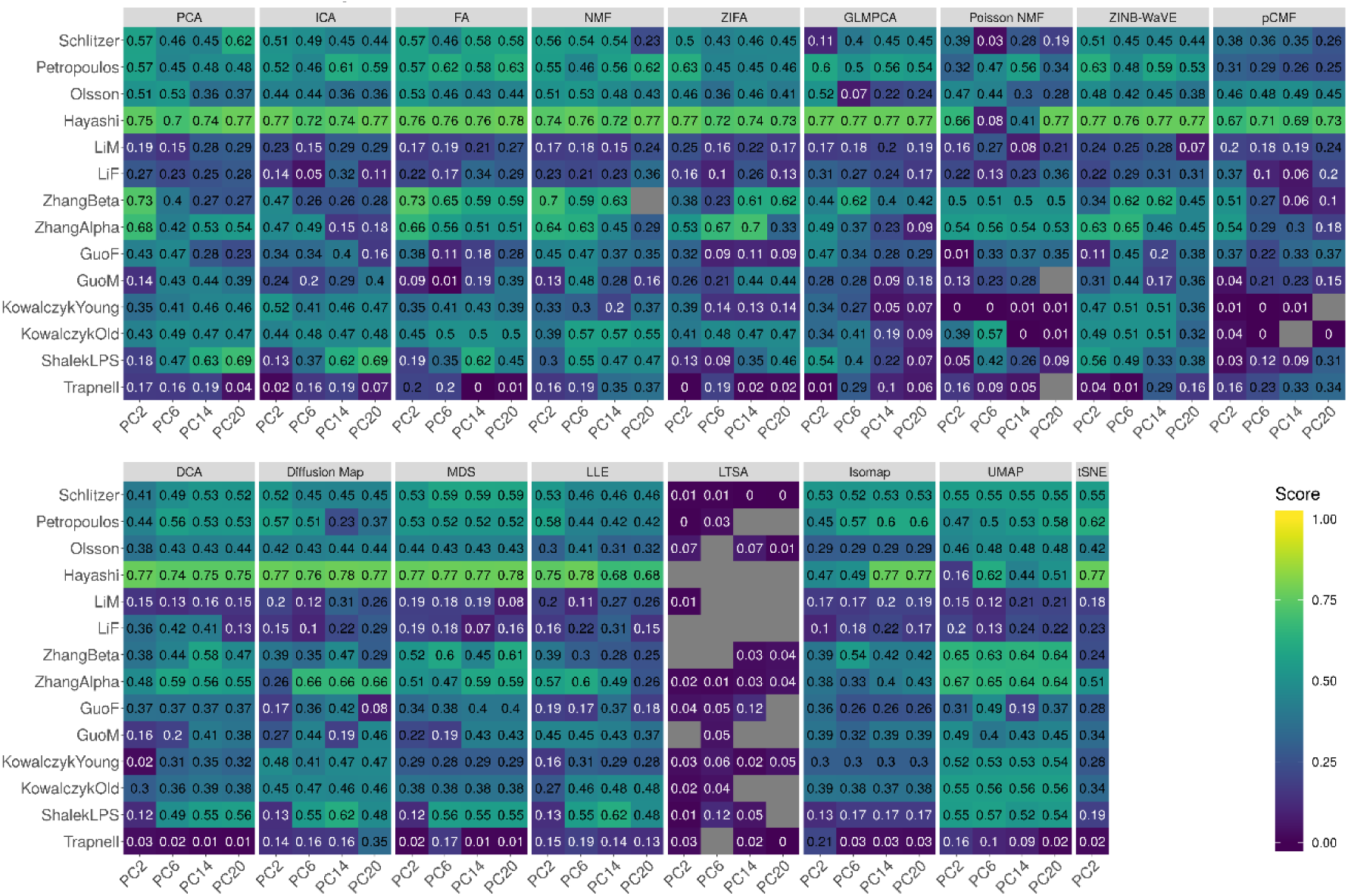
DR method performance evaluated by Kendall correlation in the downstream trajectory inference analysis. We compared 17 DR methods (columns), including factor analysis (FA), principal component analysis (PCA), independent component analysis (ICA), Diffusion Map, nonnegative matrix factorization (NMF), Poisson NMF, zero-inflated factor analysis (ZIFA), zero-inflated negative binomial based wanted variation extraction (ZINB-WaVE), probabilistic count matrix factorization (pCMF), deep count autoencoder network (DCA), generalized linear model principal component analysis (GLMPCA), multidimensional scaling (MDS), locally linear embedding (LLE), local tangent space alignment (LTSA), Isomap, uniform manifold approximation and projection (UMAP), and t-distributed stochastic neighbor embedding (tSNE). We evaluated their performance on 14 real scRNAseq data sets (rows) in terms of lineage inference accuracy. We used *Slingshot* with *k*-means as the initial step for lineage inference. The performance of each DR method is measured by Kendall correlation. For each data set, we compared four different number of low-dimensional components (2, 6, 14, and 20; four sub-columns under each column). Grey fills in the table represents missing results where *Slingshot* gave out errors when we supplied the extracted low-dimensional components from the corresponding DR method. Note that, for tSNE, we only extracted two low-dimensional components due to the limitation of the tSNE software.

#### Trajectory inference by Slingshot

We first focused on the comparison results obtained from Slingshot. Different from the clustering results where accuracy generally increases with increasing number of included low-dimensional components, the lineage tracing results from Slingshot do not show a clear increasing pattern with respect to the number of low-dimensional components, especially when we used *k*-means clustering as the initial step (Figures 3 and S39A). For example, the average Kendall correlation across all data sets and across all methods are 0.35, 0.36, 0.37 and 0.37 for increasingly large number of components, respectively. When we used hierarchical clustering algorithm as the initial step, the lineage tracing results in the case of a small number of low-dimensional components are slightly inferior as compared to the results obtained using a large number of low-dimensional components (Figures S37 and S39B). However, we do note that the lineage tracing results obtained using *k*-means are better than that obtained using hierarchical clustering as the initial step. In addition, perhaps somewhat surprisingly, the lineage tracing results obtained using Louvain clustering method are slightly better that the results obtained using *k*-means clustering (Figures S38 and S39C) – even though the clustering results from *k*-means are generally better than that from Louvain. For example, the average Kendall correlation obtained using Louvain method across all data sets and across all methods are 0.36, 0.38, 0.40 and 0.40 for increasingly large number of components, respectively. Therefore, Louvain method is recommended as the initial step for lineage inference and a small number of low-dimensional components there is often sufficient for accurate results. When conducting lineage inference based on a low number of components with Louvain method, we found that four DR methods, PCA, FA, ZINB-WaVE and UMAP, all perform well for lineage inference across varying number of low-dimension components (Figure S39C). For example, with the lowest number of components, the average Kendall correlation across data sets for PCA, FA, UMAP, and ZINB-WaVE are 0.44, 0.43, 0.40, and 0.43, respectively. Their performance is followed by ICA (0.37), ZIFA (0.36), tSNE (0.33) and Diffusion Map (0.38). While pCMF (0.26), Poisson NMF (0.26) and LTSA (0.12) do not fare well.

#### Trajectory inference by Monocle3

We next examined the comparison results based on Monocle3 (Figures S40-S41). Similar to Slingshot, we found that the lineage tracing results from Monocle3 also do not show a clear increasing pattern with respect to the number of low-dimensional components (Figure S41). For example, the average Kendall correlation across all data sets and across all methods are 0.37, 0.37, 0.38 and 0.37 for increasingly large number of components, respectively. Therefore, similar with Slingshot, we also recommend the use of a small number of low-dimensional components with Monocle3. In terms of DR method performance, we found that five DR methods, FA, MDS, GLMPCA, ZINB-WaVE and UMAP, all perform well for lineage inference. Their performance is often followed by NMF and DCA. While Poisson NMF, pCMF, LLE and LTSA do not fare well. The DR comparison results based on Monocle3 are in line with those recommendation by Monocle3 software, which uses UMAP as the default DR method [28]. In addition, the set of five top DR methods for Monocle3 are largely consistent with the set of top five DR methods for Slingshot, with only one method difference between the two (GLMPCA in place of PCA). The similarity of top DR methods based on different lineage inference methods suggests that a similar set of DR methods are likely suitable for lineage inference in general.

#### Visualization of inferred lineages

We visualized the reduced low-dimensional components from different DR methods in one trajectory data set, the ZhangBeta data. The ZhangBeta data consists of expression measurements on mouse pancreatic cells collected at seven different developmental stages. These seven different cell stages include E17.5, P0, P3, P9, P15, P18 and P60. We applied different DR methods to the data to extract the first two dimensional components. Afterwards, we performed lineage inference and visualization using Monocle3. The inferred tracking paths are shown in Figure S42. Consistent with Kendall correlation (Figure 3), all top DR methods are able to infer the correct lineage path. For example, the trajectory from GLMPCA and UMAP completely matches the truth. The trajectory inferred from FA, NMF, or ZINB-WaVE largely matches the truth with small bifurcations. In contrast, the trajectory inferred from either Poisson NMF or LTSA displays unexpected radical patterns (Figure S42), again consistent with the poor performance of these two methods in lineage inference.

#### Normalization does not influence the performance of DR methods

For DR methods that require normalized data, we further examined the influence of different data transformation approaches on their performance (Figures S43-S53). Like in the clustering comparison, we found that different transformations do not influence the performance results for most DR methods in lineage inference. For example, in Slingshot with the *k*-means clustering algorithm as the initial step, when the number of low-dimensional components is small, UMAP achieves a Kendall correlation of 0.42, 0.43 and 0.40, for log2 count transformation, log2 CPM transformation, and z-score transformation, respectively (Figures S39A, S46A, and S50A). Similar results hold for the hierarchical clustering algorithm (Figures S39B, S46B, and S50B) and Louvain method (Figures S39B, S46B, and S50B). However, some notable exceptions exist. For example, with log2 CPM transformation but not the other transformations, the performance of Diffusion Map increases with increasing number of included components when *k*-means clustering was used as the initial step: the average Kendall correlation across different low-dimensional components are 0.37, 0.42, 0.44, and 0.47, respectively (Figures S43 and S46A). As another example, with z-score transformation but not with the other transformations, FA achieves the highest performance among all DR methods across different number of low-dimensional components (Figure S50A). Similarly, in Monocle3, different transformations (log2 count transformation, log2 CPM transformation and z-score transformation) do not influence the performance of DR methods. For example, with the lowest number of low-dimensional components, UMAP achieves a Kendall correlation of 0.49, 0.47 and 0.47, for log2 count transformation, log2 CPM transformation, and z-score transformation, respectively (Figures S41, S53A and S53B).

#### Stability analysis across data splits

We also investigated the stability and robustness of different DR methods by data split in the *Hayashi* data. We applied each DR method to the two subsets and measured the lineage inference performance in the two subsets separately. We again visualize the clustering performance of different DR methods in the two subsets, separately. Such visualization allows us to check the effectiveness of DR methods with respective to reduced sample size in the subset, as well as the stability/variability of DR methods across different split replicates (Figure S54). The results show that four of the DR methods, FA, Diffusion Map, ZINB-WaVE, and MDS often achieve both accurate performance and highly stable and consistent results across the subsets. The accurate and stable performance of these is notable even with a relatively small number of low-dimensional components. For example, with very small number of low-dimensional components, FA, Diffusion Map, ZINB-WaVE and MDS achieve Kendall correlation of 0.75, 0.77, 0.77, and 0.78 averaged across the two subsets, respectively, and again with virtually no performance variability across data splits (Figure S54).

Overall, the results suggest that, in terms of downstream lineage inference accuracy and stability, the scRNAseq non-specific DR method FA, PCA, and NMF are preferable across a range of data sets examined here. The scRNAseq specific DR methods ZINB-WaVE as well as the scRNAseq non-specific DR method NMF are also preferable if one is interested in extracting a small number of low-dimensional components for lineage inference. In addition, the scRNAseq specific DR method Diffusion Map and scRNAseq non-specific DR method MDS may also be preferable if one is interested in extracting a large number of low-dimensional components for lineage inference.

### Large-scale scRNAseq data applications

Finally, we evaluated the performance of different DR methods in two large-scale scRNAseq data sets. The first data is Guo et al. [53], which consists of 12,346 single cells collected through a non-UMI based sequencing technique. Guo et al. data contains known cell cluster information and is thus used for DR method comparison based on cell clustering analysis. The second data is Cao et al. [28], which consists of approximately 2 million single cells collected through a UMI-based sequencing technique. Cao et al. data contains known lineage information and is thus used for DR method comparison based on trajectory inference. Since many DR methods are not scalable to these large-scale data sets, in addition to applying DR methods to the two data directly, we also coupled them with a recently developed sub-sampling procedure *dropClust* to make all DR methods applicable to large data [54] (details in Methods and Materials). We focus our comparison in the large-scale data using the *k*-means clustering method. We also used log2 count transformation for DR methods that require normalized data.

The comparison results when we directly applied DR methods to the Guo et al. data are shown in Figure S55. Among the methods that are directly applicable to large-scale data sets, we found that UMAP consistently outperforms the remaining DR methods across a range of low-dimensional components by a large margin. For example, the average NMI of UMAP across different number of low-dimensional components (0.5%, 1%, 2%, and 3%) are in the range between 0.60 and 0.61 (Figure S55A). In contrast, the average NMI for the other methods are in the range of 0.15-0.51. In the case of a small number of low-dimensional components, we found that the performance of both FA and NMF are reasonable and follow right after UMAP. With the subsampling procedure, we can scale all DR methods relatively easily to this large-scale data (Figure S56). As a result, several DR methods, most notably FA, can achieve similar or better performance as compared to UMAP. However, we do notice an appreciable performance loss for many DR methods through the subsampling procedure. For example, the NMI of UMAP in the sub-sampling based procedure is only 0.26, representing an approximately 56% performance loss compared to the direct application of UMAP without sub-sampling (Figure S56 vs Figure S55). Therefore, we caution the use of sub-sampling procedure and recommend users to careful examine the performance of DR methods before and after sub-sampling to decide whether sub-sampling procedure is acceptable for their own applications.

For lineage inference in the Cao et al. data, due to computational constraint, we randomly obtained 10,000 cells from each of the five different developmental stages (i.e., E9.5, E10.5, E11.5, E12.5 and E13.5) and applied different DR methods to analyze the final set of 50,000 cells. Because most DR methods are not scalable even to these 50,000 cells, we only examined the performance of DR methods when paired with the sub-sampling procedure (Figure S57). With the small number of low-dimensional components, three DR methods, GLMPCA, DCA and Isomap, all achieve better performance than the other DR methods. For example, with the lowest number of low-dimensional components, the average absolute Kendall correlation of GLMPCA, DCA and Isomap are 0.13, 0.28, and 0.17, respectively. In contrast, the average absolute Kendall correlation of the other DR methods are in the range of 0.01-0.12. With a higher number of low-dimensional components, Isomap and UMAP show better performance. For example, with 3% low-dimensional components, the average absolute Kendall correlation of Isomap and UMAP increase to 0.17 and 0.30, respectively. Their performance is followed by Diffusion Map (0.15), ZINB-WaVE (0.14), and LLE (0.12); while the remaining methods are in the range of 0.04-0.07.

### Computation time

We recorded and compared computing time for different DR methods on simulated data sets. Here, we also examined how computation time for different DR methods varies with respect to the number of low-dimensional components extracted (Figure 4A) as well as with respect to the number of cells contained in the data (Figure 4B). Overall, the computational cost of three methods, ZINB-WaVE, ZIFA, and pCMF, is substantially heavier than the remaining methods. Their computation time increase substantially with both increasingly large number of low-dimensional components and increasingly large number of cells in the data. Specifically, when the sample size equals 500 and the desired number of low dimensional components equals 22, the computing time for ZINB-WaVE, ZIFA, and pCMF to analyze 10,000 genes are 2.15, 1.33, and 1.95 hours, respectively (Figure 4A). When the sample size increases to 10,000, the computing time for ZINB-WaVE, ZIFA, and pCMF increases to 12.49, 20.50, and 15.95 hours, respectively (Figure 4B). Similarly, when the number of low-dimensional components increases to 52, the computing time for ZINB-WaVE, ZIFA, and pCMF increases to 4.56, 4.27, and 4.62 hours, respectively. Besides these three methods, the computing cost of ICA, GLMPCA, and Poisson NMF can also increase noticeably with increasingly large number of low-dimensional components. The computing cost of ICA, but to a lesser extent of GLMPCA, LLE, LTSA and Poisson NMF, also increases substantially with increasingly large number of cells. In contrast, PCA, FA, Diffusion Map, UMAP, and the two deep learning-based methods (DCA and scScope) are computationally efficient. In particular, the computation time for these six methods are stable and do not show substantial dependence on the sample size or the number of low-dimensional components. Certainly, we expect that the computation time of all DR methods will further increase as the sample size of the scRNAseq data sets increases in magnitude. Overall, in terms of computing time, PCA, FA, Diffusion Map, UMAP, DCA, and scScope are preferable.

**Figure 4.**
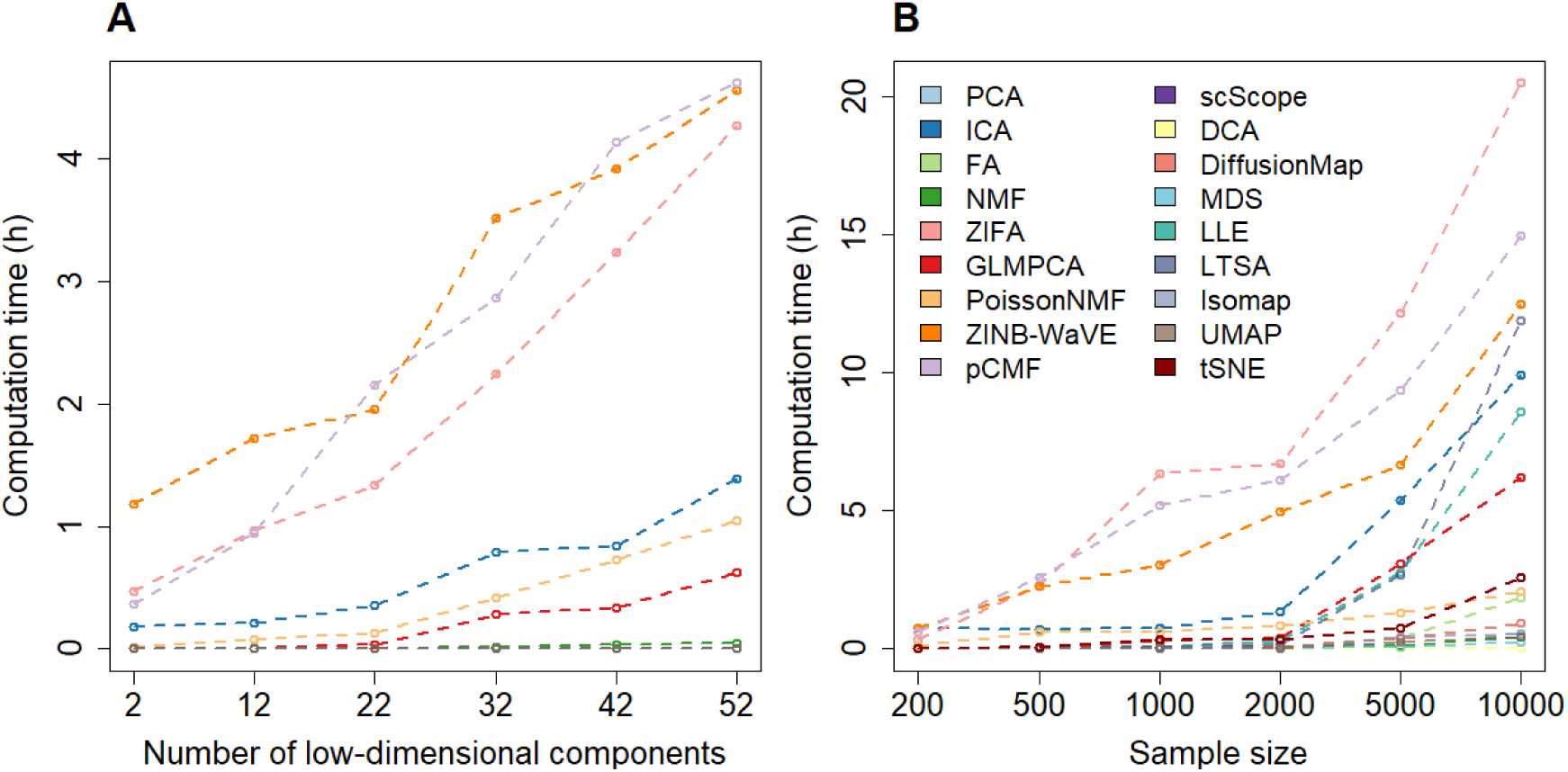
The computation time (in hours) for different DR methods. We recorded computing time for 18 DR methods on simulated data sets with varying number of low-dimensional components and varying number of sample sizes. Compared DR methods include: factor analysis (FA; light green), principal component analysis (PCA; light blue), independent component analysis (ICA; blue), Diffusion Map (pink), nonnegative matrix factorization (NMF; green), Poisson NMF(light orange), zero-inflated factor analysis (ZIFA; light pink), zero-inflated negative binomial based wanted variation extraction (ZINB-WaVE; orange), probabilistic count matrix factorization (pCMF; light purple), deep count autoencoder network (DCA; yellow), scScope (purple), generalized linear model principal component analysis (GLMPCA; red), multidimensional scaling (MDS; cyan), locally linear embedding (LLE; blue green), local tangent space alignment (LTSA; teal blue), Isomap (grey), uniform manifold approximation and projection (UMAP; brown), and t-distributed stochastic neighbor embedding (tSNE; dark red). (**A**) Computation time for different DR methods (y-axis) changes with respect to an increasing number of low-dimensional components (x-axis). The number of cells is fixed to be 500 and the number of genes is fixed to be 10,000 in this set of simulations. Three methods (ZINB-WaVE, pCMF, and ZIFA) become noticeably computationally more expensive than the remaining methods with increasing number of low-dimensional components. (**B**) Computation time for different DR methods (y-axis) changes with respect to an increasing sample size (i.e. the number of cells) in the data. Computing time is recorded on a single thread of an Intel Xeon E5-2683 2.00 GHz processor. The number of low-dimensional components is fixed to be 22 in this set of simulations for most methods, except for tSNE which used two low-dimensional components due to the limitation of the tSNE software. Note that some methods are implemented with parallelization capability (e.g. ZINB-WaVE and pCMF) though we tested them on a single thread for fair comparison across methods. Note that PCA is similar to ICA in (**A**) and scScope is similar to several other efficient methods in (**B**); thus their lines may appear to be missing. Overall, three methods (ZIFA, pCMF, and ZINB-WaVE) become noticeably computationally more expensive than the remaining methods with increasing number of cells in the data.

### Practical guidelines

In summary, our comparison analysis shows that different DR methods can have different merits for different tasks. Subsequently, it is not straightforward to identify a single DR method that strives the best in all data sets and for all downstream analyses. Instead, we provide a relatively comprehensive practical guideline for choosing DR methods in scRNAseq analysis in Figure 5. Our guideline is based on the accuracy and effectiveness of DR methods in terms of the downstream analysis, the robustness and stability of DR methods in terms of replicability and consistency across data splits, as well as their performance in large-scale data applications, data visualization, as well as computational scalability for large scRNAseq data sets. Briefly, for cell clustering analysis, PCA, ICA, FA, NMF, and ZINB-WaVE are recommended for small data where computation is not a concern. PCA, ICA, FA, NMF are also recommended for large data where computation is a concern. For lineage inference analysis, FA, PCA, NMF, UMAP and ZINB-WaVE are all recommended for small data. A subset of these methods, FA, PCA, NMF and UMAP are also recommended for large scRNAseq data. In addition, for very large scRNAseq data sets (e.g. >100,000 samples), DCA and UMAP perhaps are the only feasible approach for both downstream analyses with UMAP being the preferred choice. We also recognize that PCA, ICA, FA and NMF can be useful options in very large data sets when paired with a sub-sampling procedure [54], though care need to be taken to examine the effectiveness of the sub-sampling procedure itself. Finally, besides these general recommendations, we note that some methods have additional features that are desirable for practitioners. For example, ZINB-WaVE can include sample-level and gene-level covariates, thus allowing us to easily control for batch effects or size factors. We provide our detailed recommendations in Figure 5.

**Figure 5.**
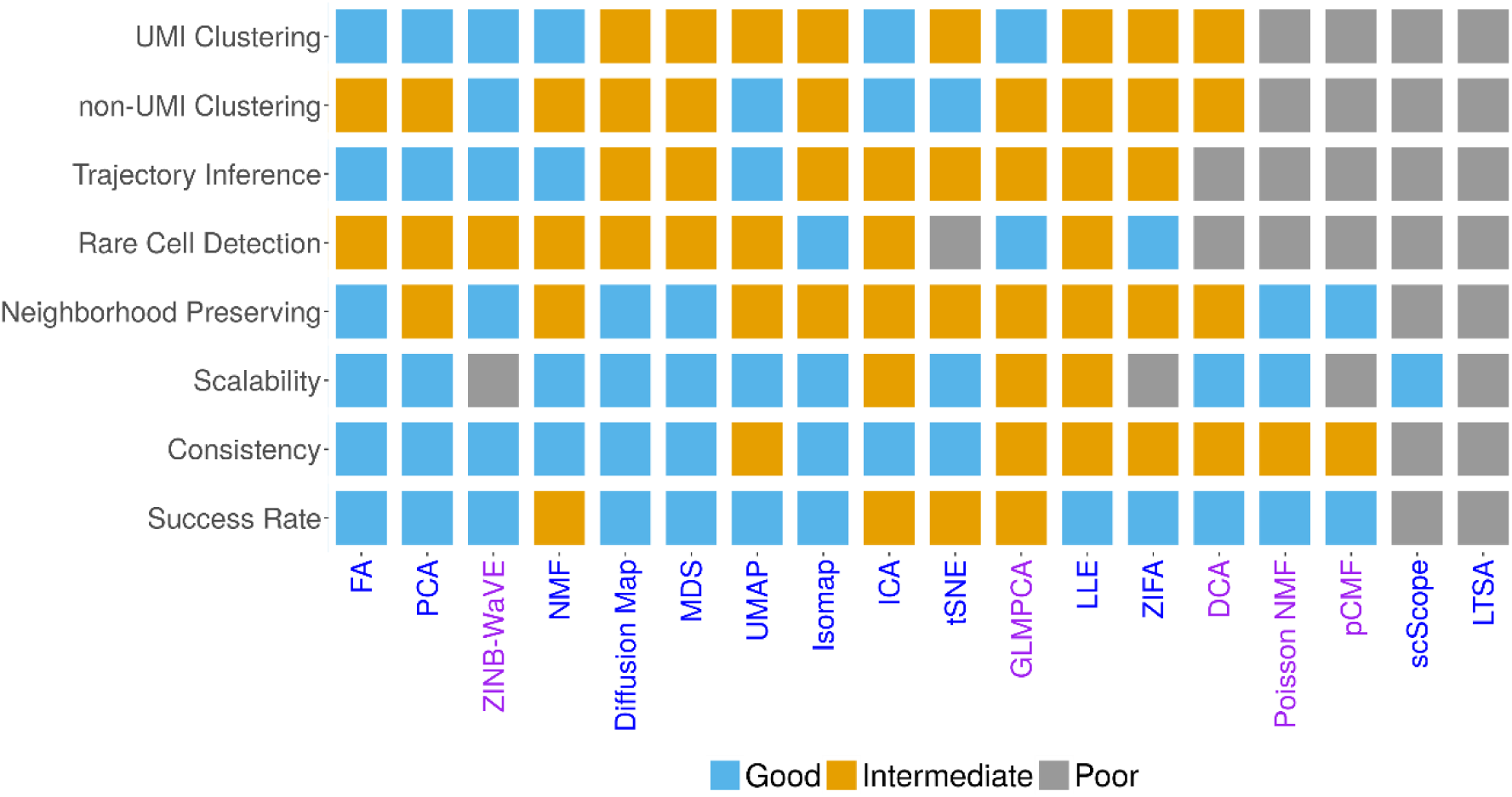
Practical guideline for choosing DR methods in scRNAseq analysis. Compared DR methods include: factor analysis (FA), principal component analysis (PCA), independent component analysis (ICA), Diffusion Map, nonnegative matrix factorization (NMF), Poisson NMF, zero-inflated factor analysis (ZIFA), zero-inflated negative binomial based wanted variation extraction (ZINB-WaVE), probabilistic count matrix factorization (pCMF), deep count autoencoder network (DCA), scScope, generalized linear model principal component analysis (GLMPCA), multidimensional scaling (MDS), locally linear embedding (LLE), local tangent space alignment (LTSA), Isomap, uniform manifold approximation and projection (UMAP), and t-distributed stochastic neighbor embedding (tSNE). The count-based methods are colored in purple while non count-based methods are colored in blue. Methods are ranked by their average performance across the criteria from left to right. The performance is colored and numerically coded: good performance = 2 (sky-blue), intermediate performance = 1 (orange), and poor performance = 0 (grey).

## DISCUSSION

We have presented a comprehensive comparison of different dimensionality reduction methods for scRNAseq analysis. We hope the summary of these state-of-the-art DR methods, the detailed comparison results, and the recommendations and guidelines for choosing DR methods can assist researchers in the analysis of their own scRNAseq data.

In the present study, we have primarily focused on three clustering methods (*k*-means, hierarchical clustering, and Louvain method) to evaluate the performance of different DR methods for downstream clustering analysis. We have also primarily focused on two lineage inference methods (Slingshot and Monocle3) to evaluate the performance of different DR methods for downstream lineage inference. In our analysis, we found that the performance of DR methods measured based on different clustering methods are often consistent with each other. Similarly, the performance of DR methods measured based on different lineage inference methods are also consistent with each other. However, it is possible that some DR methods may work well with certain clustering approaches and/or with certain lineage inference approaches. Subsequently, future comparative analysis using other clustering methods and other lineage inference methods as comparison criteria may have added benefits. In addition, besides cell clustering and trajectory inference, we note that DR methods are also used for many other analytic tasks in scRNAseq studies. For example, factor models for DR is an important modeling part for multiple scRNAseq data sets alignment [16], for integrative analysis of multiple omics data sets [55, 56], as well as for deconvoluting bulk RNAseq data using cell type specific gene expression measurements from scRNAseq [57, 58]. In addition, cell classification in scRNAseq also relies on a low-dimensional structure inferred from original scRNAseq through DR [59, 60]. Therefore, the comparative results obtained from the present study can provide important insights into these different scRNAseq analytic tasks. In addition, investigating the performance of DR methods in these different scRNAseq downstream analyses is an important future research direction.

We mostly focused on evaluating feature extraction methods for DR. Another important category of DR method is the feature selection method, which aims to select a subset of features/genes directly from the original feature space. The feature section methods rely on different criteria to select important genes and are also commonly used in the preprocessing step of scRNAseq data analysis [61].

For example, M3Drop relies on dropout events in scRNAseq data to identify informative genes [62]. Seurat uses gene expression variance to select highly variable genes [16]. Evaluating the benefits of different methods and criteria for selecting informative genes for different downstream tasks is another important future direction.

With the advance of scRNAseq technologies and with the increase collaborations across scientific groups, new consortium projects such as the Human Cell Atlas (HCA) will generate scRNAseq data sets that contain millions of cells [34]. The large data at this scale poses critical computational and statistical challenges to many current DR methods. Many existing DR methods, in particular those that require the computation and memory storage of a covariance or distance matrix among cells, will no longer be applicable there. We have examined a particular sub-sampling strategy to scale all DR methods to large data sets. However, while the sub-sampling strategy is computationally efficient, it unfortunately reduces the performance of many DR methods by a substantial margin. Therefore, new algorithmic innovations and new efficient computational approximations will likely be needed to effectively scale many of the existing DR methods to millions of cells.

## METHODS AND MATERIALS

### ScRNAseq data sets

We obtained a total of 30 scRNAseq data sets from public domains for benchmarking DR methods. All data sets were retrieved from the Gene Expression Omnibus (GEO) database (https://www.ncbi.nlm.nih.gov/geo/) or the 10X genomics website (https://support.10xgenomics.com/single-cell-gene-expression/datasets). These data sets cover a wide variety of sequencing techniques that include Smart-Seq2 (8 data sets), 10X genomics (6 data sets), Smart-Seq (5 data sets), inDrop (1 data set), RamDA-seq (1 data set), sci-RNA-seq3 (1 data set), SMARTer (5 data sets) and others (3 data sets). In addition, these data cover a range of sample sizes from a couple hundred cells to tens of thousands of cells measured in either human (19 data sets) or mouse (11 data sets). In each data set, we evaluated the effectiveness of different DR methods for one of the two important downstream analysis tasks: cell clustering and lineage inference. In particular, 15 data sets were used for cell clustering evaluation while another 15 data sets were used for lineage inference evaluation. For cell clustering, we followed the same criteria listed in [12, 41] to select these datasets. In particular, the selected data sets need to contain true cell clustering information which is to be treated as the ground truth in the comparative analysis. In our case, 11 of the 15 data sets were obtained by mixing cells from different cell types either pre-determined by fluorescence activated cell sorting (FACS) or cultured on different conditions. Therefore, these 11 studies contain the *true* cell type labels for all cells. The remaining 4 data sets contain cell labels that were determined in the original study and we simply treated them as truth though we do acknowledge that such “true” clustering information may not be accurate. For lineage inference, we followed the same criteria listed in [14] to select these datasets. In particular, the selected data sets need to contain true linear lineage information which is to be treated as the ground truth in the comparative analysis. In our case, 4 of the 15 data sets were obtained by mixing cells from different cell types pre-determined by FACS. These different cell types are at different developmental stages of a single linear lineage; thus these 4 studies contain the *true* lineage information for all cells. The remaining 11 data sets contain cells that were collected at multiple time points during the development process. For these data, we simply treated cells at these different time points as part of a single linear lineage, though we do acknowledge that different cells collected at the same time point may represent different developmental trajectories from an early time point if the cells at the early time are heterogeneous. In either case, the true lineage in all these 15 data sets are treated as linear, without any bifurcation or multifurcation patterns.

A detailed list of the selected scRNAseq datasets with corresponding data features is provided in Tables S1-S2. In each of the above 30 data sets, we removed genes that are expressed in less than five cells. For methods modeling normalized data, we transformed the raw counts data into continuous data with the *normalize* function implemented in *scater* (R package v1.12.0). We then applied log2 transformation on the normalized counts by adding one to avoid log transforming zero values. We simply term this normalization as log2 count transformation, though we do acknowledge that such transformation does take into account of cell size factor etc. through the *scater* software. In addition to log2 count transformation, we also explored the utility of two additional data transformation: log2 CPM transformation and z-score transformation. In the log2 CPM transformation, we first computed counts per million reads (CPM) and then performed log2 transformation on the resulted CPM value by adding a constant of one to avoid log transformation of zero quantities. In the z-score transformation, for each gene in turn, we standardized CPM values to achieve a mean of zero and variance of one across cells using *Seurat* package (v2.3).

Besides the above 30 real scRNAseq data sets, we also simulated 2 additional scRNAseq data sets for cell clustering evaluation. In the simulations, we used all 94 cells from one cell type (*v6.5 mouse 2i+LIF*) in the Kumar data as input. We simulated scRNAseq data with 500 cells and a known number of cell types, which were set to be either 4 or 8, using the *Splatter* package v1.2.0. All parameters used in the *Splatter* (e.g., mean rate, shape, dropout rate, etc.) were set to be approximately those estimated from the real data. In the case of 4 cell types, we set the group parameter in *Splatter* as 4. We set the percentage of cells in each group as 0.1, 0.15, 0.5 and 0.25, respectively. We set the proportion of the differentially expressed genes in each group as 0.02, 0.03, 0.05 and 0.1, respectively. In the case of 8 cell types, we set group/cell type parameter as 8. We set the percentage of cells in each group as 0.12, 0.08, 0.1, 0.05, 0.3, 0.1, 0.2 and 0.05, respectively. We set the proportion of the differentially expressed genes in each group as 0.03, 0.03, 0.03, 0.1, 0.05, 0.07, 0.08, and 0.1, respectively.

### Compared dimensionality reduction methods

DR methods aim to transform an originally high-dimensional feature space into a low-dimensional representation with a much-reduced number of components. These components are in the form of a linear or non-linear combination of the original features (known as feature extraction DR methods) [63] and in the extreme case are themselves a subset of the original features (known as feature selection DR methods) [64]. In the present study, we have collected and compiled a list of 18 popular and widely used DR methods in the field of scRNAseq analysis. These DR methods include factor analysis (FA; R package *psych*, v1.8.12), principal component analysis (PCA; R package *stats*, v3.6.0), independent component analysis (ICA; R package *ica*, v1.0.2), Diffusion Map (Diffusion Map; R package *destiny*, v2.14.0), nonnegative matrix factorization (NMF; R package NNLM, v1.0.0), Kullback-Leibler divergence-based NMF (Poisson NMF; R package NNLM, v1.0.0), zero-inflated factor analysis (ZIFA; Python package *ZIFA*), zero-inflated negative binomial based wanted variation extraction (ZINB-WaVE; R package *zinbwave*, v1.6.0), probabilistic count matrix factorization (pCMF; R package pCMF, v1.0.0), deep count autoencoder network (DCA; Python package *dca*), a scalable deep-learning-based approach (scScope; Python package *scscope*), generalized linear model principal component analysis (GLMPCA; R package on github), multidimensional scaling (MDS; Rdimtools R package v.0.4.2), locally linear embedding (LLE; Rdimtools R packge v.0.4.2), local tangent space alignment (LTSA; Rdimtools R package v.0.4.2), Isomap (Rdimtools R package v.0.4.2), t-distributed stochastic neighbor embedding (tSNE; FIt-SNE, *fftRtnse* R function), and uniform manifold approximation and projection (UMAP; Python package). One of these methods, tSNE, can only extract a maximum of two or three low-dimensional components [42–44]. Therefore, we only included tSNE results based on two low-dimensional components extracted from the recently developed fast *FIt-SNE* R package [44] in all figures. An overview of these 18 DR methods with their corresponding modeling characteristics is provided in Table 1.

### Assess the performance of dimensionality reduction methods

We first evaluated the performance of DR methods by neighborhood preserving that aims to access whether the reduced dimensional space resembles the original gene expression matrix. To do so, we first identified the *k*-nearest neighbors for each single cell in the original space (denoted as a set A) and in the reduced space (denoted as a set B). We set *k* = 10, 20, or 30 in our study. We then computed the Jaccard index (JI) [45] to measure the neighborhood similarity between the original space and the reduced space: 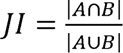, where |·| denotes the cardinality of a set. We finally obtained the averaged Jaccard index (AJI) across all cells to serve as the measurement for neighborhood preserving. We note, however, that neighborhood preserving is primarily used to measure the effectiveness of pure dimensionality reduction in terms of preserving the original space and may not be relevant for single cell analytic tasks that are the main focus of the present study: a DR method that preserve the original gene expression matrix effectively may not be effective in extracting useful biological information from the expression matrix that are essential for key downstream single cell applications. Preserving the original gene expression matrix is rarely the purpose of DR methods for single cell applications: indeed, the original gene expression matrix (which is the best-preserved matrix of itself) is rarely, if ever, used directly in any downstream single cell applications including cell clustering and lineage inference, even though it is computationally easy to do so.

Therefore, more importantly, we also evaluated the performance of DR methods by evaluating how effective the low-dimensional components extracted from DR methods are for downstream single cell analysis. We evaluated either of the two commonly applied downstream analysis, clustering analysis and lineage reconstruction analysis, in the 32 data sets described above. In the analysis, we varied the number of low-dimensional components extracted from these DR methods. Specifically, for cell clustering data sets, in a data with less than or equal to 300 cells, we varied the number of low dimensional components to be either 2, 6, 14, or 20. In a data with more than 300 cells, we varied the number of low dimensional components to be either 0.5%, 1%, 2%, or 3% of the total number of cells. For lineage inference data sets, we varied the number of low dimensional components to be either 2, 6, 14, or 20 for all data sets, since common lineage inference methods prefer a relatively small number of components.

For clustering analysis, after DR with these DR methods, we used three different clustering methods, the hierarchical clustering (R function *hclust*; stats v3.5.3), *k*-means clustering (R function *kmeans*; stats v3.6.0), or Louvain method (R function *clusterCells*; monocle v2.12.0) to perform clustering on the reduced feature space. The *k*-means clustering is a key ingredient of commonly applied scRNAseq clustering methods such as SC3 [18] and Waterfall [25]. The hierarchical clustering is a key ingredient of commonly applied scRNAseq clustering methods such as CIDR [17] and CHETAH [65]. The Louvain method is also a commonly used clustering method for common single cell analysis software such as Seurat [16] and Monocle [27, 66]. In all these clustering methods, we set the number of clusters *k* to be the known number of cell types in the data. We compared the cell clusters inferred using the low dimensional components to the true cell cluster and evaluated clustering accuracy by two criteria: the adjusted rand index (ARI) [67] and the normalized mutual information (NMI) [68]. The ARI and NMI are defined as:

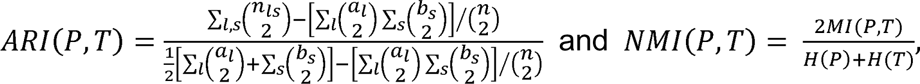

where *P* = (*p*_1_, *p*_2_, …, *p_n_*)*^T^* denotes the inferred cell type cluster labels from clustering analysis while, *T* = (*t*_1_, *t*_2_, *…, t_n_*)*^T^* denotes the known true cell type labels for *n* samples in the data; *l* and *s* enumerate the clusters, with *l* = 1, …, *r* and *s* = 1, …, *k* where *r*. and *k* are the number of inferred cell type clusters and the number of true cell type clusters, respectively; *n_ls_* = Σ*_ij_ I*(*p_i_* = *l*)*I*(*t_j_* = *s*) is the number of times where the *i*’th cell belongs to the cluster *l* in the inferred cluster labeling and *j*’th cell belongs to the cluster *s* in the true cluster labeling; note that *n_ls_* is an entry of contingency table which effectively measures the number of cells that are in common between *P* and *T*, with *I*(·)·being an indicator function; *a_l_* = Σ*_s_ n_ls_* is the sum of the *s*th column of the contingency table; and *b_s_* = Σ*_l_ n_ls_* is the sum of the Ith row of the contingency table;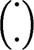 denotes a binomial coefficient; 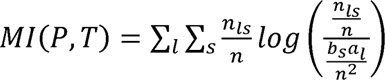 is the mutual information between two cluster labels; 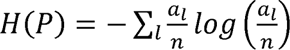 is the entropy function for inferred cell type labeling; and 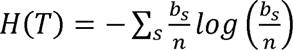 is the entropy function for true cell type labeling. We used the *compare* function in the *igraph* R package (v1.0.0) to compute both ARI and NMI criteria. For rare cell type identification, we used the -measure, that is commonly used for quantifying rare cell type identification performance [49, 50]. The *F*-measure is the harmonic mean of the clustering’s precision and recall, and is formulated as:

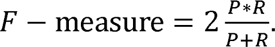

where *P* represents the precision for identifying the rare cluster, with 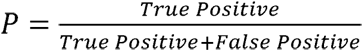; while *R* represents the recall for identifying the rare cluster, with 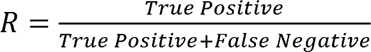. For each data set, we repeated the above procedure five times and report the averaged results to avoid the influence of the stochasticity embedded in some DR methods and/or the clustering algorithm.

While it is straightforward to apply different DR methods to most scRNAseq data sets, we found that many DR methods are not computationally scalable and cannot be directly applied for clustering analysis in two large-scale scRNAseq data sets we examined in the present study. For these non-scalable DR methods, we made use of a recently developed subsampling procedure described in *dropClust* to scale them to large data [54]. In particular, we first applied dropClust to the original large-scale data to infer rare cell populations. We then created a small data by combining all cells in the rare cell populations along with a subset set of cells in the remaining cell populations. The subset of cells in the non-rare populations are obtained through subsampling using the structure preserving sampling procedure (details in [54]). Afterwards, we applied different DR methods to the small data and performed clustering analysis there. The cells in the small data are then directly assigned with their clustering label after clustering analysis. For each cell that is not in the small data, we computed the Pearson correlation between the cell and each of the cluster centers inferred in the small data. We assigned the cell to the cluster with the closest cluster center in the small data as the cluster assignment.

For trajectory inference, after DR with these DR methods, we used *Slingshot* [51] (R package, v1.2.0) and Monocle3 [28] (R package, v0.1.2). The *Slingshot* software is the recommended lineage inference method based on a recent comparative study [14]. Monocle3 is one of the most recent lineage inference methods. Slingshot takes two input data: the low-dimensional components extracted from DR methods and a vector of cluster labels predicted by clustering algorithms. Monocle3 also takes two input data: the low-dimensional components extracted by DR methods and starting state which is to the beginning of the lineage. For the cluster labels, we used either *k*-means, hierarchical clustering algorithm or Louvain method on the extracted low-dimensional components to obtain cluster labels. For the starting state, we supplied with the true beginning state of the lineage in the data. After obtaining the two types of input through the *slingshot* function, we used the *getLineages* function to fit a minimum spanning tree (MST) to identify lineage. The final output from *Slingshot* is an object of class *SlingshotDataSet* that contains the inferred lineage information. We follow the original *Slingshot* paper [51] to evaluate the accuracy of the inferred lineage using the Kendall rank correlation coefficient. To do so, for each data, we first ranked genes based on their position on the true lineage. We ordered all *m* genes based on this rank order and denoted the corresponding rank in ascending order for these genes as {*x*_1_, …, *x_m_*}, where *x_i_* ≤ *x_i_*_+1_. Note that the true lineage is linear without any bifurcation or multifurcation patterns, while the inferred lineage may contain multiple ending points in addition to the single starting point. Therefore, for each inferred lineage, we examined one trajectory at a time, where each trajectory consists of the starting point and one of the ending points. In each trajectory, we ranked genes in order based on their position in the trajectory. We denote the corresponding rank order in the inferred trajectory for all *m* genes as {*y*_1_, …, *y_m_*}, where we set *y_i_* as missing if *l*’th gene is not included in the inferred trajectory. For each pair of non-missing genes, we labeled the gene pair (*i*, *j*) as a concordant pair if their relative rank in the inferred lineage are consistent with their relative rank in the true lineage; that is, either (*x_i_* ≥ *x_j_* & *y_i_* ≥ *y_j_*) or (*x_i_* < *x_j_* & *y_i_* < *y_j_*). Otherwise, we labeled the gene pair (*i*, *j*) as discordant. We denoted *C* as the number of concordant pairs, *D* as the number of discordant pairs, and *U* as the total number of non-missing genes. The Kendell correlation coefficient is then computed as

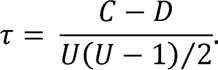

Afterwards, we obtained the maximum absolute over all these trajectories as the final Kendall correlation score to evaluate the similarity between the inferred lineage and the true lineage. For each data set, we repeated the above procedure five times and report the averaged results to avoid the influence of the stochasticity embedded in some DR methods and/or the lineage inference algorithm. For the large-scale data application to Cao et al., we also applied the sub-sampling approach dropClust to scale different DR methods for lineage inference.

We investigated the stability and robustness of different DR methods in both cell clustering and lineage inference applications through data splitting. Here, we focused on two representative scRNAseq data sets, the *Kumar* data set for cell clustering and the *Hayashi* data set for lineage inference. For each data, we randomly split the data into two subsets with an equal number of cells in each cell type in the two subsets. We repeated the split procedure 10 times to capture the potential stochasticity during the data split. In each split replicate, we applied different DR methods to analyze each subset separately. We used *k*-means clustering algorithm to infer the clustering labels in each subset. We used NMI to measure cell clustering accuracy and used Kendall correlation to measure lineage inference accuracy.

Finally, to summarize the performance of the evaluated DR methods across the range of criteria in Figure 5, we consider either “good”, “intermediate” or “poor” to categorize the DR methods for each criterion. For UMI and non-UMI in cell clustering, we evaluated the performance of different DR methods based on 0.5% low-dimensional components in Figures S31A and S31B: average NMI ≥ 0.73 (good); ≤ 0.64 average NMI < 0.73 (intermediate); average NMI < 0.64 (poor). For Trajectory Inference, we evaluated the performance of different DR methods based on 2 low-dimensional components in Figure S39A: average Kendall ≥ 0.41 (good); 0.35 ≤ average Kendall < 0.41 (intermediate); average Kendall < 0.35 (poor). For Rare Cell Detection, we evaluated the performance of different DR methods based on 0.5% low-dimensional components in Figure S35A: F-measure ≥ 0.74 (good); 0.69 ≤ F-measure < 0.74 (intermediate); F-measure < 0.69 (poor). For Neighborhood Preserving, we evaluated the performance of different DR methods based on 0.5% low-dimensional components in Figure S7A: average Jaccard index ≥ 0.15 (good); 0.12 ≤ average Jaccard index < 0.15 (intermediate); average Jaccard index < 0.12 (poor). For Scalability, we evaluated the performance of different DR methods when sample size is 10,000 in Figure 4B: computation time ≤ 0.25h (good); 0.25h ≤ computation time < 10 (intermediate); computation time ≥ 10h (poor). For Consistency, we evaluated the performance of different DR methods based on the absolute mean value of the difference of average NMI between two splits from Figures S36 and S54: difference of average NMI ≤ 0.005 (good); 0.005 ≤ difference of average NMI < 0.01 (intermediate); difference of average NMI ≥ 0.01 (poor). For Success Rate, since both scScope and LTSA do not work for most trajectory inference data sets, we set as poor; NMF, ICA, tSNE, and GLMPCA do not work for some of data sets, we set as intermediate; the rest of DR methods are all good.

## ACKNOWLEDGEMENTS

We would like to thank Angelo Duò for the part of single cell RNAseq data sets.

## AVAILABILITY OF DATA AND MATERIALS

All source code and data sets used in our experiments have been deposited at www.xzlab.org/reproduce.html or https://github.com/xzhoulab/DRComparison.

## AUTHORS’ CONTRIBUTIONS

XZ conceived the idea and provided funding support. SS and XZ designed the experiments. SS adapted software, performed simulations and analyzed real data. JQ and YM collected data sets and helped interpreting results. SS and XZ wrote the manuscript with input from all other authors.

## ETHICS APPROVAL AND CONSENT TO PARTICIPATE

No ethnical approval was required for this study. All utilized public data sets were generated by other organizations that obtained ethical approval.

## CONSENT FOR PUBLICATION

Not applicable.

