## supplemental text for "Accuracy, Robustness and Scalability of Dimensionality Reduction Methods for Single Cell RNAseq Analysis"

**Supplementary Figure 1. Popularity of different DR methods as measured by their use in existing scRNAseq cell clustering tools or lineage inference tools**. We counted the number of DR methods that are used in each of the 32 scRNAseq cell clustering methods (**A**) or in each of the 46 scRNAseq lineage inference methods (**B**). We used *geom*_*treemap* functions in R package to draw both panels, where the font size of each DR method represents its popularity, which is measured by the frequency of DR method used in these scRNAseq tools.

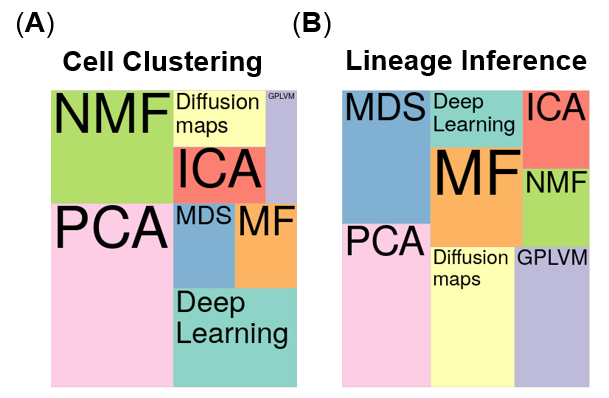

**Supplementary Figure 2. DR method performance evaluated by Jaccard index on cell clustering data sets with 10 neighborhood cells.** We compared 18 DR methods (columns), including factor analysis (FA), principal component analysis (PCA), independent component analysis (ICA), Diffusion Map, nonnegative matrix factorization (NMF), Poisson NMF, zero-inflated factor analysis (ZIFA), zero-inflated negative binomial based wanted variation extraction (ZINB-WaVE), probabilistic count matrix factorization (pCMF), deep count autoencoder network (DCA), scScope, generalized linear model principal component analysis (GLMPCA), multidimensional scaling (MDS), locally linear embedding (LLE), local tangent space alignment (LTSA), Isomap, uniform manifold approximation and projection (UMAP), and t-distributed stochastic neighbor embedding (tSNE). We evaluated their performance on 14 real scRNAseq data sets (UMI-based data are colored as blue while non-UMI based data are colored as purple) and 2 simulated data sets (rows). The simulated data based on Kumar data is labeled with #. We used log2 count transformation for the subset of DR methods that use normalized data. For each data set, we compared the four different number of low-dimensional data. The four numbers we used equal to 0.5%, 1%, 2%, and 3% of the total number of cells in big data and equal to 2, 6, 14, and 20 in small data (which are labeled with *). No results for ICA are shown in the Pancreatic data (grey fills) because ICA cannot handle the large number of features in the data. Note that, for tSNE, we only extracted two low-dimensional components due to the limitation of the tSNE software.

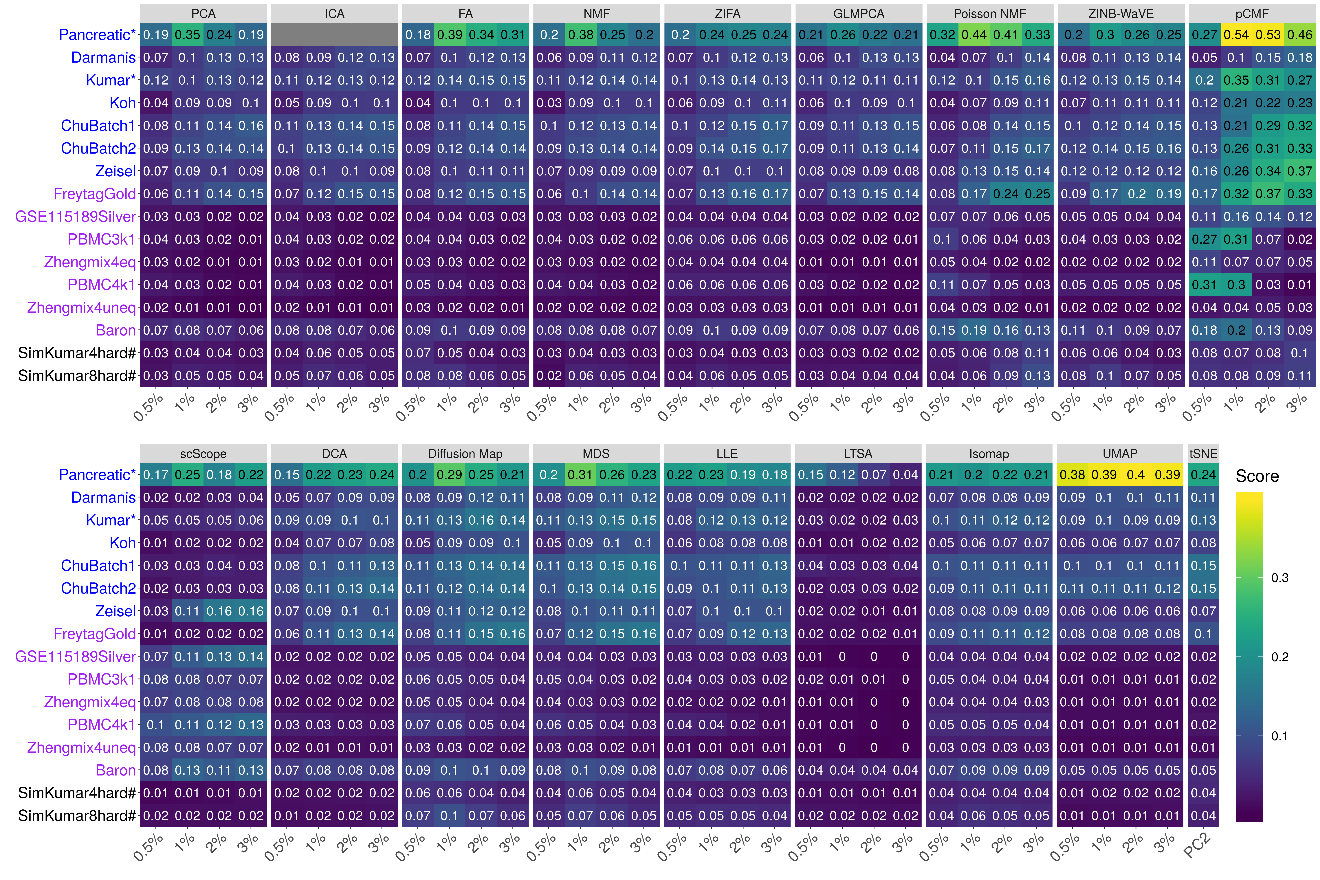

**Supplementary Figure 3. Bar plots show the average Jaccard index of different DR methods with 10 neighborhood cells based on cell clustering data.** The performance is measured by average Jaccard index across 16 data sets. We compared 18 DR methods, including factor analysis (FA; light green), principal component analysis (PCA; light blue), independent component analysis (ICA; blue), Diffusion Map (pink), nonnegative matrix factorization (NMF; green), Poisson NMF(light orange), zero-inflated factor analysis (ZIFA; light pink), zero-inflated negative binomial based wanted variation extraction (ZINB-WaVE; orange), probabilistic count matrix factorization (pCMF; light purple), deep count autoencoder network (DCA; yellow), scScope (purple), generalized linear model principal component analysis (GLMPCA; red), multidimensional scaling (MDS; cyan), locally linear embedding (LLE; blue green), local tangent space alignment (LTSA; teal blue), Isomap (grey), uniform manifold approximation and projection (UMAP; brown), and t-distributed stochastic neighbor embedding (tSNE; dark red). We used log2 count transformation for the subset of DR methods that use normalized data. For each data set, we compared the four different number of low-dimensional data. The four numbers we used equal to 0.5%, 1%, 2%, and 3% of the total number of cells in big data and equal to 2, 6, 14, and 20 in small data. For convenience, we only listed 0.5%, 1%, 2%, and 3% on x-axis. Note that, for tSNE, we only extracted two low-dimensional components due to the limitation of the tSNE software.

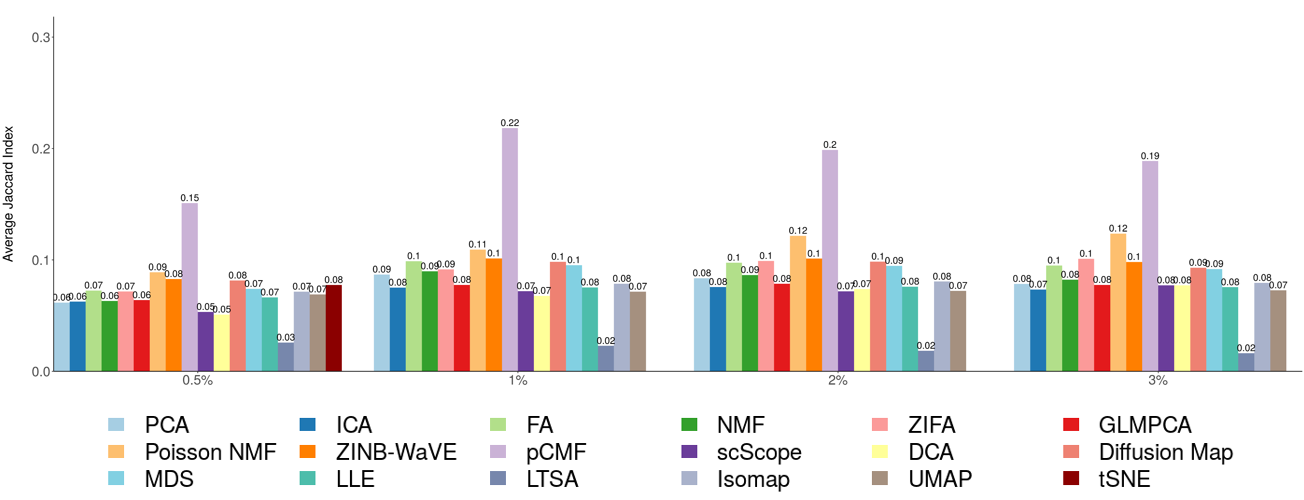

**Supplementary Figure 4. DR method performance evaluated by Jaccard index on cell clustering data sets with 20 neighborhood cells.** We compared 18 DR methods (columns), including factor analysis (FA), principal component analysis (PCA), independent component analysis (ICA), Diffusion Map, nonnegative matrix factorization (NMF), Poisson NMF, zero-inflated factor analysis (ZIFA), zero-inflated negative binomial based wanted variation extraction (ZINB-WaVE), probabilistic count matrix factorization (pCMF), deep count autoencoder network (DCA), scScope, generalized linear model principal component analysis (GLMPCA), multidimensional scaling (MDS), locally linear embedding (LLE), local tangent space alignment (LTSA), Isomap, uniform manifold approximation and projection (UMAP), and t-distributed stochastic neighbor embedding (tSNE). We evaluated their performance on 14 real scRNAseq data sets (UMI-based data are colored as blue while non-UMI based data are colored as purple) and 2 simulated data sets (rows). The simulated data based on Kumar data is labeled with #. We used log2 count transformation for the subset of DR methods that use normalized data. For each data set, we compared the four different number of low-dimensional data. The four numbers we used equal to 0.5%, 1%, 2%, and 3% of the total number of cells in big data and equal to 2, 6, 14, and 20 in small data (which are labeled with *). No results for ICA are shown in the Pancreatic data (grey fills) because ICA cannot handle the large number of features in the data. Note that, for tSNE, we only extracted two low-dimensional components due to the limitation of the tSNE software.

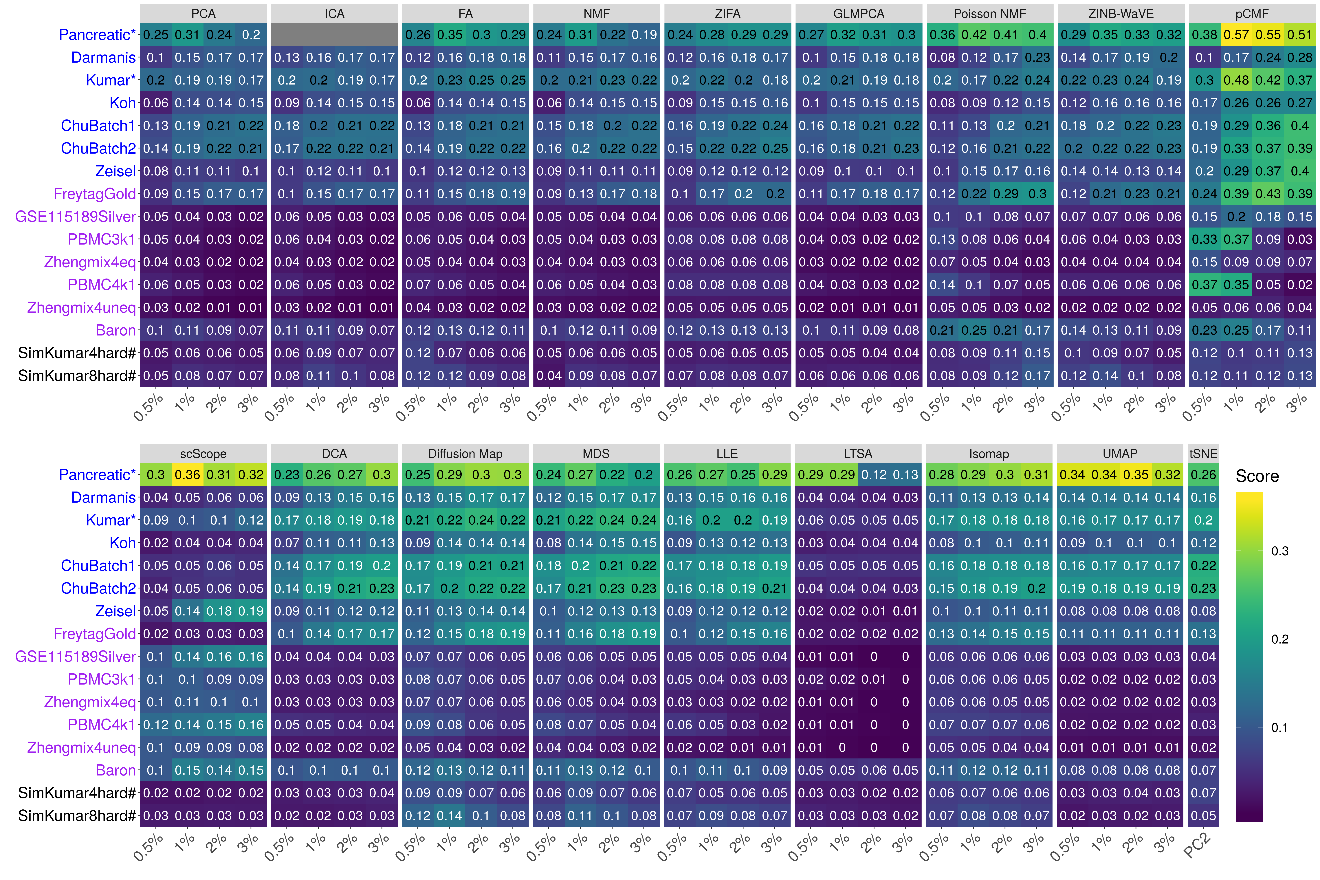

**Supplementary Figure 5. Bar plots show the average Jaccard index of different DR methods with 20 neighborhood cells based on cell clustering data.** The performance is measured by average Jaccard index across 16 data sets. We compared 18 DR methods, including factor analysis (FA; light green), principal component analysis (PCA; light blue), independent component analysis (ICA; blue), Diffusion Map (pink), nonnegative matrix factorization (NMF; green), Poisson NMF(light orange), zero-inflated factor analysis (ZIFA; light pink), zero-inflated negative binomial based wanted variation extraction (ZINB-WaVE; orange), probabilistic count matrix factorization (pCMF; light purple), deep count autoencoder network (DCA; yellow), scScope (purple), generalized linear model principal component analysis (GLMPCA; red), multidimensional scaling (MDS; cyan), locally linear embedding (LLE; blue green), local tangent space alignment (LTSA; teal blue), Isomap (grey), uniform manifold approximation and projection (UMAP; brown), and t-distributed stochastic neighbor embedding (tSNE; dark red). We used log2 count transformation for the subset of DR methods that use normalized data. For each data set, we compared the four different number of low-dimensional data. The four numbers we used equal to 0.5%, 1%, 2%, and 3% of the total number of cells in big data and equal to 2, 6, 14, and 20 in small data. For convenience, we only listed 0.5%, 1%, 2%, and 3% on x-axis. Note that, for tSNE, we only extracted two low-dimensional components due to the limitation of the tSNE software.

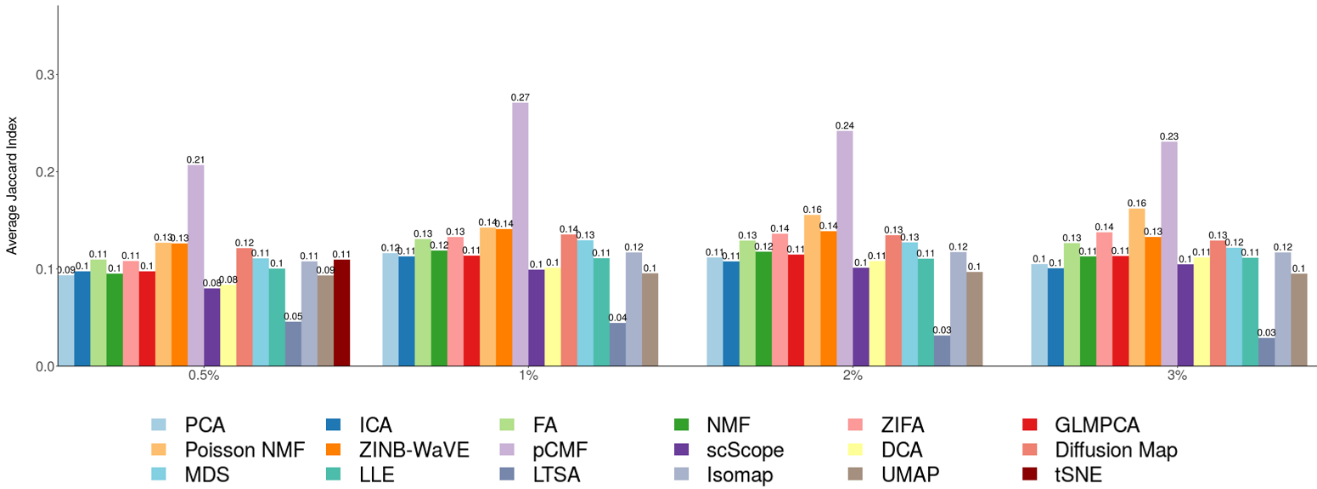

**Supplementary Figure 6. DR method performance evaluated by Jaccard index on cell clustering data sets with 30 neighborhood cells.** We compared 18 DR methods (columns), including factor analysis (FA), principal component analysis (PCA), independent component analysis (ICA), Diffusion Map, nonnegative matrix factorization (NMF), Poisson NMF, zero-inflated factor analysis (ZIFA), zero-inflated negative binomial based wanted variation extraction (ZINB-WaVE), probabilistic count matrix factorization (pCMF), deep count autoencoder network (DCA), scScope, generalized linear model principal component analysis (GLMPCA), multidimensional scaling (MDS), locally linear embedding (LLE), local tangent space alignment (LTSA), Isomap, uniform manifold approximation and projection (UMAP), and t-distributed stochastic neighbor embedding (tSNE). We evaluated their performance on 14 real scRNAseq data sets (UMI-based data are colored as blue while non-UMI based data are colored as purple) and 2 simulated data sets (rows). The simulated data based on Kumar data is labeled with #. We used log2 count transformation for the subset of DR methods that use normalized data. For each data set, we compared the four different number of low-dimensional data. The four numbers we used equal to 0.5%, 1%, 2%, and 3% of the total number of cells in big data and equal to 2, 6, 14, and 20 in small data (which are labeled with *). No results for ICA are shown in the Pancreatic data (grey fills) because ICA cannot handle the large number of features in the data. Note that, for tSNE, we only extracted two low-dimensional components due to the limitation of the tSNE software.

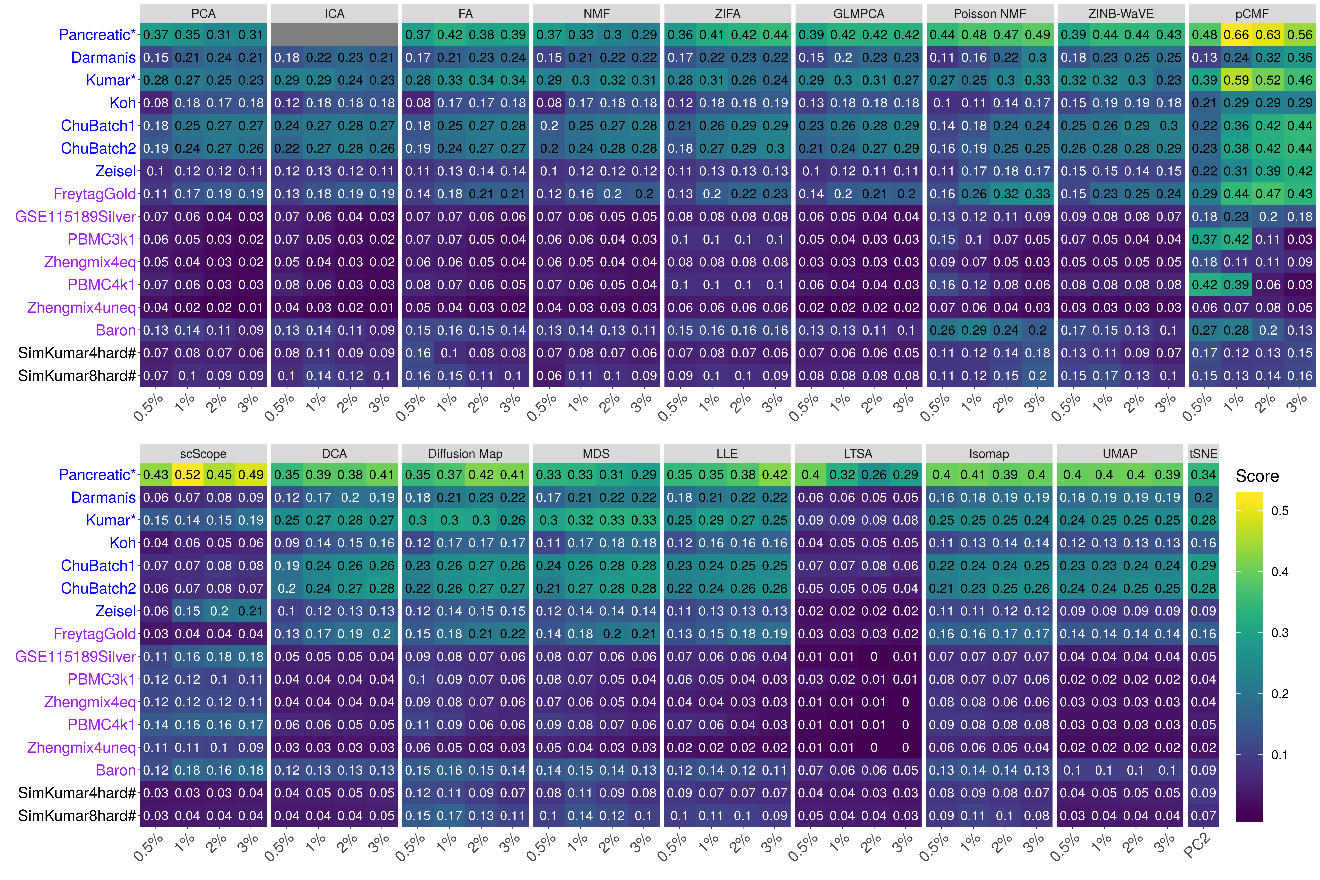

**Supplementary Figure 7. Bar plots show the average Jaccard index of different DR methods with 30 neighborhood cells based on cell clustering data.** The performance is measured by average Jaccard index across 16 data sets. We compared 18 DR methods, including factor analysis (FA; light green), principal component analysis (PCA; light blue), independent component analysis (ICA; blue), Diffusion Map (pink), nonnegative matrix factorization (NMF; green), Poisson NMF(light orange), zero-inflated factor analysis (ZIFA; light pink), zero-inflated negative binomial based wanted variation extraction (ZINB-WaVE; orange), probabilistic count matrix factorization (pCMF; light purple), deep count autoencoder network (DCA; yellow), scScope (purple), generalized linear model principal component analysis (GLMPCA; red), multidimensional scaling (MDS; cyan), locally linear embedding (LLE; blue green), local tangent space alignment (LTSA; teal blue), Isomap (grey), uniform manifold approximation and projection (UMAP; brown), and t-distributed stochastic neighbor embedding (tSNE). We used log2 count transformation for the subset of DR methods that use normalized data. For each data set, we compared the four different number of low-dimensional data. The four numbers we used equal to 0.5%, 1%, 2%, and 3% of the total number of cells in big data and equal to 2, 6, 14, and 20 in small data. For convenience, we only listed 0.5%, 1%, 2%, and 3% on x-axis. Note that, for tSNE, we only extracted two low-dimensional components due to the limitation of the tSNE software.

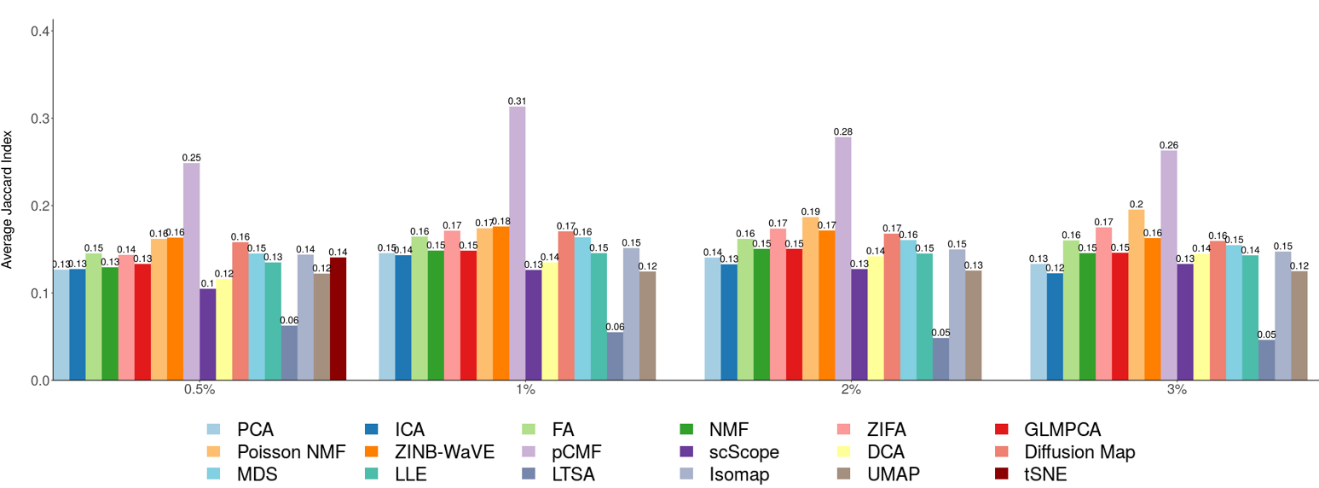

**Supplementary Figure 8. Bar plots show the average Jaccard index of different DR methods on UMI-based data and nonUMI-based data with 30 neighborhood cells.** The performance is measured by average Jaccard index across UMI-based data (**A**; 7 data sets) or nonUMI-based data (**B**; 7 data sets). We compared 18 DR methods, including factor analysis (FA; light green), principal component analysis (PCA; light blue), independent component analysis (ICA; blue), Diffusion Map (pink), nonnegative matrix factorization (NMF; green), Poisson NMF(light orange), zero-inflated factor analysis (ZIFA; light pink), zero-inflated negative binomial based wanted variation extraction (ZINB-WaVE; orange), probabilistic count matrix factorization (pCMF; light purple), deep count autoencoder network (DCA; yellow), scScope (purple), generalized linear model principal component analysis (GLMPCA; red), multidimensional scaling (MDS; cyan), locally linear embedding (LLE; blue green), local tangent space alignment (LTSA; teal blue), Isomap (grey), uniform manifold approximation and projection (UMAP; brown), and t-distributed stochastic neighbor embedding (tSNE; dark red). We used log2 count transformation for the subset of DR methods that use normalized data. For each data set, we compared the four different number of low-dimensional data. The four numbers we used equal to 0.5%, 1%, 2%, and 3% of the total number of cells in big data and equal to 2, 6, 14, and 20 in small data. For convenience, we only listed 0.5%, 1%, 2%, and 3% on x-axis. Note that, for tSNE, we only extracted two low-dimensional components due to the limitation of the tSNE software.

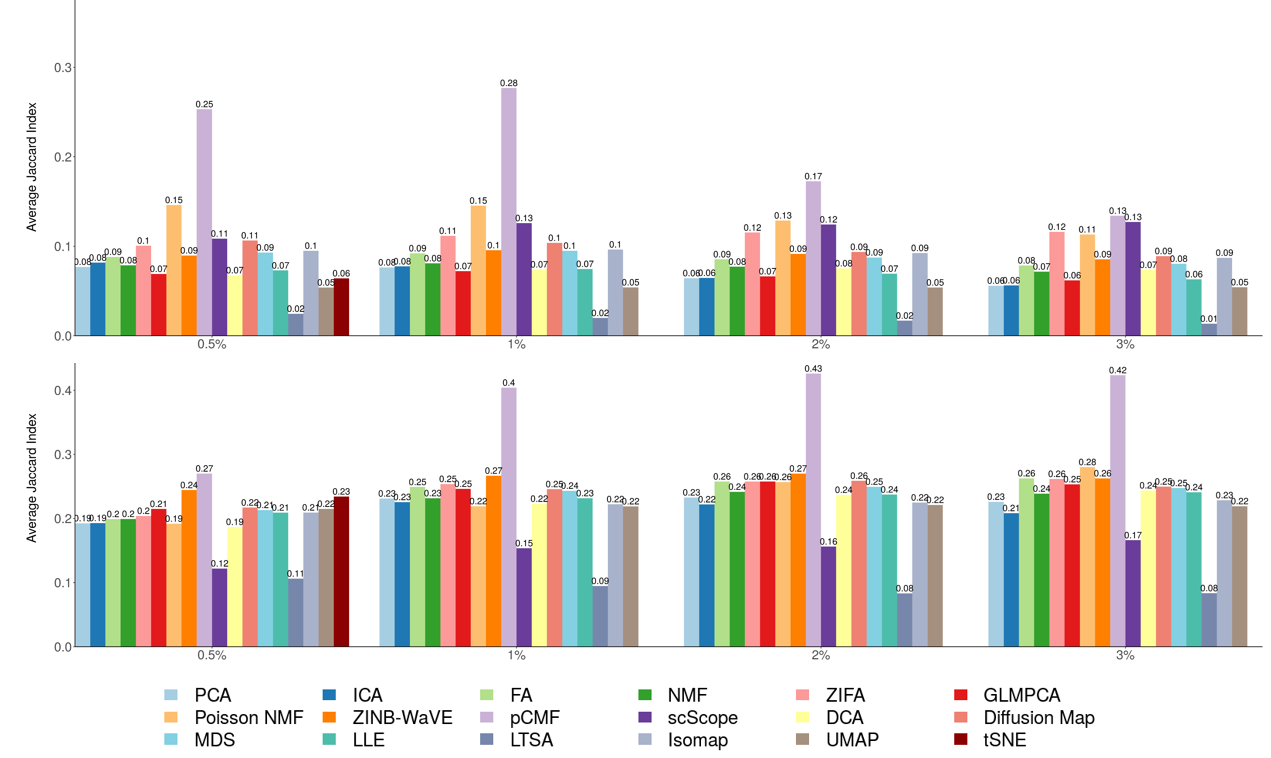

**Supplementary Figure 9. DR method performance evaluated by Jaccard index on trajectory inference data sets with 10 neighborhood cells.** We compared 18 DR methods (columns), including factor analysis (FA), principal component analysis (PCA), independent component analysis (ICA), Diffusion Map, nonnegative matrix factorization (NMF), Poisson NMF, zero-inflated factor analysis (ZIFA), zero-inflated negative binomial based wanted variation extraction (ZINB-WaVE), probabilistic count matrix factorization (pCMF), deep count autoencoder network (DCA), scScope, generalized linear model principal component analysis (GLMPCA), multidimensional scaling (MDS), locally linear embedding (LLE), local tangent space alignment (LTSA), Isomap, and uniform manifold approximation and projection (UMAP), and t-distributed stochastic neighbor embedding (tSNE). We evaluated their performance on 14 real scRNAseq data sets (rows) in terms of lineage inference accuracy. The simulated data based on Kumar data is labeled with #. We used log2 count transformation for the subset of DR methods that use normalized data. For each data set, we compared the four different number of low-dimensional data. The four numbers we used equal to 2, 6, 14, and 20. Note that, for tSNE, we only extracted two low-dimensional components due to the limitation of the tSNE software.

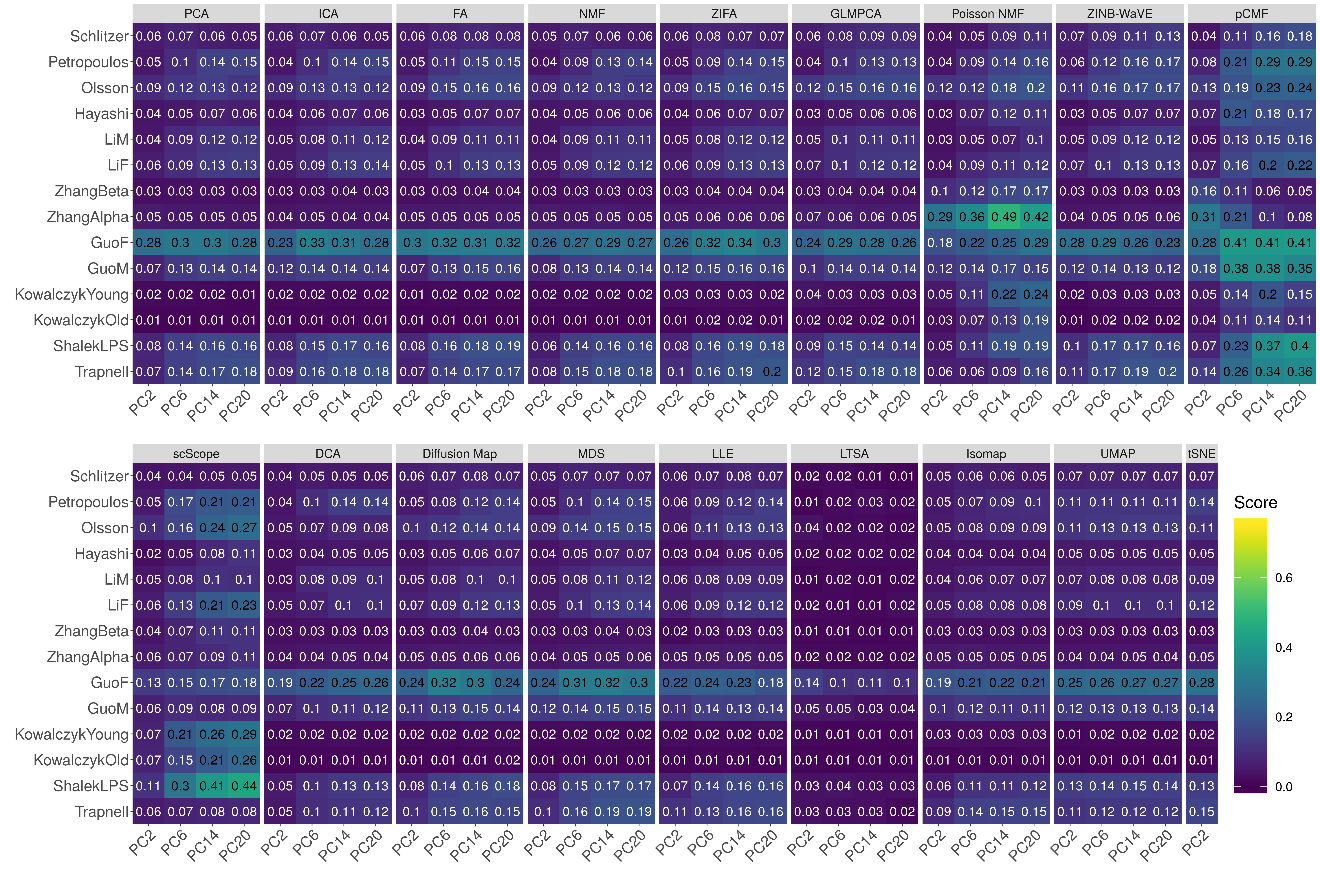

**Supplementary Figure 10. Bar plots show the average Jaccard index of different DR methods with 10 neighborhood cells based on trajectory inference data.** The performance is measured by average Jaccard index across 16 data sets. We compared 18 DR methods, including factor analysis (FA; light green), principal component analysis (PCA; light blue), independent component analysis (ICA; blue), Diffusion Map (pink), nonnegative matrix factorization (NMF; green), Poisson NMF(light orange), zero-inflated factor analysis (ZIFA; light pink), zero-inflated negative binomial based wanted variation extraction (ZINB-WaVE; orange), probabilistic count matrix factorization (pCMF; light purple), deep count autoencoder network (DCA; yellow), scScope (purple), generalized linear model principal component analysis (GLMPCA; red), multidimensional scaling (MDS; cyan), locally linear embedding (LLE; blue green), local tangent space alignment (LTSA; teal blue), Isomap (grey), uniform manifold approximation and projection (UMAP; brown), and t-distributed stochastic neighbor embedding (tSNE; dark red). We used log2 count transformation for the subset of DR methods that use normalized data. For each data set, we compared the four different number of low-dimensional data. The four numbers we used equal to 2, 6, 14, and 20 on x-axis. Note that, for tSNE, we only extracted two low-dimensional components due to the limitation of the tSNE software.

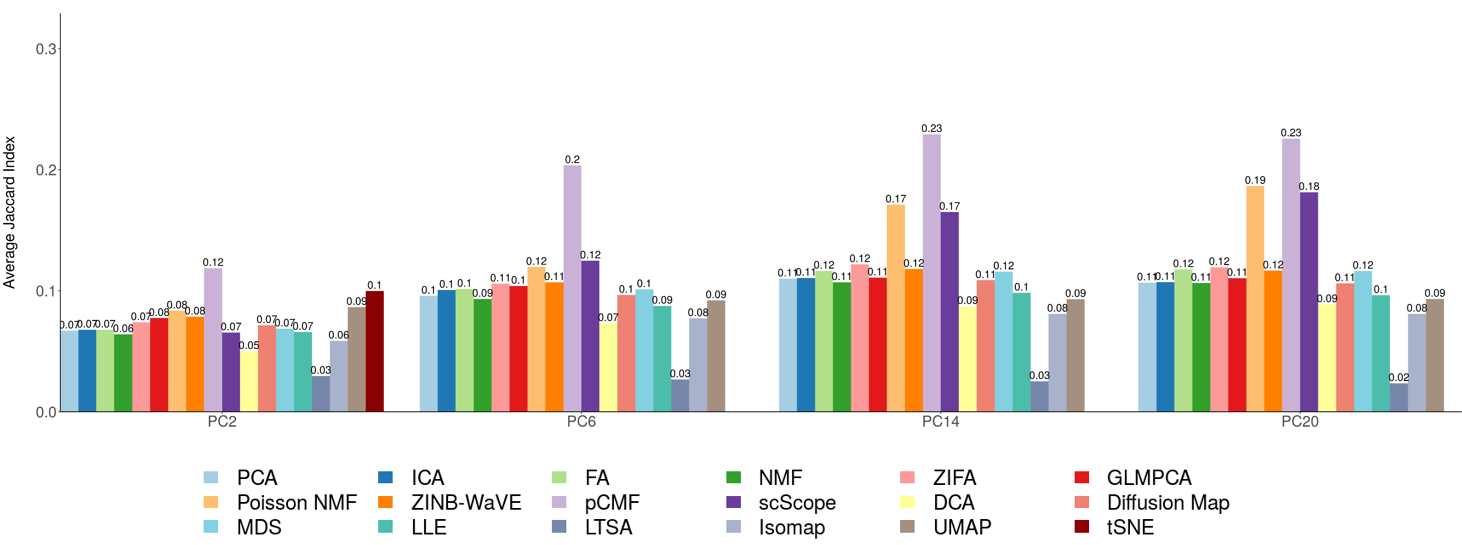

**Supplementary Figure 11. DR method performance evaluated by Jaccard index on trajectory inference data sets with 20 neighborhood cells.** We compared 18 DR methods (columns), including factor analysis (FA), principal component analysis (PCA), independent component analysis (ICA), Diffusion Map, nonnegative matrix factorization (NMF), Poisson NMF, zero-inflated factor analysis (ZIFA), zero-inflated negative binomial based wanted variation extraction (ZINB-WaVE), probabilistic count matrix factorization (pCMF), deep count autoencoder network (DCA), scScope, generalized linear model principal component analysis (GLMPCA), multidimensional scaling (MDS), locally linear embedding (LLE), local tangent space alignment (LTSA), Isomap, uniform manifold approximation and projection (UMAP), and t-distributed stochastic neighbor embedding (tSNE). We evaluated their performance on 14 real scRNAseq data sets (rows) in terms of lineage inference accuracy. The simulated data based on Kumar data is labeled with #. We used log2 count transformation for the subset of DR methods that use normalized data. For each data set, we compared the four different number of low-dimensional data. The four numbers we used equal to 2, 6, 14, and 20. Note that, for tSNE, we only extracted two low-dimensional components due to the limitation of the tSNE software.

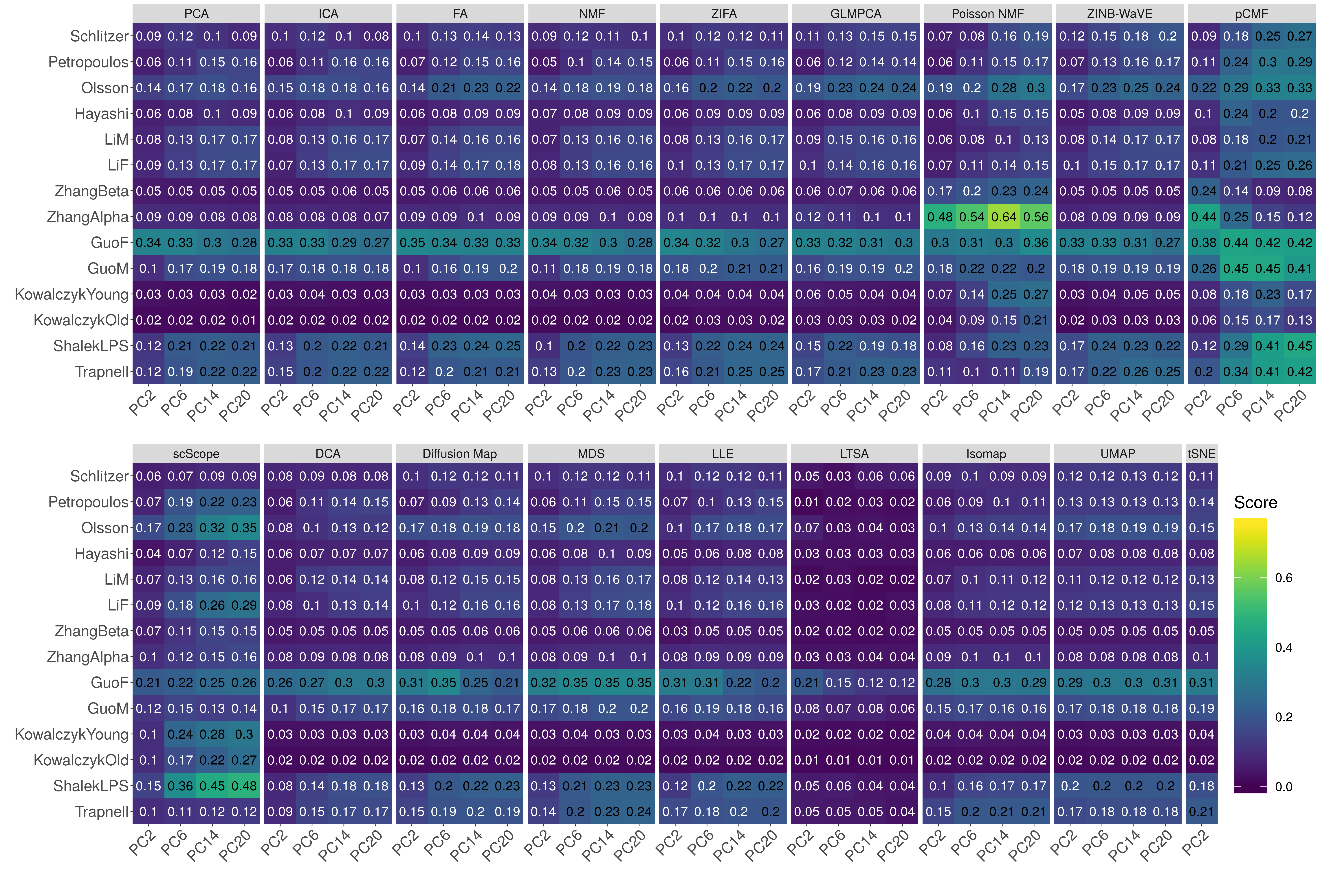

**Supplementary Figure 12. Bar plots show the average Jaccard index of different DR methods with 20 neighborhood cells based on trajectory inference data.** The performance is measured by average Jaccard index across 16 data sets. We compared 18 DR methods, including factor analysis (FA; light green), principal component analysis (PCA; light blue), independent component analysis (ICA; blue), Diffusion Map (pink), nonnegative matrix factorization (NMF; green), Poisson NMF(light orange), zero-inflated factor analysis (ZIFA; light pink), zero-inflated negative binomial based wanted variation extraction (ZINB-WaVE; orange), probabilistic count matrix factorization (pCMF; light purple), deep count autoencoder network (DCA; yellow), scScope (purple), generalized linear model principal component analysis (GLMPCA; red), multidimensional scaling (MDS; cyan), locally linear embedding (LLE; blue green), local tangent space alignment (LTSA; teal blue), Isomap (grey), uniform manifold approximation and projection (UMAP; brown), and t-distributed stochastic neighbor embedding (tSNE; dark red). We used log2 count transformation for the subset of DR methods that use normalized data. For each data set, we compared the four different number of low-dimensional data. The four numbers we used equal to 2, 6, 14, and 20 on x-axis. Note that, for tSNE, we only extracted two low-dimensional components due to the limitation of the tSNE software.

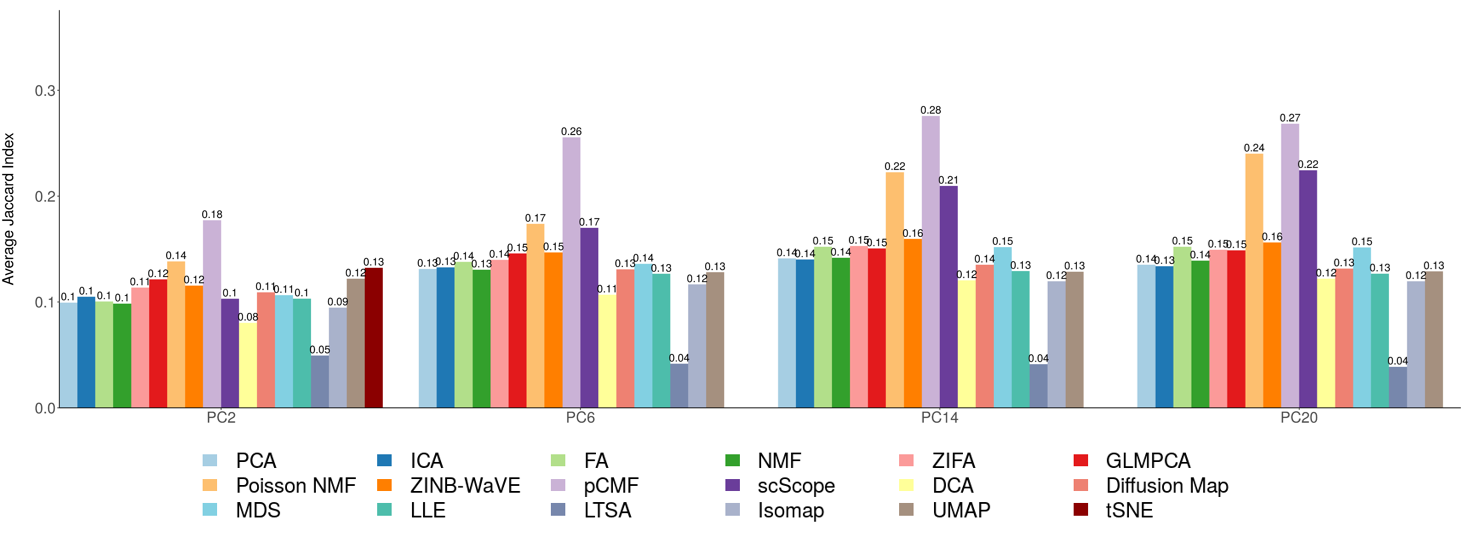

**Supplementary Figure 13. DR method performance evaluated by Jaccard index on trajectory inference data sets with 30 neighborhood cells.** We compared 18 DR methods (columns), including factor analysis (FA), principal component analysis (PCA), independent component analysis (ICA), Diffusion Map, nonnegative matrix factorization (NMF), Poisson NMF, zero-inflated factor analysis (ZIFA), zero-inflated negative binomial based wanted variation extraction (ZINB-WaVE), probabilistic count matrix factorization (pCMF), deep count autoencoder network (DCA), scScope, generalized linear model principal component analysis (GLMPCA), multidimensional scaling (MDS), locally linear embedding (LLE), local tangent space alignment (LTSA), Isomap, uniform manifold approximation and projection (UMAP), and t-distributed stochastic neighbor embedding (tSNE). We evaluated their performance on 14 real scRNAseq data sets (rows) in terms of lineage inference accuracy. The simulated data based on Kumar data is labeled with #. We used log2 count transformation for the subset of DR methods that use normalized data. For each data set, we compared the four different number of low-dimensional data. The four numbers we used equal to 2, 6, 14, and 20. Note that, for tSNE, we only extracted two low-dimensional components due to the limitation of the tSNE software.

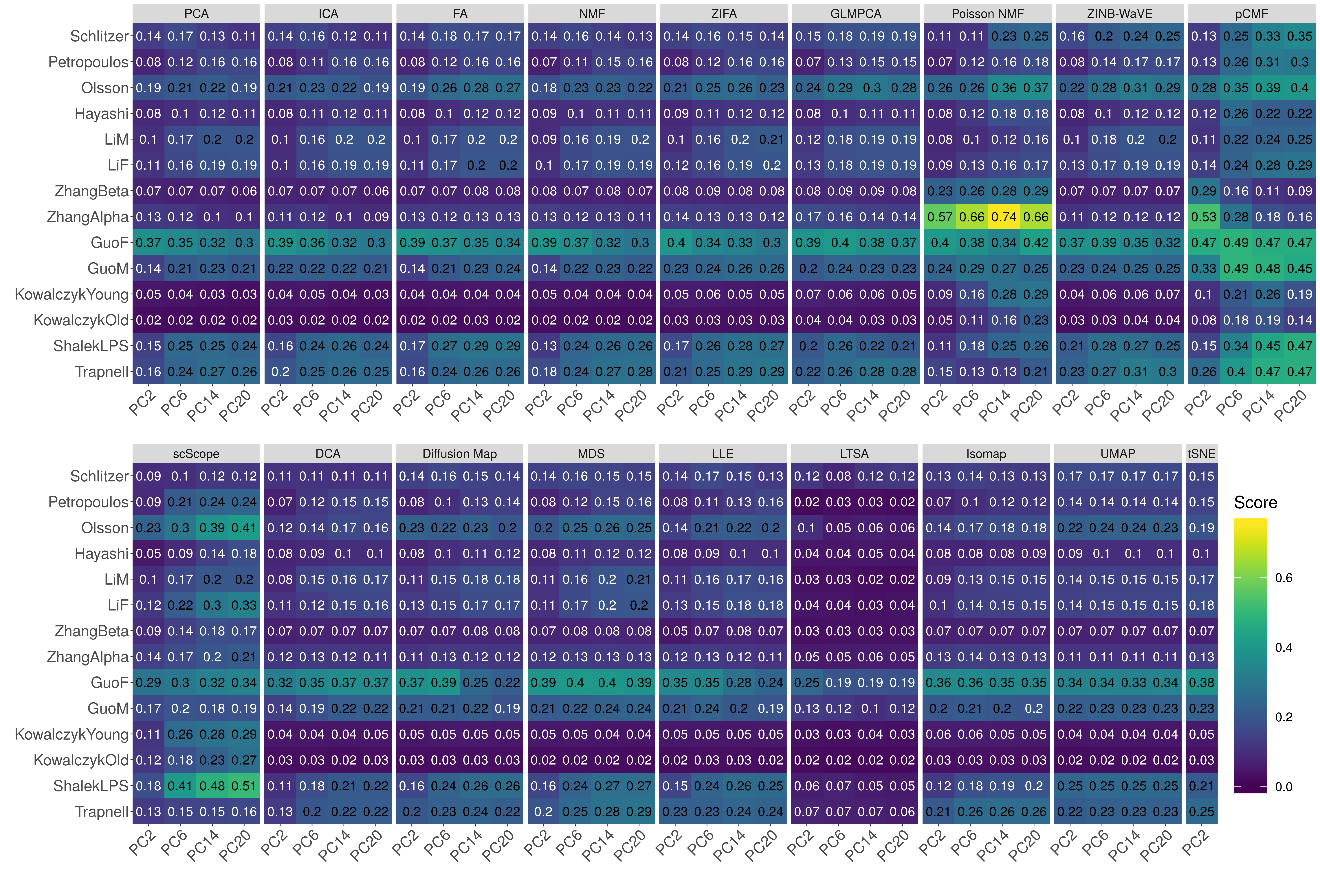

**Supplementary Figure 14. Bar plots show the average Jaccard index of different DR methods with 30 neighborhood cells based on trajectory inference data.** The performance is measured by average Jaccard index across 16 data sets. We compared 18 DR methods, including factor analysis (FA; light green), principal component analysis (PCA; light blue), independent component analysis (ICA; blue), Diffusion Map (pink), nonnegative matrix factorization (NMF; green), Poisson NMF(light orange), zero-inflated factor analysis (ZIFA; light pink), zero-inflated negative binomial based wanted variation extraction (ZINB-WaVE; orange), probabilistic count matrix factorization (pCMF; light purple), deep count autoencoder network (DCA; yellow), scScope (purple), generalized linear model principal component analysis (GLMPCA; red), multidimensional scaling (MDS; cyan), locally linear embedding (LLE; blue green), local tangent space alignment (LTSA; teal blue), Isomap (grey), uniform manifold approximation and projection (UMAP; brown), and t-distributed stochastic neighbor embedding (tSNE; dark red). We used log2 count transformation for the subset of DR methods that use normalized data. For each data set, we compared the four different number of low-dimensional data. The four numbers we used equal to 2, 6, 14, and 20 on x-axis. Note that, for tSNE, we only extracted two low-dimensional components due to the limitation of the tSNE software.

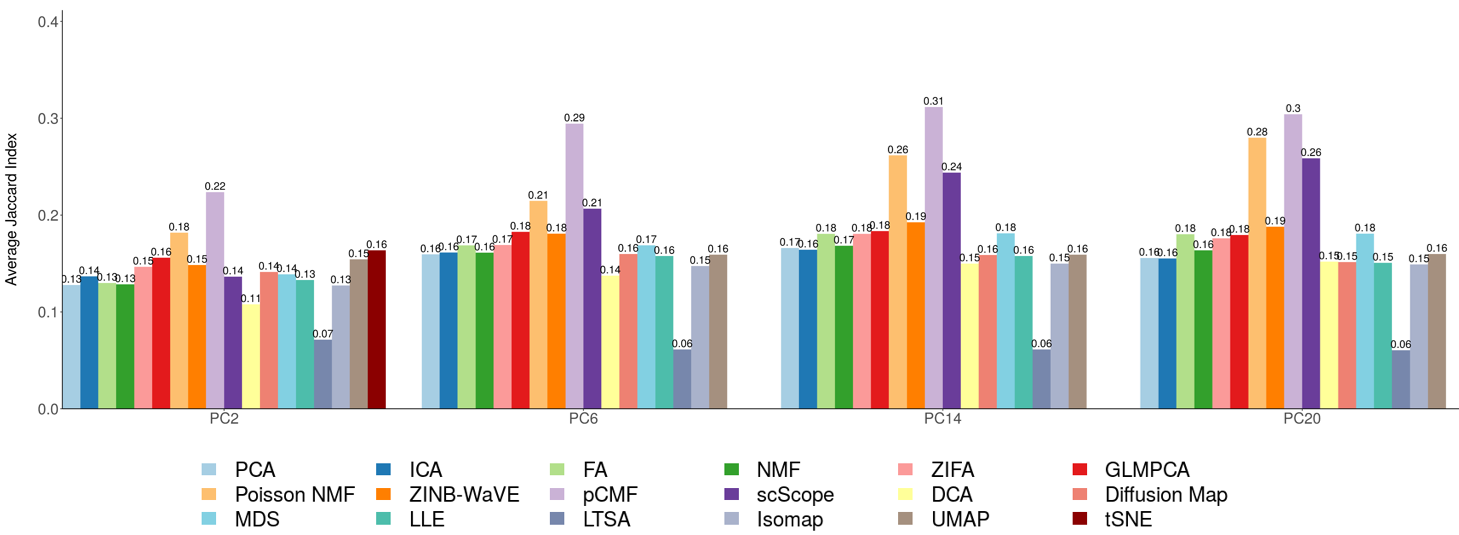

**Supplementary Figure 15. DR method performance evaluated by ARI in the downstream cell clustering analysis with *k*-means.** We compared 18 DR methods (columns), including factor analysis (FA), principal component analysis (PCA), independent component analysis (ICA), Diffusion Map, nonnegative matrix factorization (NMF), Poisson NMF, zero-inflated factor analysis (ZIFA), zero-inflated negative binomial based wanted variation extraction (ZINB-WaVE), probabilistic count matrix factorization (pCMF), deep count autoencoder network (DCA), scScope, generalized linear model principal component analysis (GLMPCA), multidimensional scaling (MDS), locally linear embedding (LLE), local tangent space alignment (LTSA), Isomap, uniform manifold approximation and projection (UMAP), and t-distributed stochastic neighbor embedding (tSNE). We evaluated their performance on 14 real scRNAseq data sets (UMI-based data are labeled as purple; nonUMI–based data are labeled as blue) and 2 simulated data sets (rows). The simulated data based on Kumar data is labeled with #. The performance of each DR method is measured by normalized mutual information (NMI). For each data set, we compared the four different number of low-dimensional components. The four numbers equal to 0.5%, 1%, 2%, and 3% of the total number of cells in big data and equal to 2, 6, 14, and 20 in small data (which are labeled with *). For convenience, we only listed 0.5%, 1%, 2%, and 3% on x-axis. No results for ICA and LTSA are shown in the table (grey fills) because ICA cannot handle the large number of features in that data or estimated zero low-dimensional components. Note that, for tSNE, we only extracted two low-dimensional components due to the limitation of the tSNE software.

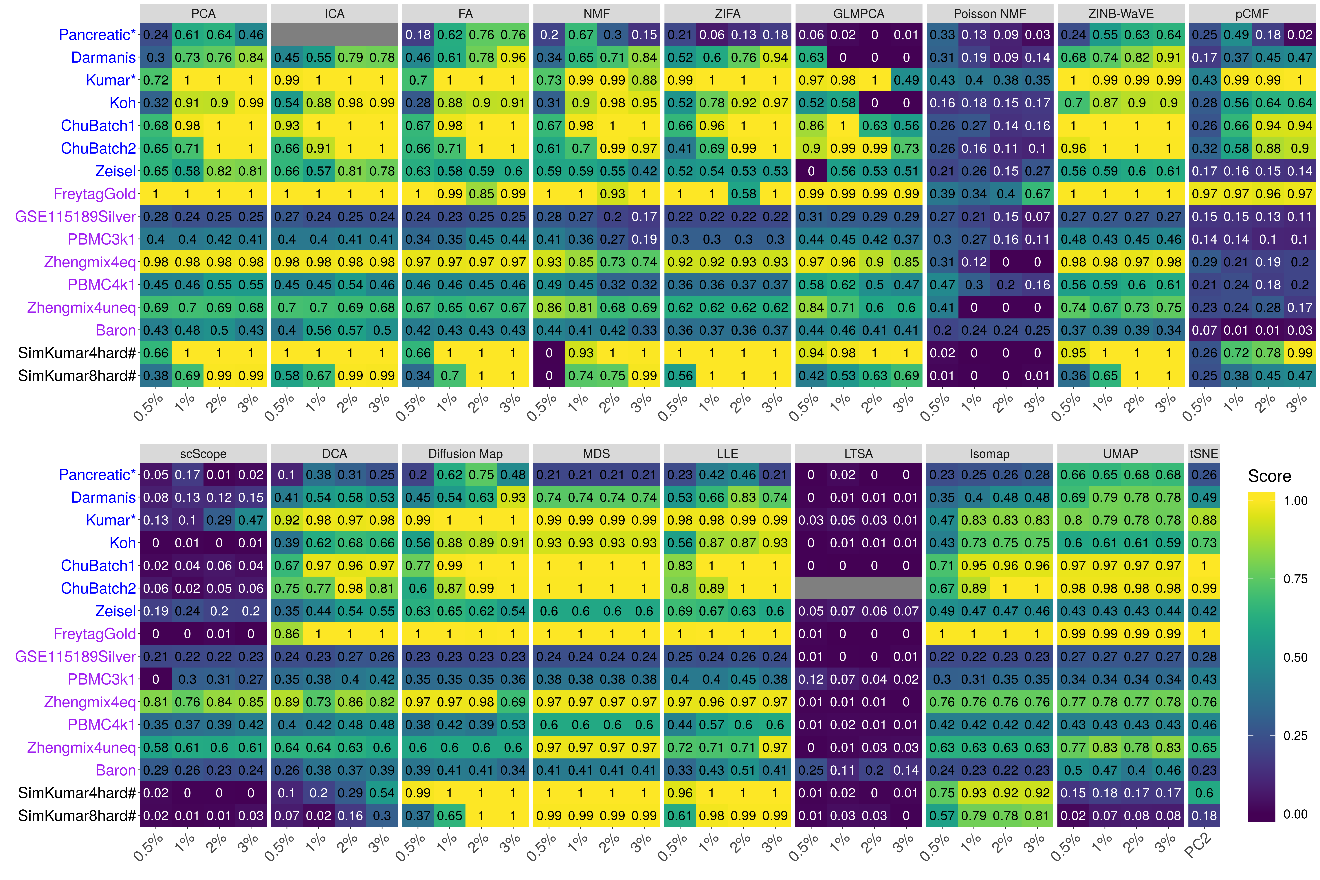

**Supplementary Figure 16. Bar plots show the average performance of different DR methods based on *k*-means clustering.** The performance is measured by average normalized mutual information (NMI; **A**) or average adjusted rand index (ARI; **B**), both averaged across 16 data sets. We compared 18 DR methods, including factor analysis (FA; light green), principal component analysis (PCA; light blue), independent component analysis (ICA; blue), Diffusion Map (pink), nonnegative matrix factorization (NMF; green), Poisson NMF(light orange), zero-inflated factor analysis (ZIFA; light pink), zero-inflated negative binomial based wanted variation extraction (ZINB-WaVE; orange), probabilistic count matrix factorization (pCMF; light purple), deep count autoencoder network (DCA; yellow), scScope (purple), generalized linear model principal component analysis (GLMPCA; red), multidimensional scaling (MDS; cyan), locally linear embedding (LLE; blue green), local tangent space alignment (LTSA; teal blue), Isomap (grey), uniform manifold approximation and projection (UMAP; brown), and t-distributed stochastic neighbor embedding (tSNE; dark red). We used log2 Count transformation for the subset of DR methods that use normalized data. For each data set, we compared the four different number of low-dimensional data. The four numbers we used equal to 0.5%, 1%, 2%, and 3% of the total number of cells in big data and equal to 2, 6, 14, and 20 in small data. For convenience, we only listed 0.5%, 1%, 2%, and 3% on x-axis. Note that, for tSNE, we only extracted two low-dimensional components due to the limitation of the tSNE software.

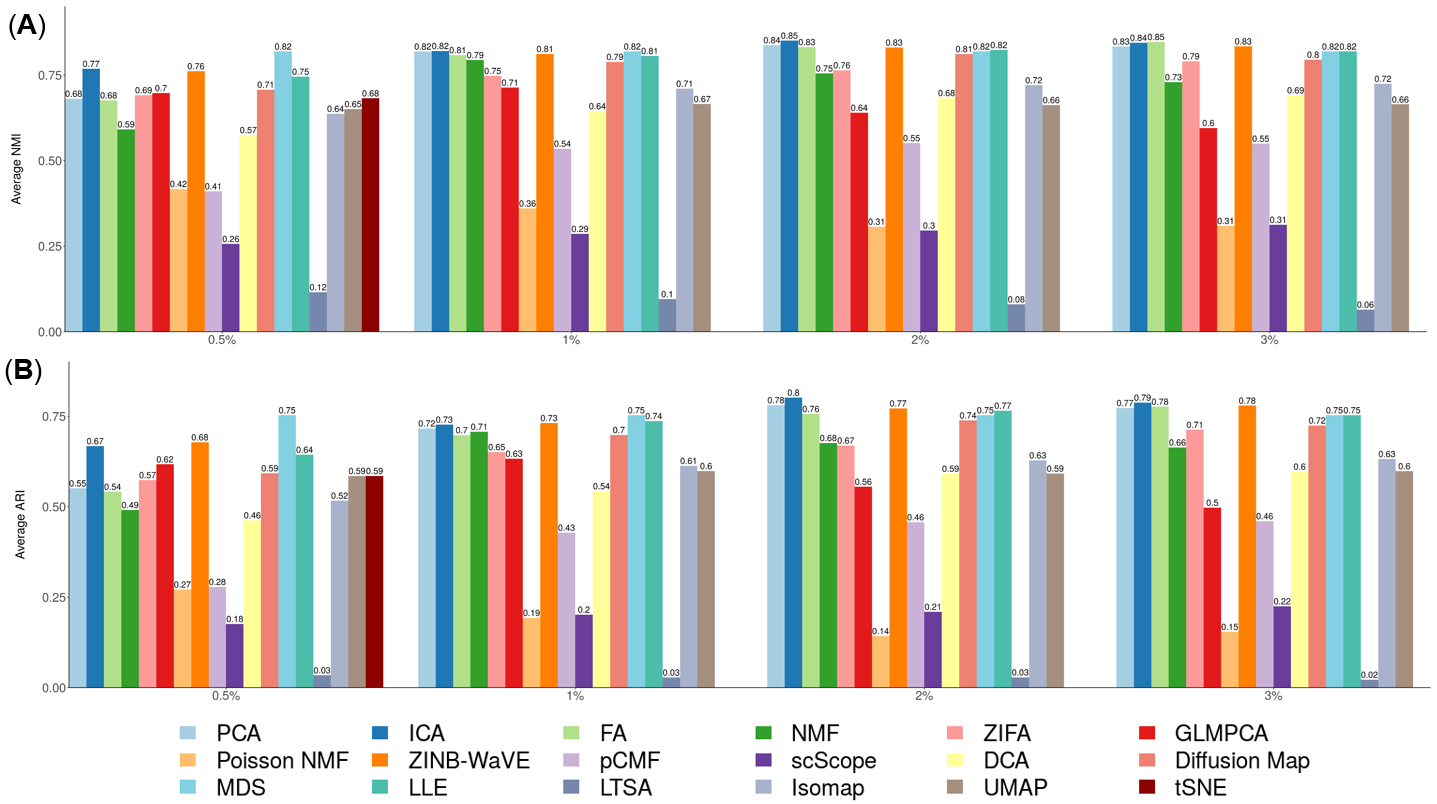

**Supplementary Figure 17. DR method performance evaluated by NMI in the downstream cell clustering analysis with hierarchical clustering.** We compared 17 DR methods (columns), including factor analysis (FA), principal component analysis (PCA), independent component analysis (ICA), Diffusion Map, nonnegative matrix factorization (NMF), Poisson NMF, zero-inflated factor analysis (ZIFA), zero-inflated negative binomial based wanted variation extraction (ZINB-WaVE), probabilistic count matrix factorization (pCMF), deep count autoencoder network (DCA), generalized linear model principal component analysis (GLMPCA), multidimensional scaling (MDS), locally linear embedding (LLE), local tangent space alignment (LTSA), Isomap, uniform manifold approximation and projection (UMAP), and t-distributed stochastic neighbor embedding (tSNE). We evaluated their performance on 14 real scRNAseq data sets and 2 simulated data sets (rows). The simulated data based on Kumar data is labeled with #. We used log2 Count transformation for the subset of DR methods that use normalized data. For each data set, we compared the four different number of low-dimensional data. The four numbers we used equal to 0.5%, 1%, 2%, and 3% of the total number of cells in big data and equal to 2, 6, 14, and 20 in small data (which were labeled with *). For convenience, we only listed 0.5%, 1%, 2%, and 3% on x-axis. No results for ICA, tSNE or GLMPCA are shown in the table (grey fills) because ICA cannot handle the large number of features; tSNE and GLMPCA have error when clustering cells. Note that, for tSNE, we only extracted two low-dimensional components due to the limitation of the tSNE software.

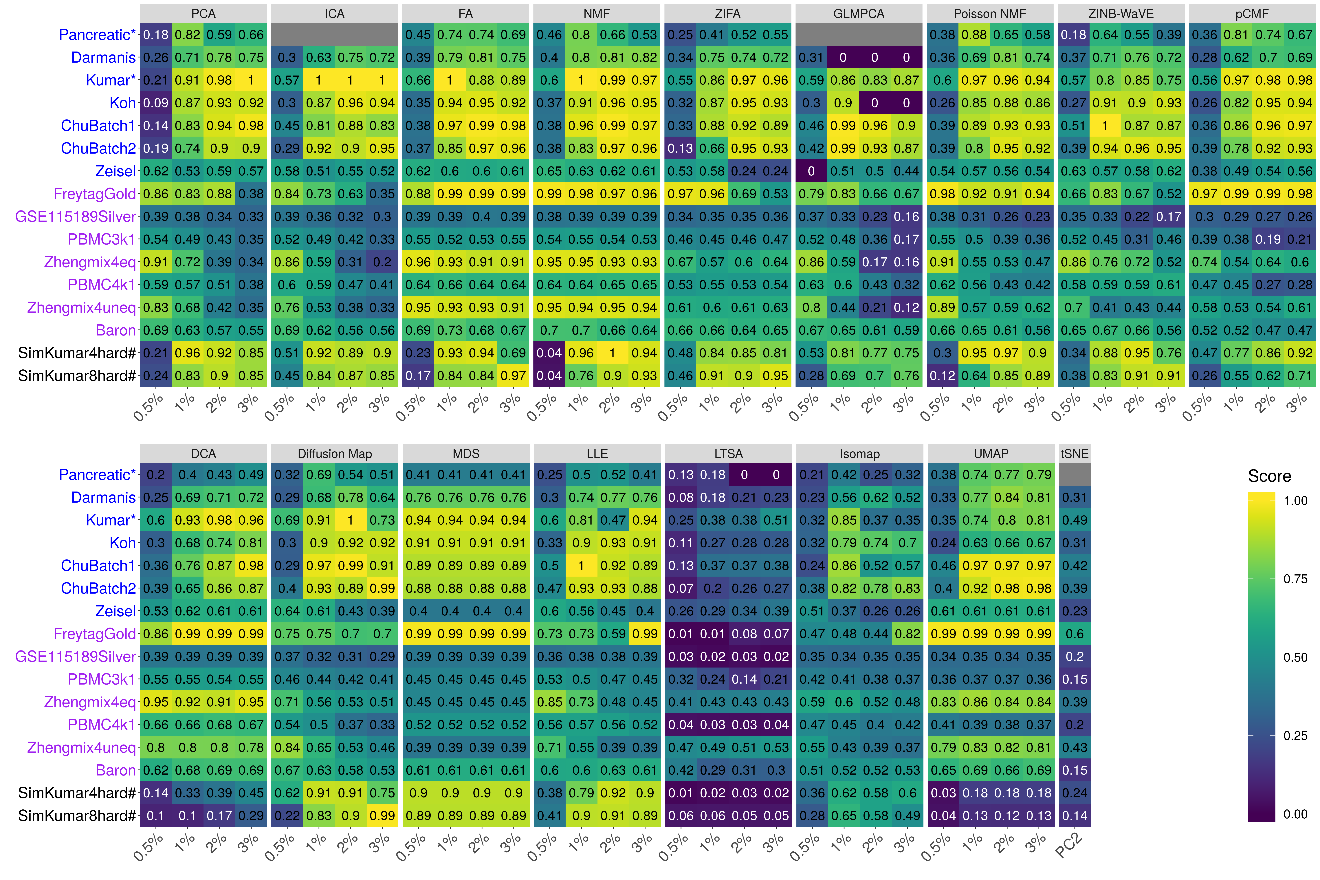

**Supplementary Figure 18. DR method performance evaluated by ARI in the downstream cell clustering analysis with hierarchical clustering.** We compared 17 DR methods (columns), including factor analysis (FA), principal component analysis (PCA), independent component analysis (ICA), Diffusion Map, nonnegative matrix factorization (NMF), Poisson NMF, zero-inflated factor analysis (ZIFA), zero-inflated negative binomial based wanted variation extraction (ZINB-WaVE), probabilistic count matrix factorization (pCMF), deep count autoencoder network (DCA), generalized linear model principal component analysis (GLMPCA), multidimensional scaling (MDS), locally linear embedding (LLE), local tangent space alignment (LTSA), Isomap, uniform manifold approximation and projection (UMAP), and t-distributed stochastic neighbor embedding (tSNE). We evaluated their performance on 14 real scRNAseq data sets and 2 simulated data sets (rows). The simulated data based on Kumar data is labeled with #. We used log2 Count transformation for the subset of DR methods that use normalized data. For each data set, we compared the four different number of low-dimensional data. The four numbers we used equal to 0.5%, 1%, 2%, and 3% of the total number of cells in big data and equal to 2, 6, 14, and 20 in small data (which were labeled with *). For convenience, we only listed 0.5%, 1%, 2%, and 3% on x-axis. No results for ICA or GLMPCA are shown in the table (grey fills) because ICA cannot handle the large number of features; GLMPCA has error when clustering cells. Note that, for tSNE, we only extracted two low-dimensional components due to the limitation of the tSNE software.

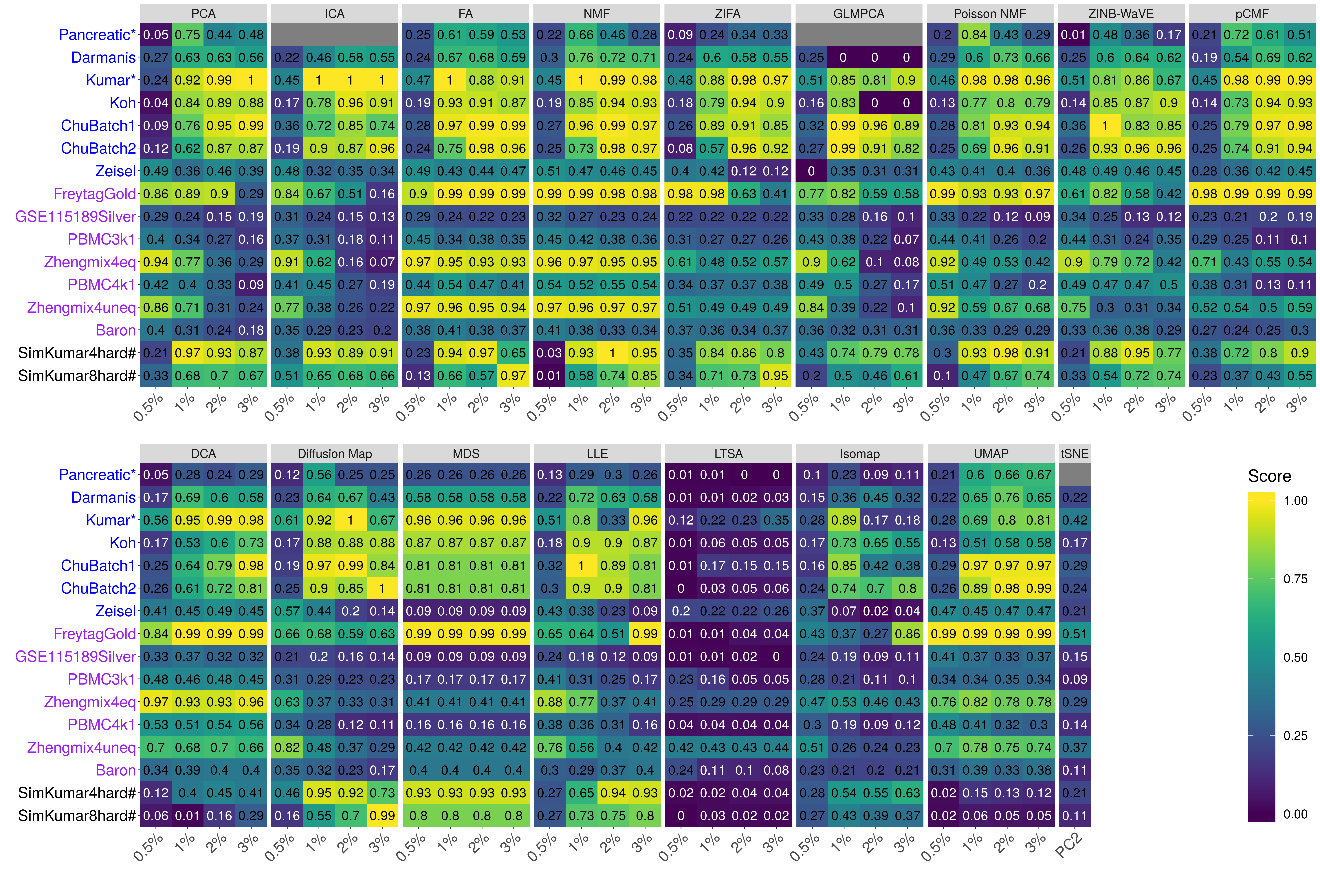

**Supplementary Figure 19. Bar plots show the average performance of different DR methods based on hierarchical clustering.** The performance is measured by normalized mutual information (NMI; **A**) or adjusted rand index (ARI; **B**), both averaged across 16 data sets. We compared 17 DR methods, including factor analysis (FA; light green), principal component analysis (PCA; light blue), independent component analysis (ICA; blue), Diffusion Map (pink), nonnegative matrix factorization (NMF; green), Poisson NMF(light orange), zero-inflated factor analysis (ZIFA; light pink), zero-inflated negative binomial based wanted variation extraction (ZINB-WaVE; orange), probabilistic count matrix factorization (pCMF; light purple), deep count autoencoder network (DCA; yellow), generalized linear model principal component analysis (GLMPCA; red), multidimensional scaling (MDS; cyan), locally linear embedding (LLE; blue green), local tangent space alignment (LTSA; teal blue), Isomap (grey), uniform manifold approximation and projection (UMAP; brown), and t-distributed stochastic neighbor embedding (tSNE; dark red). We used log2 Count transformation for the subset of DR methods that use normalized data. For each data set, we compared the four different number of low-dimensional data. The four numbers we used equal to 0.5%, 1%, 2%, and 3% of the total number of cells in big data and equal to 2, 6, 14, and 20 in small data. For convenience, we only listed 0.5%, 1%, 2%, and 3% on x-axis. Note that, for tSNE, we only extracted two low-dimensional components due to the limitation of the tSNE software.

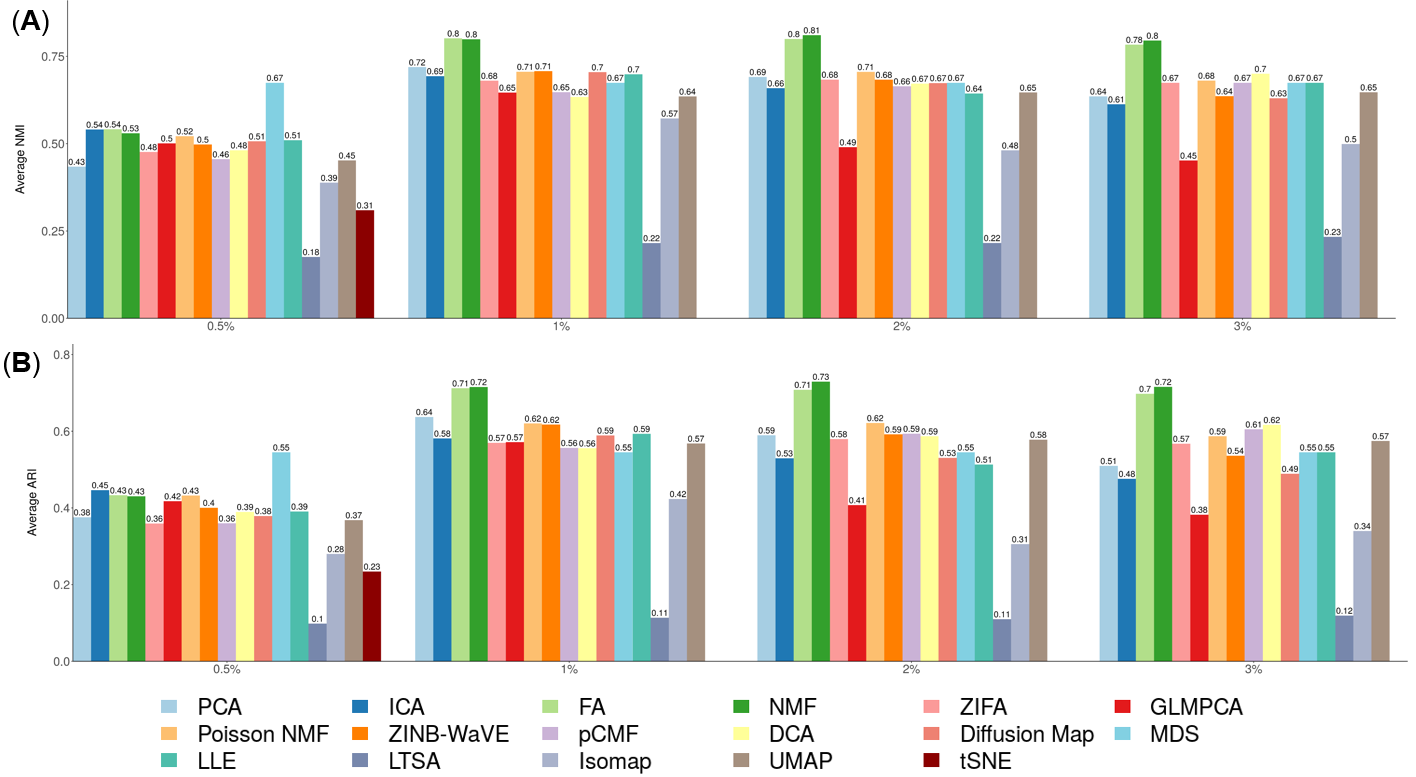

**Supplementary Figure 20. DR method performance evaluated by NMI in the downstream cell clustering analysis with Louvain clustering.** We compared 18 DR methods (columns), including factor analysis (FA), principal component analysis (PCA), independent component analysis (ICA), Diffusion Map, nonnegative matrix factorization (NMF), Poisson NMF, zero-inflated factor analysis (ZIFA), zero-inflated negative binomial based wanted variation extraction (ZINB-WaVE), probabilistic count matrix factorization (pCMF), deep count autoencoder network (DCA), scScope, generalized linear model principal component analysis (GLMPCA), multidimensional scaling (MDS), locally linear embedding (LLE), local tangent space alignment (LTSA), Isomap, uniform manifold approximation and projection (UMAP), and t-distributed stochastic neighbor embedding (tSNE). We evaluated their performance on 14 real scRNAseq data sets and 2 simulated data sets (rows). The simulated data based on Kumar data is labeled with #. We used log2 Count transformation for the subset of DR methods that use normalized data. For each data set, we compared the four different number of low-dimensional data. The four numbers we used equal to 0.5%, 1%, 2%, and 3% of the total number of cells in big data and equal to 2, 6, 14, and 20 in small data (which were labeled with *). For convenience, we only listed 0.5%, 1%, 2%, and 3% on x-axis. No results for ICA are shown in the table (grey fills) because ICA cannot handle the large number of features. Note that, for tSNE, we only extracted two low-dimensional components due to the limitation of the tSNE software.

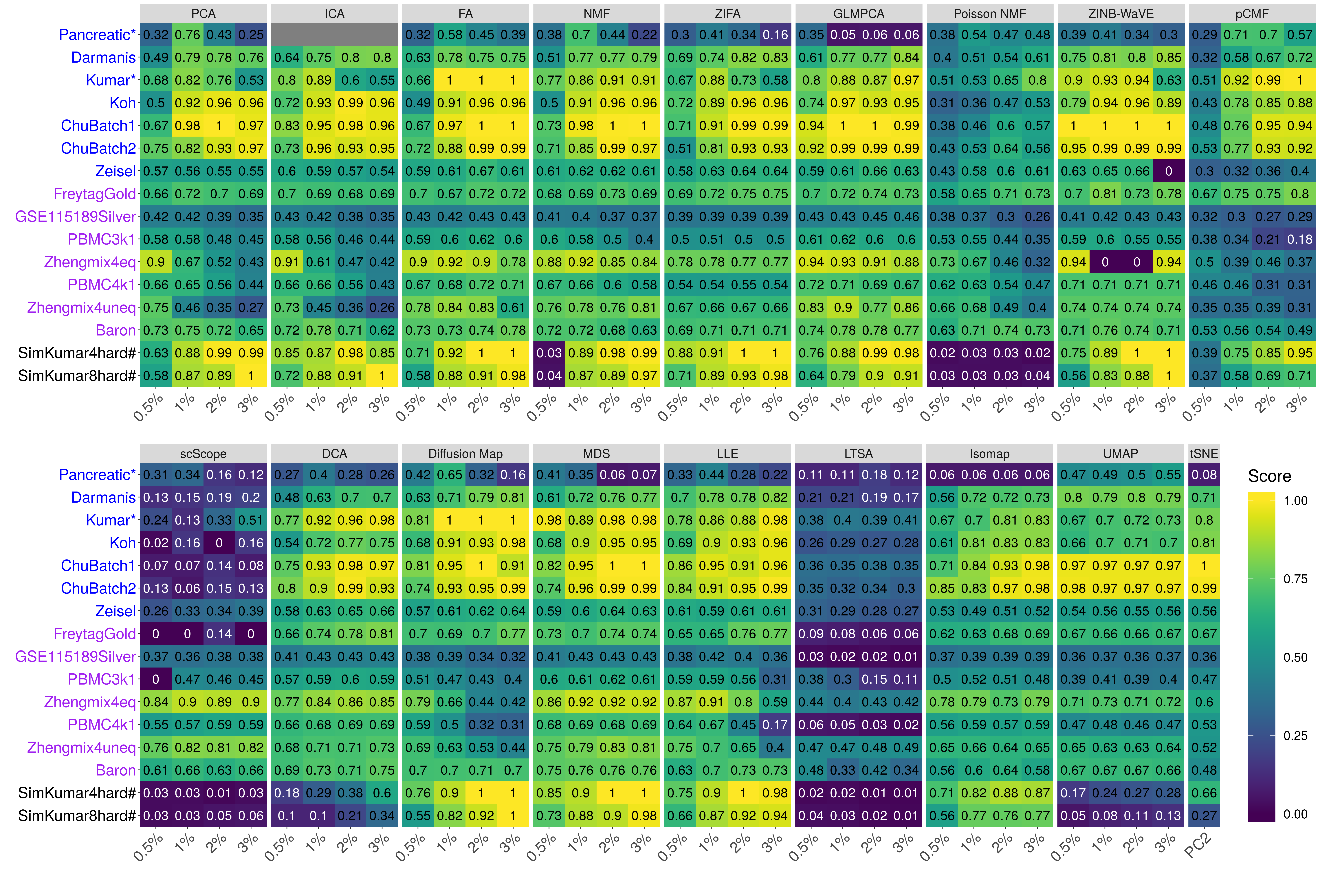

**Supplementary Figure 21. DR method performance evaluated by ARI in the downstream cell clustering analysis with Louvain clustering.** We compared 18 DR methods (columns), including factor analysis (FA), principal component analysis (PCA), independent component analysis (ICA), Diffusion Map, nonnegative matrix factorization (NMF), Poisson NMF, zero-inflated factor analysis (ZIFA), zero-inflated negative binomial based wanted variation extraction (ZINB-WaVE), probabilistic count matrix factorization (pCMF), deep count autoencoder network (DCA), scScope, generalized linear model principal component analysis (GLMPCA), multidimensional scaling (MDS), locally linear embedding (LLE), local tangent space alignment (LTSA), Isomap, uniform manifold approximation and projection (UMAP), and t-distributed stochastic neighbor embedding (tSNE; dark red). We evaluated their performance on 14 real scRNAseq data sets and 2 simulated data sets (rows). The simulated data based on Kumar data is labeled with #. We used log2 Count transformation for the subset of DR methods that use normalized data. For each data set, we compared the four different number of low-dimensional data. The four numbers we used equal to 0.5%, 1%, 2%, and 3% of the total number of cells in big data and equal to 2, 6, 14, and 20 in small data (which were labeled with *). For convenience, we only listed 0.5%, 1%, 2%, and 3% on x-axis. No results for ICA are shown in the table (grey fills) because ICA cannot handle the large number of features. Note that, for tSNE, we only extracted two low-dimensional components due to the limitation of the tSNE software.

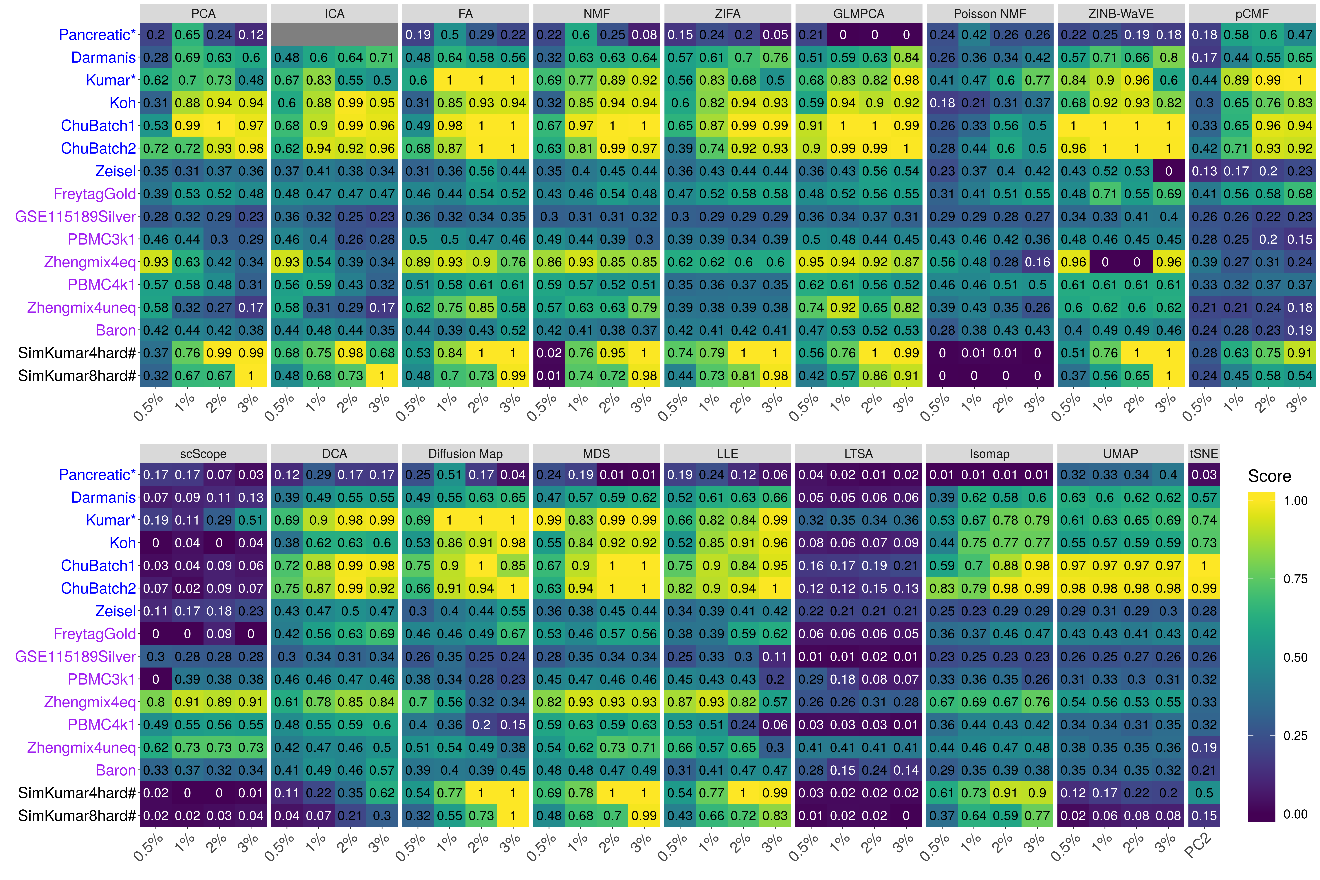

**Supplementary Figure 22. Bar plots show the average performance of different DR methods based on Louvain clustering.** The performance is measured by normalized mutual information (NMI; **A**) or adjusted rand index (ARI; **B**), both averaged across 16 data sets. We compared 18 DR methods, including factor analysis (FA; light green), principal component analysis (PCA; light blue), independent component analysis (ICA; blue), Diffusion Map (pink), nonnegative matrix factorization (NMF; green), Poisson NMF(light orange), zero-inflated factor analysis (ZIFA; light pink), zero-inflated negative binomial based wanted variation extraction (ZINB-WaVE; orange), probabilistic count matrix factorization (pCMF; light purple), deep count autoencoder network (DCA; yellow), scScope, generalized linear model principal component analysis (GLMPCA; red), multidimensional scaling (MDS; cyan), locally linear embedding (LLE; blue green), local tangent space alignment (LTSA; teal blue), Isomap (grey), uniform manifold approximation and projection (UMAP; brown), and t-distributed stochastic neighbor embedding (tSNE; dark red). We used log2 Count transformation for the subset of DR methods that use normalized data. For each data set, we compared the four different number of low-dimensional data. The four numbers we used equal to 0.5%, 1%, 2%, and 3% of the total number of cells in big data and equal to 2, 6, 14, and 20 in small data. For convenience, we only listed 0.5%, 1%, 2%, and 3% on x-axis. Note that, for tSNE, we only extracted two low-dimensional components due to the limitation of the tSNE software.

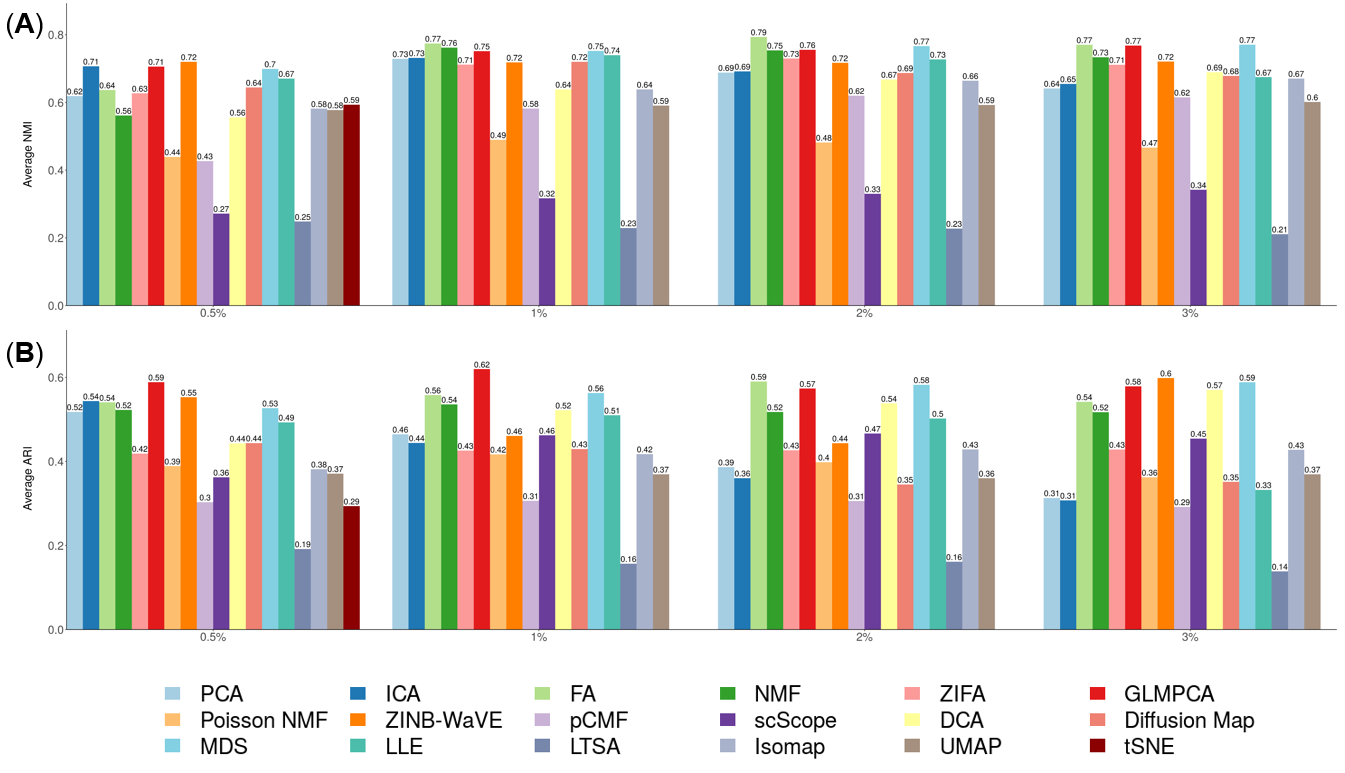

**Supplementary Figure 23. DR method performance with log-2 CPM transformed data evaluated based on downstream cell clustering analysis.** Performance is evaluated by normalized mutual index (NMI) either based on *k*-means clustering. We compared 11 DR methods (columns), including factor analysis (FA), principal component analysis (PCA), independent component analysis (ICA), Diffusion Map, nonnegative matrix factorization (NMF) , multidimensional scaling (MDS), locally linear embedding (LLE), local tangent space alignment (LTSA), Isomap, uniform manifold approximation and projection (UMAP), and t-distributed stochastic neighbor embedding (tSNE). We evaluated their performance on 14 real scRNAseq data sets and 2 simulated data sets (rows). The simulated data based on Kumar data is labeled with #. We used log2 CPM transformation for the subset of DR methods that use normalized data. For each data set, we compared the four different number of low-dimensional data. The four numbers we used equal to 0.5%, 1%, 2%, and 3% of the total number of cells in big data and equal to 2, 6, 14, and 20 in small data (labeled with *). No results for ICA are shown in the Pancreatic data (grey fills) because ICA cannot handle the large number of features in the data. Note that, for tSNE, we only extracted two low-dimensional components due to the limitation of the tSNE software.

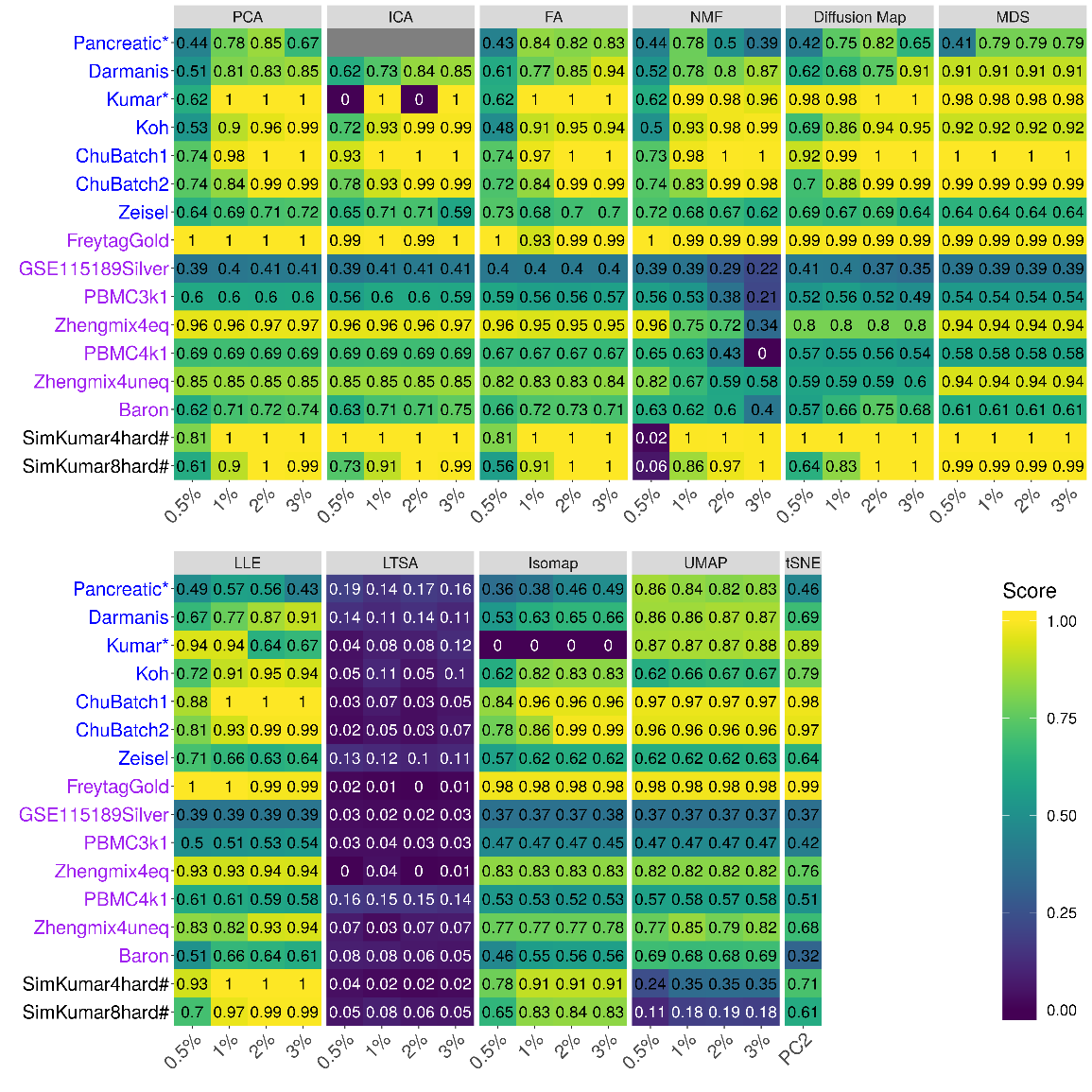

**Supplementary Figure 24. DR method performance with log-2 CPM transformed data evaluated based on downstream cell clustering analysis.** Performance is evaluated by normalized mutual index (NMI) either based on hierarchical clustering. We compared 11 DR methods (columns), including factor analysis (FA), principal component analysis (PCA), independent component analysis (ICA), Diffusion Map, nonnegative matrix factorization (NMF) , multidimensional scaling (MDS), locally linear embedding (LLE), local tangent space alignment (LTSA), Isomap, uniform manifold approximation and projection (UMAP), and t-distributed stochastic neighbor embedding (tSNE). We evaluated their performance on 14 real scRNAseq data sets and 2 simulated data sets (rows). The simulated data based on Kumar data is labeled with #. We used log2 CPM transformation for the subset of DR methods that use normalized data. For each data set, we compared the four different number of low-dimensional data. The four numbers we used equal to 0.5%, 1%, 2%, and 3% of the total number of cells in big data and equal to 2, 6, 14, and 20 in small data (labeled with *). No results for ICA are shown in the Pancreatic data (grey fills) because ICA cannot handle the large number of features in the data. Note that, for tSNE, we only extracted two low-dimensional components due to the limitation of the tSNE software.

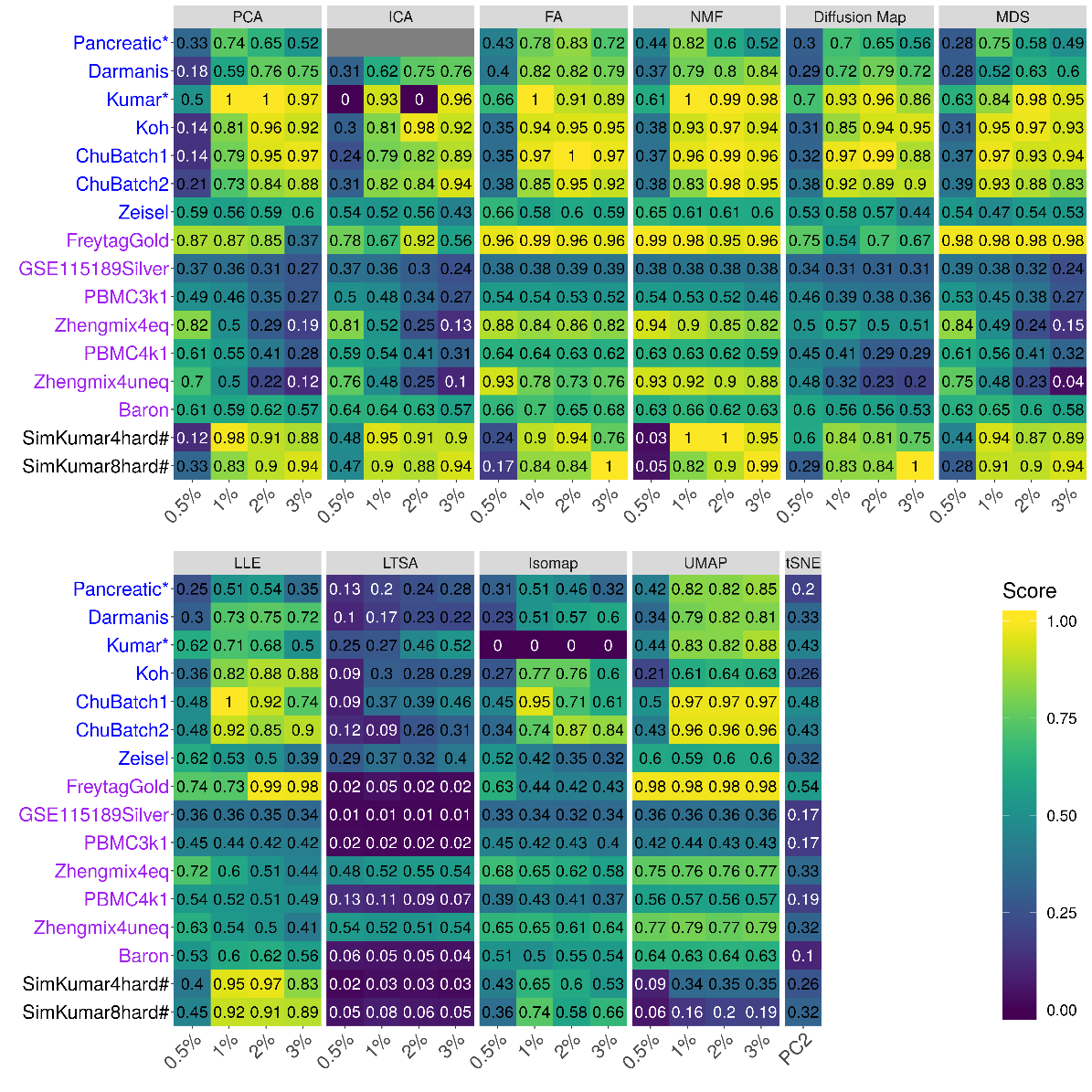

**Supplementary Figure 25. DR method performance with log-2 CPM transformed data evaluated based on downstream cell clustering analysis.** Performance is evaluated by normalized mutual index (NMI) either based on Louvain method. We compared 11 DR methods (columns), including factor analysis (FA), principal component analysis (PCA), independent component analysis (ICA), Diffusion Map, nonnegative matrix factorization (NMF), multidimensional scaling (MDS), locally linear embedding (LLE), local tangent space alignment (LTSA), Isomap, uniform manifold approximation and projection (UMAP), and t-distributed stochastic neighbor embedding (tSNE). We evaluated their performance on 14 real scRNAseq data sets and 2 simulated data sets (rows). The simulated data based on Kumar data is labeled with #. We used log2 CPM transformation for the subset of DR methods that use normalized data. For each data set, we compared the four different number of low-dimensional data. The four numbers we used equal to 0.5%, 1%, 2%, and 3% of the total number of cells in big data and equal to 2, 6, 14, and 20 in small data (labeled with *). No results for ICA are shown in the Pancreatic data (zeros) because ICA cannot handle the large number of features in the data. Note that, for tSNE, we only extracted two low-dimensional components due to the limitation of the tSNE software.

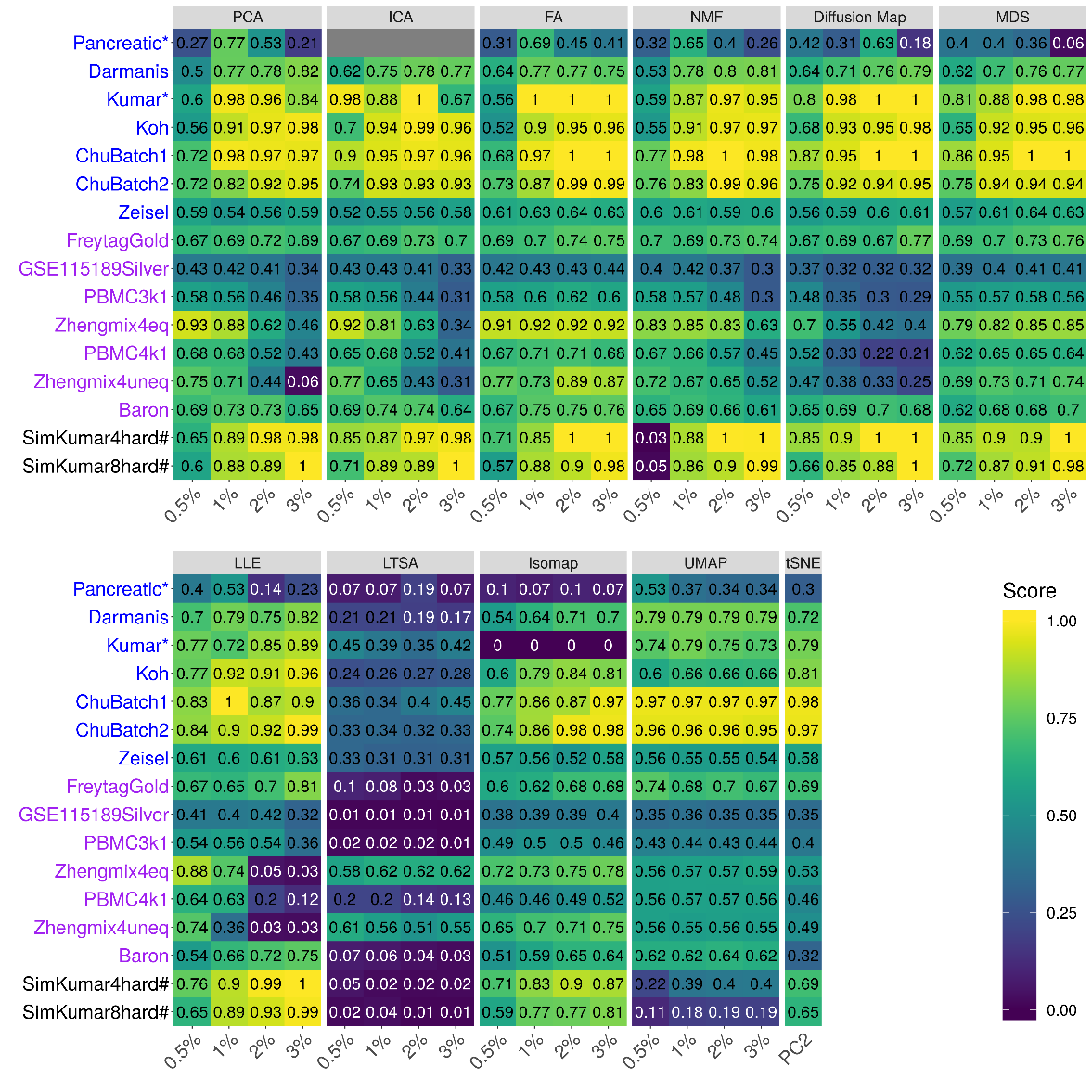

**Supplementary Figure 26. Bar plots show the average performance of a subset of DR methods that use log2 CPM normalized data.** The performance of each DR method is measured by normalized mutual index (NMI) using either *k*-means clustering (**A**), hierarchical clustering (**B**) or Louvain method (**C**), the averaged across 16 date sets. We used log2 CPM transformation for the subset of DR methods that use normalized data. We compared 11 DR methods that use normalized data, including factor analysis (FA; light green), principal component analysis (PCA; light blue), independent component analysis (ICA; blue), Diffusion Map (pink), nonnegative matrix factorization (NMF; green), multidimensional scaling (MDS; cyan), locally linear embedding (LLE; blue green), local tangent space alignment (LTSA; teal blue), Isomap (grey), uniform manifold approximation and projection (UMAP; brown), and t-distributed stochastic neighbor embedding (tSNE; dark red). For each data set, we compared the four different number of low-dimensional data. The four numbers we used equal to 0.5%, 1%, 2%, and 3% of the total number of cells in big data and equal to 2, 6, 14, and 20 in small data. For convenience, we only listed 0.5%, 1%, 2%, and 3% on x-axis. Note that, for tSNE, we only extracted two low-dimensional components due to the limitation of the tSNE software.

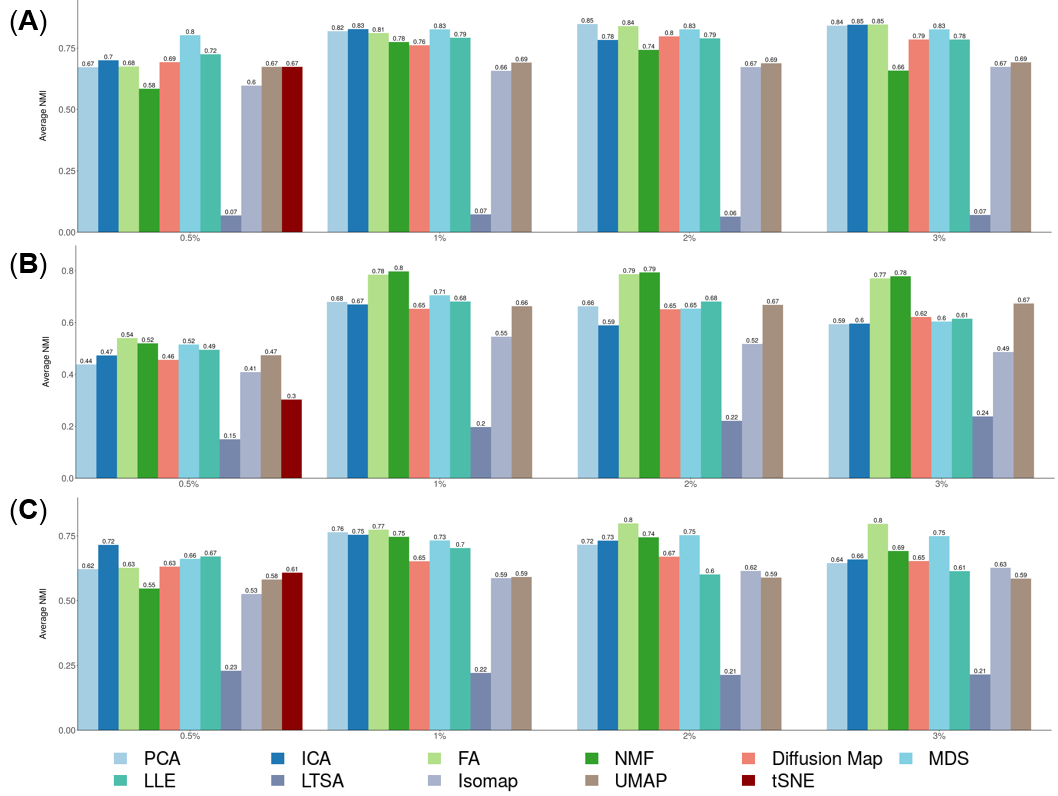

**Supplementary Figure 27. DR method performance with z-score transformed data in downstream cell clustering analysis.** Performance is evaluated by normalized mutual index (NMI) either based on *k*-means clustering. We compared 10 DR methods (columns), including factor analysis (FA), principal component analysis (PCA), independent component analysis (ICA), Diffusion Map, multidimensional scaling (MDS), locally linear embedding (LLE), local tangent space alignment (LTSA), Isomap, uniform manifold approximation and projection (UMAP), and t-distributed stochastic neighbor embedding (tSNE). We evaluated their performance on 14 real scRNAseq data sets and 2 simulated data sets (rows). The simulated data based on Kumar data is labeled with #. We used z-score transformation for the subset of DR methods that use normalized data. For each data set, we compared the four different number of low-dimensional data. The four numbers we used equal to 0.5%, 1%, 2%, and 3% of the total number of cells in big data and equal to 2, 6, 14, and 20 in small data which were labeled with *. For convenience, we only listed 0.5%, 1%, 2%, and 3% on x-axis. No results for ICA are shown in the Pancreatic data (grey fills) because ICA cannot handle the large number of features in the data. No results for NMF are shown because z-score normalization have negative values. Note that, for tSNE, we only extracted two low-dimensional components due to the limitation of the tSNE software.

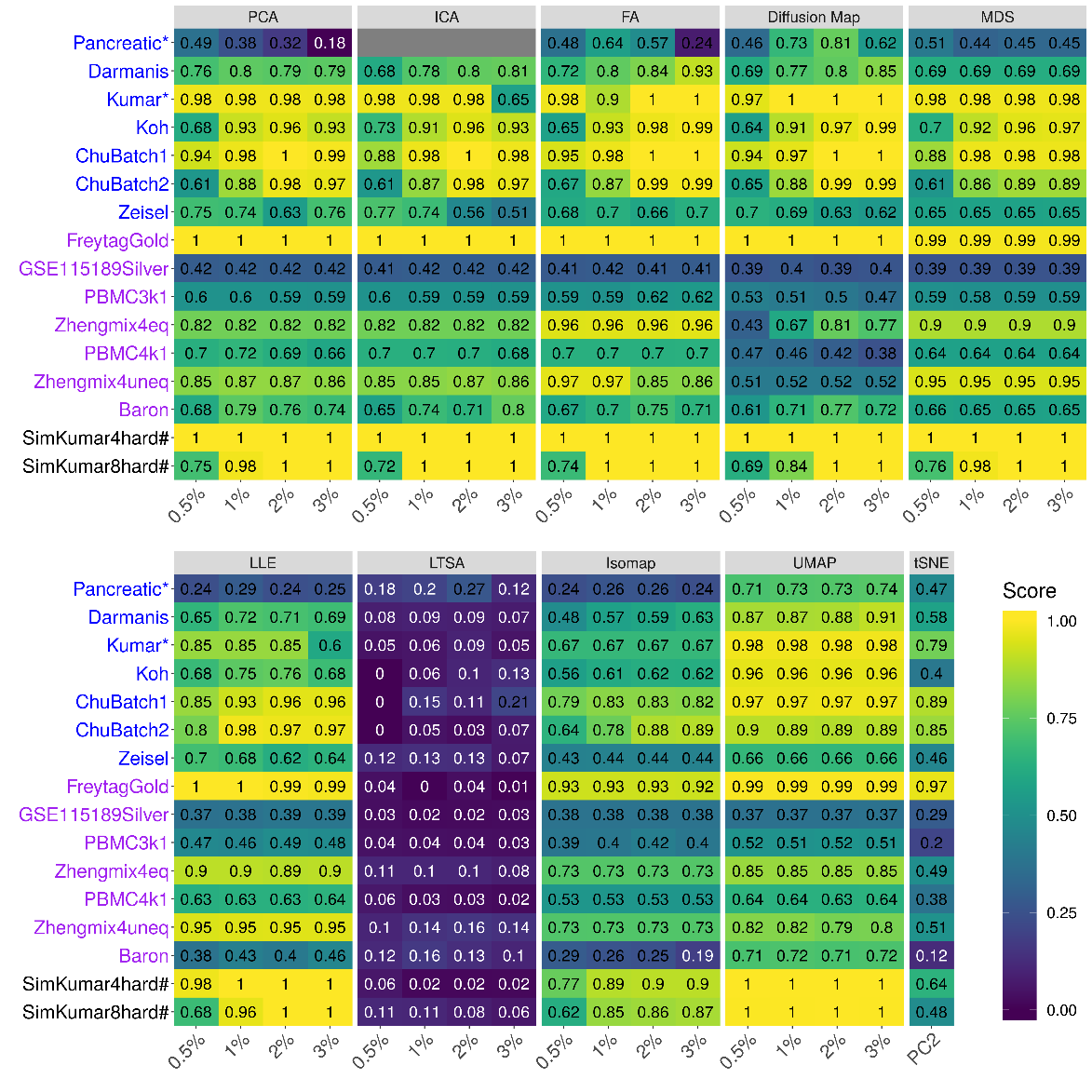

**Supplementary Figure 28. DR method performance with z-score transformed data in downstream cell clustering analysis.** Performance is evaluated by normalized mutual index (NMI) either based on hierarchical clustering. We compared 10 DR methods (columns), including factor analysis (FA), principal component analysis (PCA), independent component analysis (ICA), Diffusion Map, multidimensional scaling (MDS), locally linear embedding (LLE), local tangent space alignment (LTSA), Isomap, uniform manifold approximation and projection (UMAP), and t-distributed stochastic neighbor embedding (tSNE). We evaluated their performance on 14 real scRNAseq data sets and 2 simulated data sets (rows). The simulated data based on Kumar data is labeled with #. We used z-score transformation for the subset of DR methods that use normalized data. For each data set, we compared the four different number of low-dimensional data. The four numbers we used equal to 0.5%, 1%, 2%, and 3% of the total number of cells in big data and equal to 2, 6, 14, and 20 in small data which were labeled with *. For convenience, we only listed 0.5%, 1%, 2%, and 3% on x-axis. No results for ICA are shown in the Pancreatic data (grey fills) because ICA cannot handle the large number of features in the data. No results for NMF are shown because z-score normalization have negative values. Note that, for tSNE, we only extracted two low-dimensional components due to the limitation of the tSNE software.

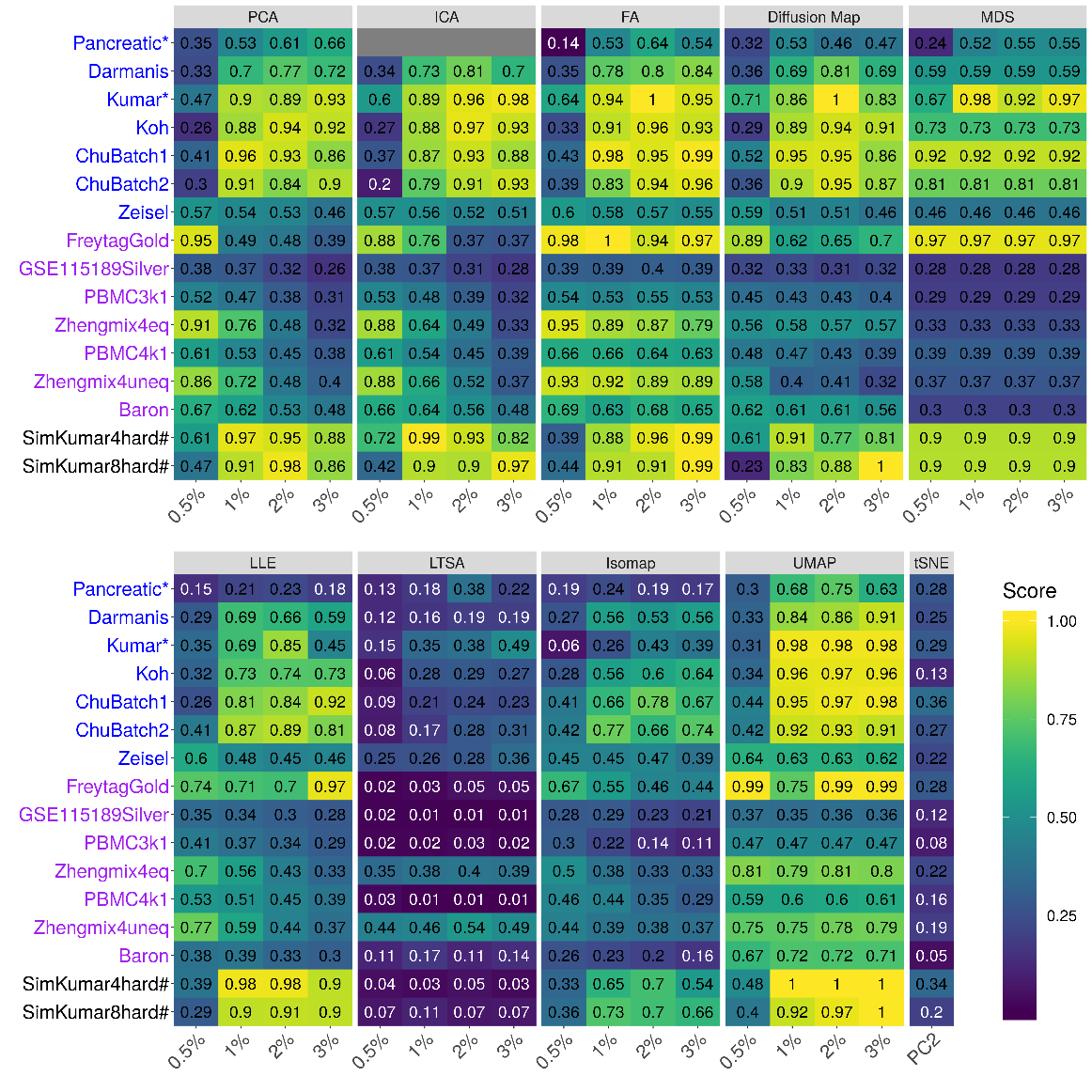

**Supplementary Figure 29. DR method performance with z-score transformed data in downstream cell clustering analysis.** Performance is evaluated by normalized mutual index (NMI) either based on Louvain method. We compared 10 DR methods (columns), including factor analysis (FA), principal component analysis (PCA), independent component analysis (ICA), Diffusion Map, multidimensional scaling (MDS), locally linear embedding (LLE), local tangent space alignment (LTSA), Isomap, uniform manifold approximation and projection (UMAP), and t-distributed stochastic neighbor embedding (tSNE). We evaluated their performance on 14 real scRNAseq data sets and 2 simulated data sets (rows). The simulated data based on Kumar data is labeled with #. We used z-score transformation for the subset of DR methods that use normalized data. For each data set, we compared the four different number of low-dimensional data. The four numbers we used equal to 0.5%, 1%, 2%, and 3% of the total number of cells in big data and equal to 2, 6, 14, and 20 in small data which were labeled with *. For convenience, we only listed 0.5%, 1%, 2%, and 3% on x-axis. No results for ICA are shown in the Pancreatic data (grey fills) because ICA cannot handle the large number of features in the data. No results for NMF are shown because z-score normalization have negative values. Note that, for tSNE, we only extracted two low-dimensional components due to the limitation of the tSNE software.

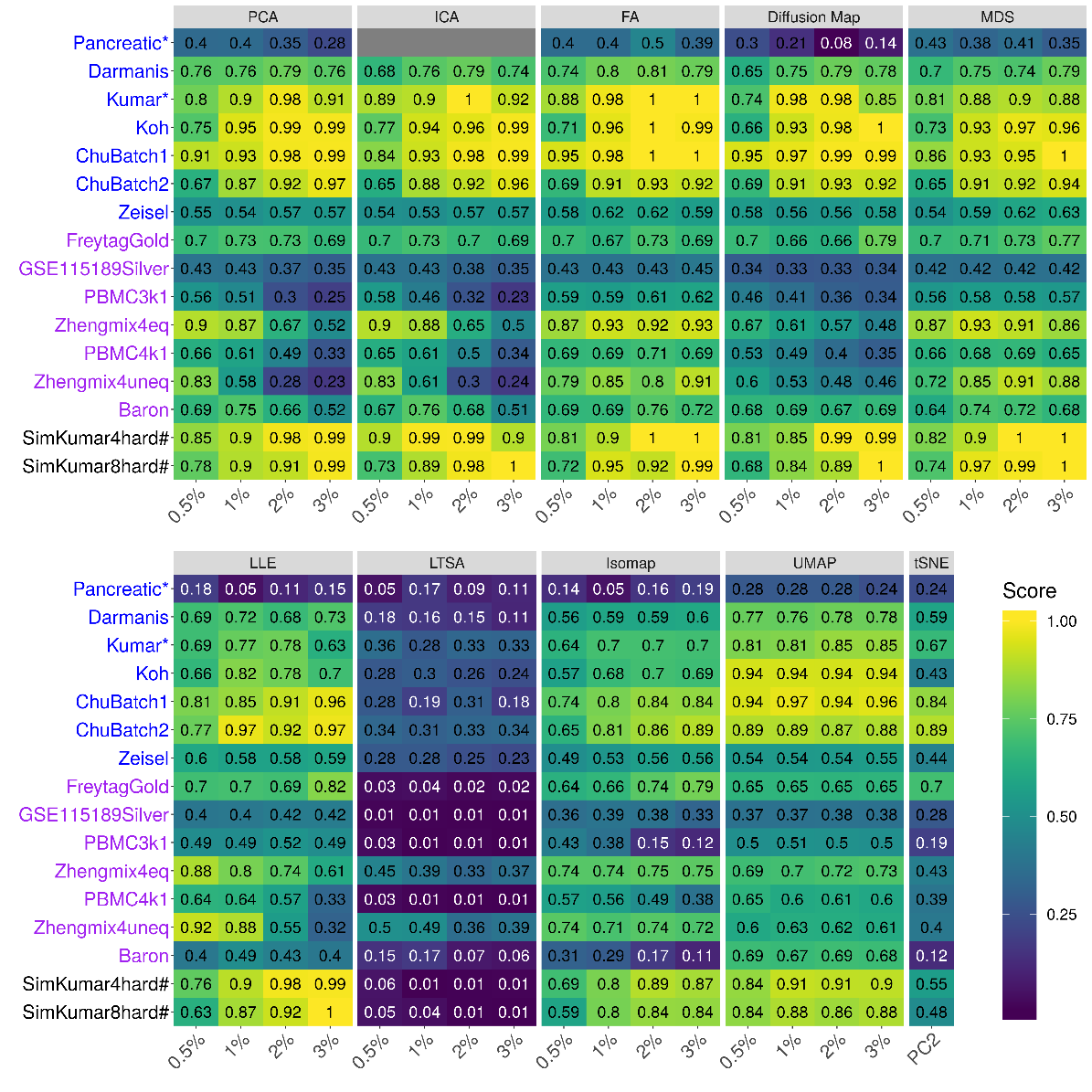

**Supplementary Figure 30. Bar plots show the average performance of a subset of DR methods that use z-score normalized data.** The performance of each DR method is measured by normalized mutual index (NMI) using either *k*-means clustering (**A**), hierarchical clustering (**B**) or Louvain method (**C**), the averaged across 16 date sets. We used z-score transformation for the subset of DR methods that use normalized data. We compared 10 DR methods that use normalized data, including factor analysis (FA; light green), principal component analysis (PCA; light blue), independent component analysis (ICA; blue), Diffusion Map (pink), multidimensional scaling (MDS; cyan), locally linear embedding (LLE; blue green), local tangent space alignment (LTSA; teal blue), Isomap (grey), uniform manifold approximation and projection (UMAP; brown), and t-distributed stochastic neighbor embedding (tSNE; dark red). For each data set, we compared the four different number of low-dimensional data. The four numbers we used equal to 0.5%, 1%, 2%, and 3% of the total number of cells in big data and equal to 2, 6, 14, and 20 in small data. For convenience, we only listed 0.5%, 1%, 2%, and 3% on x-axis. The four numbers we used equal to 0.5%, 1%, 2%, and 3% of the total number of cells in big data and equal to 2, 6, 14, and 20 in small data. For convenience, we only listed 0.5%, 1%, 2%, and 3% on x-axis. Note that, for tSNE, we only extracted two low-dimensional components due to the limitation of the tSNE software.

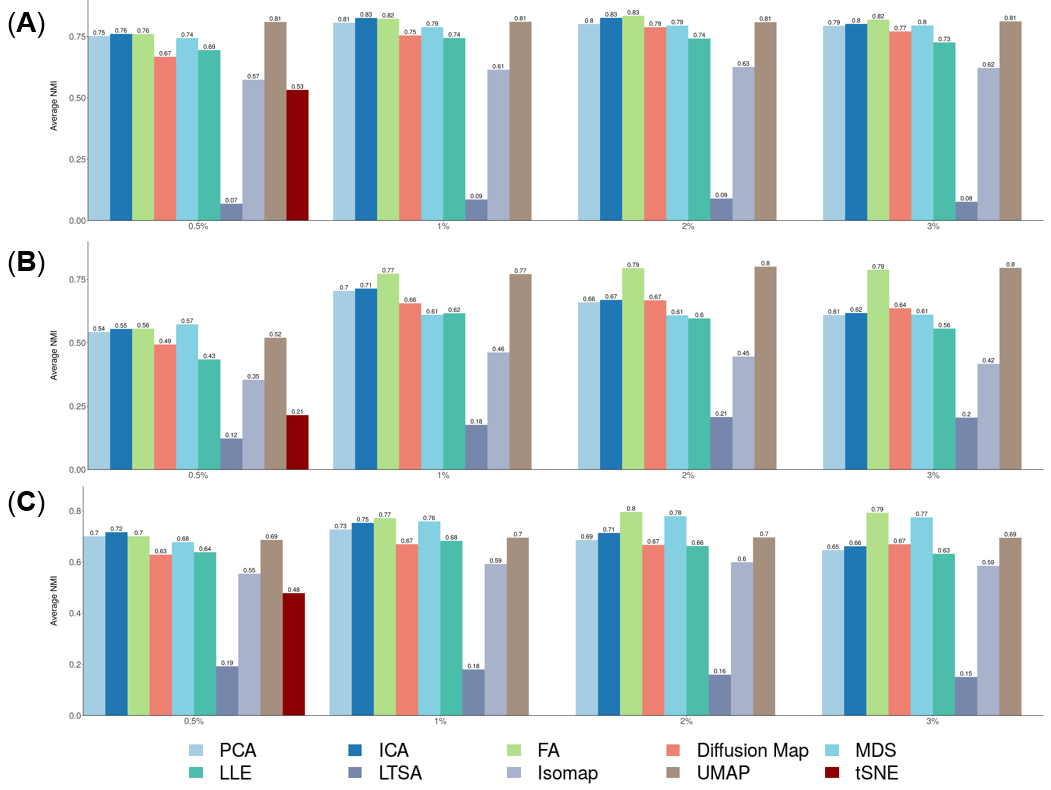

**Supplementary Figure 31. Bar plots show the average NMI of different DR methods on UMI-based data and nonUMI-based data.** The performance is measured by average NMI across UMI-based data (**A**; 7 data sets) or nonUMI-based data (**B**; 7 data sets). We compared 18 DR methods, including factor analysis (FA; light green), principal component analysis (PCA; light blue), independent component analysis (ICA; blue), Diffusion Map (pink), nonnegative matrix factorization (NMF; green), Poisson NMF(light orange), zero-inflated factor analysis (ZIFA; light pink), zero-inflated negative binomial based wanted variation extraction (ZINB-WaVE; orange), probabilistic count matrix factorization (pCMF; light purple), deep count autoencoder network (DCA; yellow), scScope (purple), generalized linear model principal component analysis (GLMPCA; red), multidimensional scaling (MDS; cyan), locally linear embedding (LLE; blue green), local tangent space alignment (LTSA; teal blue), Isomap (grey), uniform manifold approximation and projection (UMAP; brown), and t-distributed stochastic neighbor embedding (tSNE; dark red). We used log2 count transformation for the subset of DR methods that use normalized data. For each data set, we compared the four different number of low-dimensional data. The four numbers we used equal to 0.5%, 1%, 2%, and 3% of the total number of cells in big data and equal to 2, 6, 14, and 20 in small data. For convenience, we only listed 0.5%, 1%, 2%, and 3% on x-axis. Note that, for tSNE, we only extracted two low-dimensional components due to the limitation of the tSNE software.

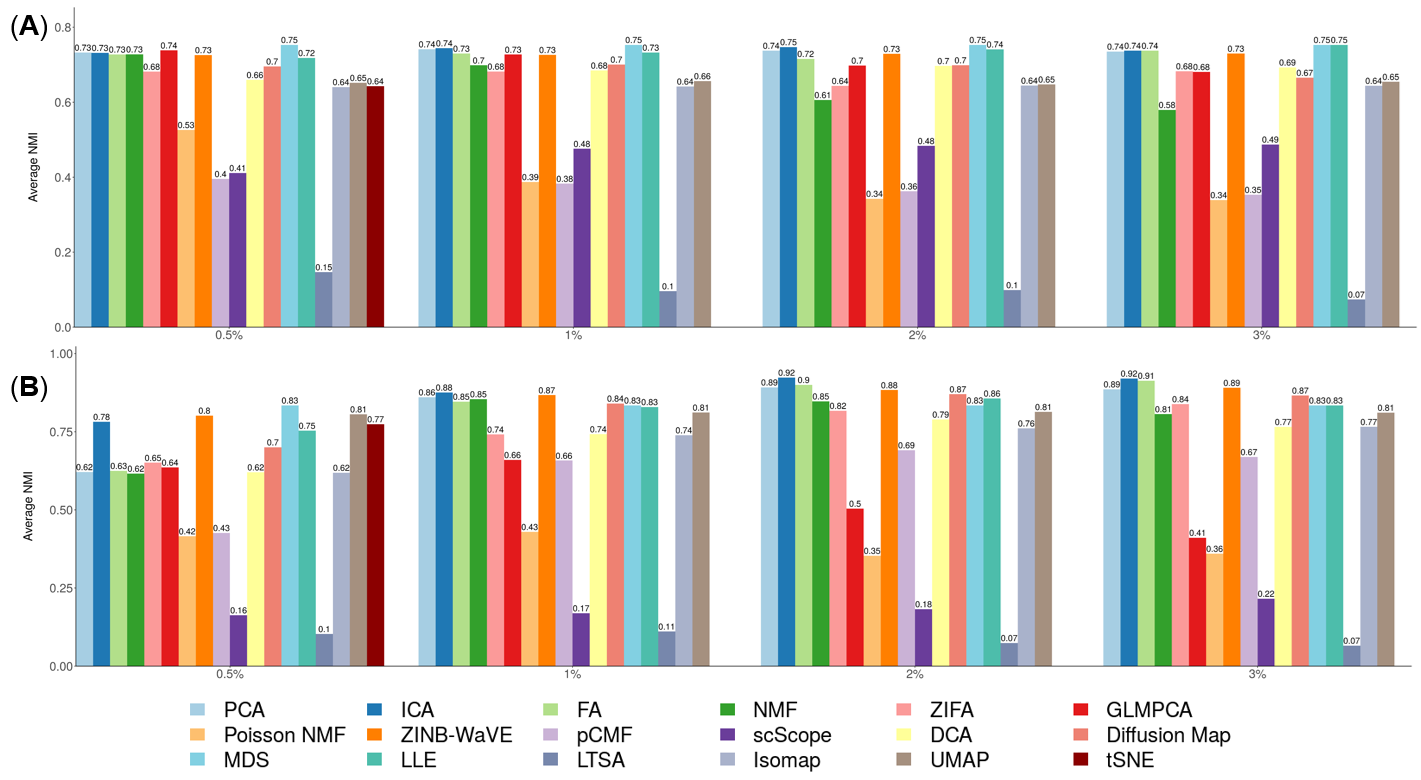

**Supplementary Figure 32. Scatter plots visualize the clusters of different DR methods on Kumar data set.** We compared 17 DR methods (columns), including factor analysis (FA), principal component analysis (PCA), independent component analysis (ICA), Diffusion Map, nonnegative matrix factorization (NMF), Poisson NMF, zero-inflated factor analysis (ZIFA), zero-inflated negative binomial based wanted variation extraction (ZINB-WaVE), probabilistic count matrix factorization (pCMF), deep count autoencoder network (DCA), scScope, generalized linear model principal component analysis (GLMPCA), multidimensional scaling (MDS), locally linear embedding (LLE) , Isomap, uniform manifold approximation and projection (UMAP), and t-distributed stochastic neighbor embedding (tSNE). We used log2 count transformation for the subset of DR methods that use normalized data. For each data set, we first reduced the dimension to 20, then applied tSNE to visualize the clusters. Local tangent space alignment (LTSA) was excluded because of failing to extract low-dimensional components.

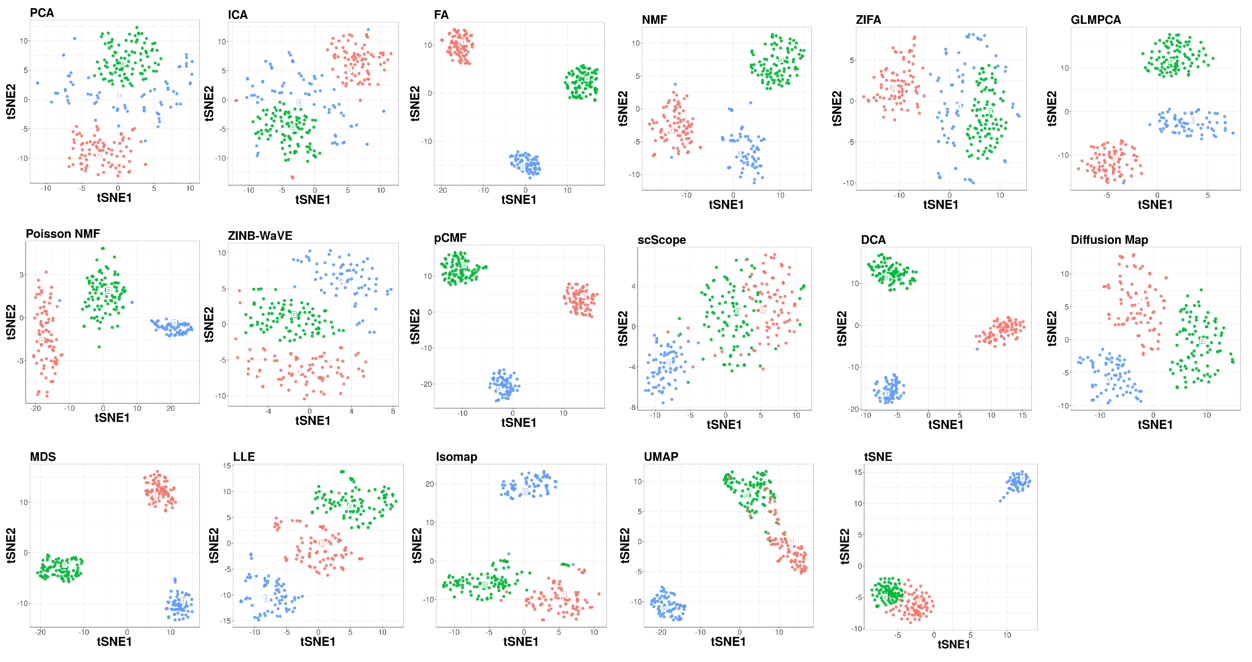

**Supplementary Figure 33. Scatter plots show the clusters of different DR methods on PBMC3k data set.** We compared 18 DR methods (columns), including factor analysis (FA), principal component analysis (PCA), independent component analysis (ICA), Diffusion Map, nonnegative matrix factorization (NMF), Poisson NMF, zero-inflated factor analysis (ZIFA), zero-inflated negative binomial based wanted variation extraction (ZINB-WaVE), probabilistic count matrix factorization (pCMF), deep count autoencoder network (DCA), scScope, generalized linear model principal component analysis (GLMPCA), multidimensional scaling (MDS), locally linear embedding (LLE), local tangent space alignment (LTSA), Isomap, uniform manifold approximation and projection (UMAP), and t-distributed stochastic neighbor embedding (tSNE). We used log2 count transformation for the subset of DR methods that use normalized data. For each data set, we first reduced the dimension to 32, then applied tSNE to visualize the clusters.

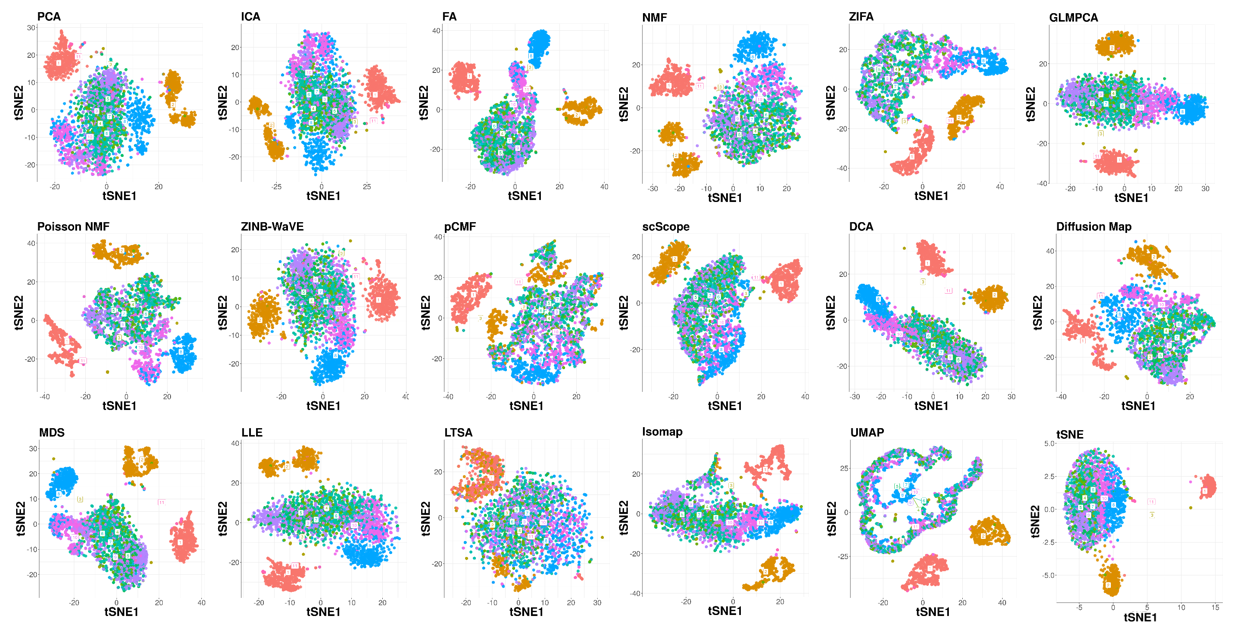

**Supplementary Figure 34. Bar plots show the F-measure of different DR methods on the rare data set PBMC3k1Rare1 (contains 4.0% CD34+ cells).** The performance is measured by *k*-means clustering (**A**), hierarchical clustering (**B**) or Louvain method (**C**). We compared 18 DR methods, including factor analysis (FA; light green), principal component analysis (PCA; light blue), independent component analysis (ICA; blue), Diffusion Map (pink), nonnegative matrix factorization (NMF; green), Poisson NMF(light orange), zero-inflated factor analysis (ZIFA; light pink), zero-inflated negative binomial based wanted variation extraction (ZINB-WaVE; orange), probabilistic count matrix factorization (pCMF; light purple), deep count autoencoder network (DCA; yellow), scScope (purple), generalized linear model principal component analysis (GLMPCA; red), multidimensional scaling (MDS; cyan), locally linear embedding (LLE; blue green), local tangent space alignment (LTSA; teal blue), Isomap (grey), uniform manifold approximation and projection (UMAP; brown), and t-distributed stochastic neighbor embedding (tSNE; dark red). We used log2 count transformation for the subset of DR methods that use normalized data. For each data set, we compared the four different number of low-dimensional data. The four numbers we used equal to 0.5%, 1%, 2%, and 3% of the total number of cells. Note that, for tSNE, we only extracted two low-dimensional components due to the limitation of the tSNE software.

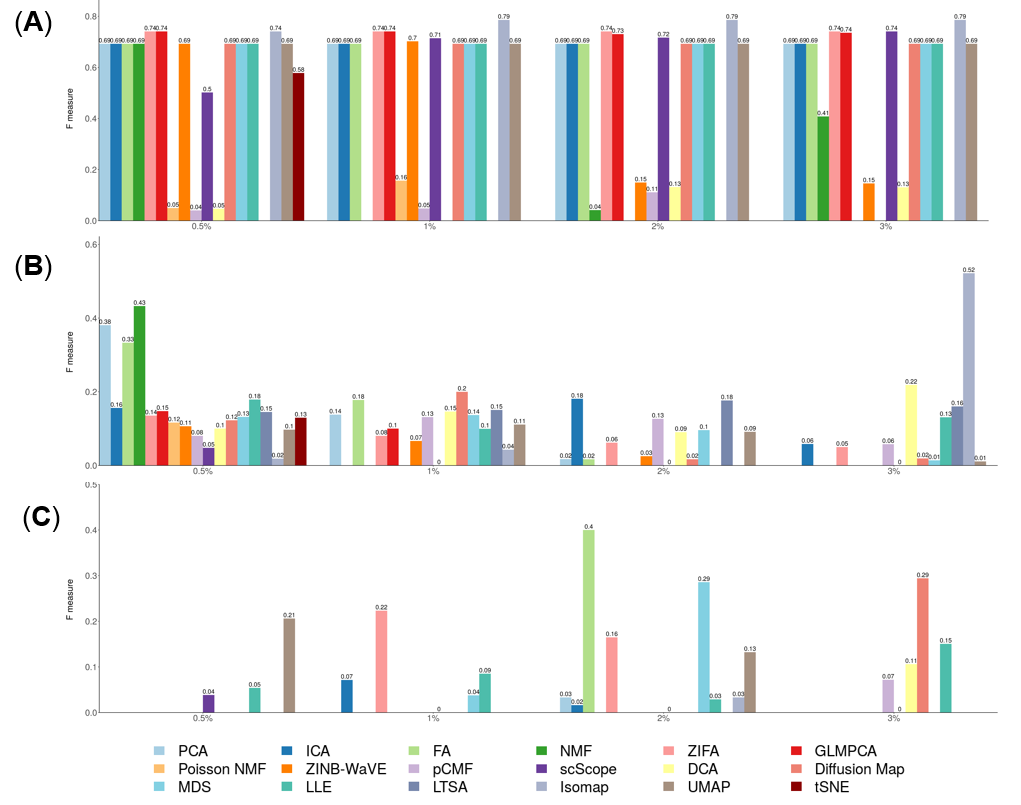

**Supplementary Figure 35. Bar plots show the F-measure of different DR methods on the rare data set PBMC3k1Rare2 (contains 7.9% CD34+ cells).** The performance is measured by k-means clustering (**A**), hierarchical clustering (**B**) or Louvain method (**C**). We compared 18 DR methods, including factor analysis (FA; light green), principal component analysis (PCA; light blue), independent component analysis (ICA; blue), Diffusion Map (pink), nonnegative matrix factorization (NMF; green), Poisson NMF(light orange), zero-inflated factor analysis (ZIFA; light pink), zero-inflated negative binomial based wanted variation extraction (ZINB-WaVE; orange), probabilistic count matrix factorization (pCMF; light purple), deep count autoencoder network (DCA; yellow), scScope (purple), generalized linear model principal component analysis (GLMPCA; red), multidimensional scaling (MDS; cyan), locally linear embedding (LLE; blue green), local tangent space alignment (LTSA; teal blue), Isomap (grey), uniform manifold approximation and projection (UMAP; brown), and t-distributed stochastic neighbor embedding (tSNE; dark red). We used log2 count transformation for the subset of DR methods that use normalized data. For each data set, we compared the four different number of low-dimensional data. The four numbers we used equal to 0.5%, 1%, 2%, and 3% of the total number of cells. Note that, for tSNE, we only extracted two low-dimensional components due to the limitation of the tSNE software.

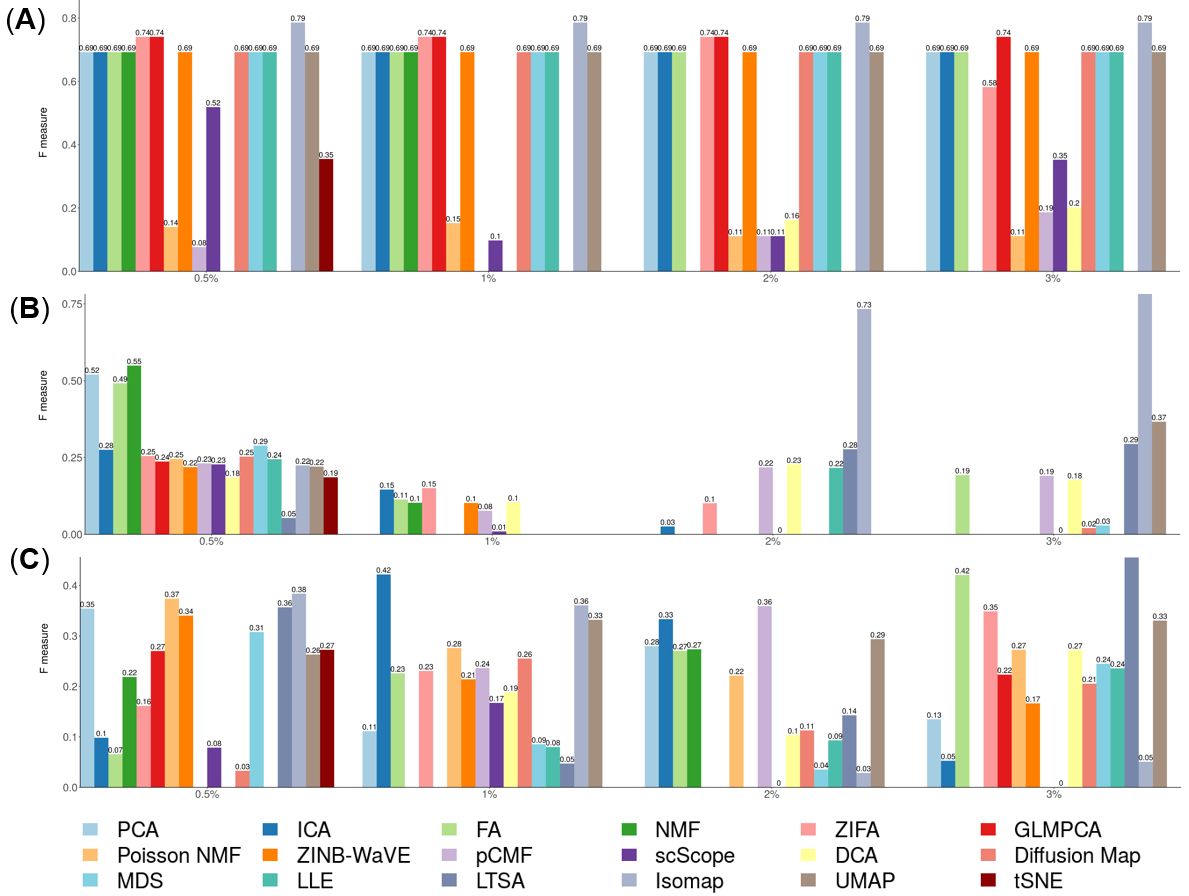

**Supplementary Figure 36. The stability and robustness of DR methods evaluated based on data split of the Kumar data.** We compared 18 DR methods (columns), including factor analysis (FA), principal component analysis (PCA), independent component analysis (ICA), Diffusion Map, nonnegative matrix factorization (NMF), Poisson NMF, zero-inflated factor analysis (ZIFA), zero-inflated negative binomial based wanted variation extraction (ZINB-WaVE), probabilistic count matrix factorization (pCMF), deep count autoencoder network (DCA), scScope, generalized linear model principal component analysis (GLMPCA), multidimensional scaling (MDS), locally linear embedding (LLE), local tangent space alignment (LTSA), Isomap, uniform manifold approximation and projection (UMAP), and t-distributed stochastic neighbor embedding (tSNE). We split the Kumar data randomly into two subsets with equal number of cells in each cell type (one colored red; the other colored green). We applied each DR method to each subset separately to obtain clustering results. Performance is measured by normalized mutual index (NMI) after *k*-means clustering. We performed 10 replicates for each data and show the performance results across these replicates using boxplot. For each data set, we compared the four different number of low-dimensional data. The four numbers we used equal to 2, 6, 14, and 20 (x-axis). Note that, for tSNE, we only extracted two low-dimensional components due to the limitation of the tSNE software.

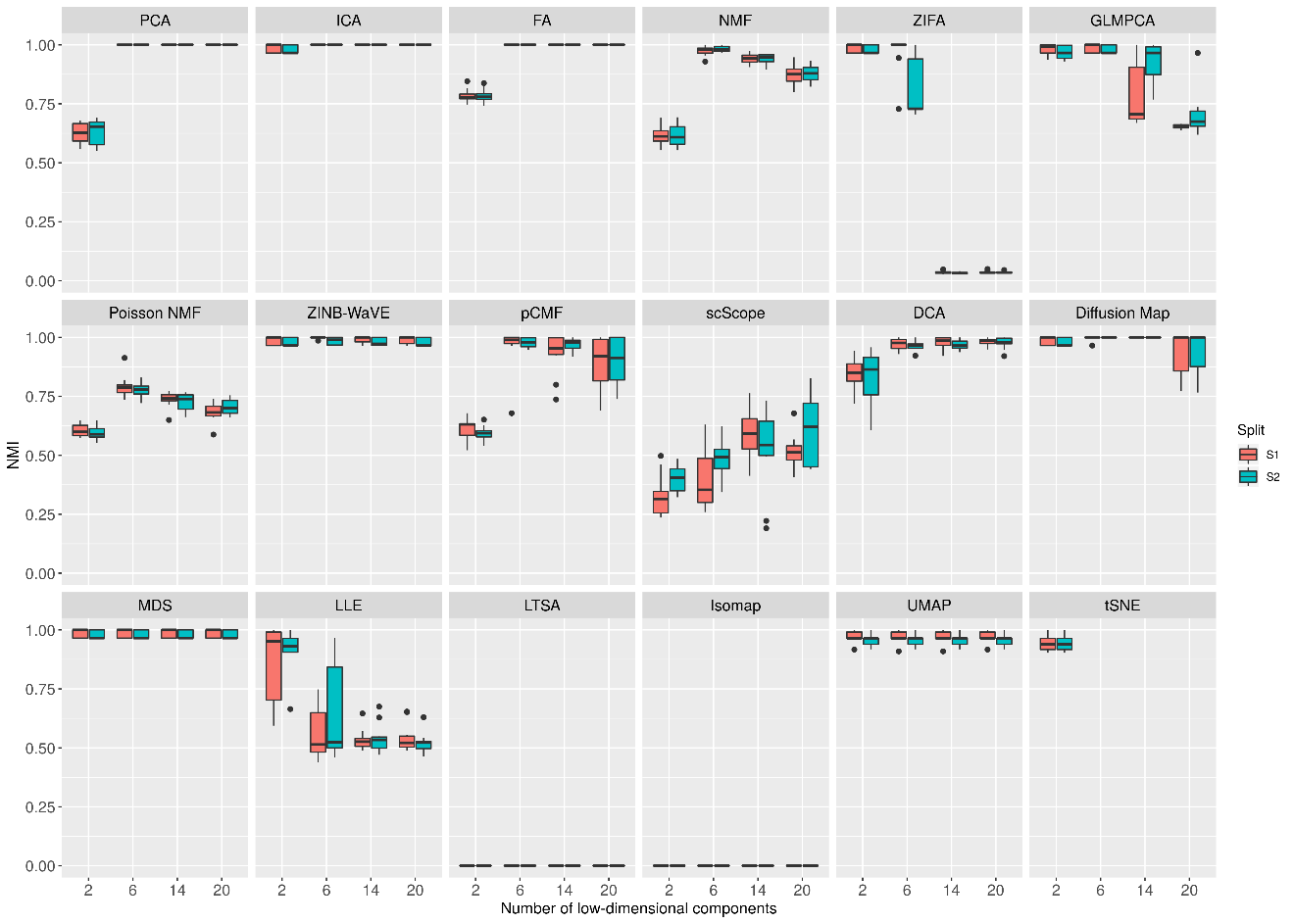

**Supplementary Figure 37. DR method performance evaluated by Kendal correlation for downstream trajectory inference analysis.** For each method, we used *Slingshot* to perform trajectory inference with hierarchical clustering as the initial step. We compared 17 DR methods (columns), including factor analysis (FA), principal component analysis (PCA), independent component analysis (ICA), Diffusion Map, nonnegative matrix factorization (NMF), Poisson NMF, zero-inflated factor analysis (ZIFA), zero-inflated negative binomial based wanted variation extraction (ZINB-WaVE), probabilistic count matrix factorization (pCMF), deep count autoencoder network (DCA), generalized linear model principal component analysis (GLMPCA), multidimensional scaling (MDS), locally linear embedding (LLE), local tangent space alignment (LTSA), Isomap, uniform manifold approximation and projection (UMAP), and t-distributed stochastic neighbor embedding (tSNE). We evaluated their performance on 14 real scRNAseq data sets. We used log2 Count transformation for the subset of DR methods that use normalized data. For each data set, we compared the four different number of low-dimensional data. The four numbers we used equal to 2, 6, 14, and 20 (x-axis). Some results of LTSA are empty because it is not compatible with *Slingshot.* Note that, for tSNE, we only extracted two low-dimensional components due to the limitation of the tSNE software.

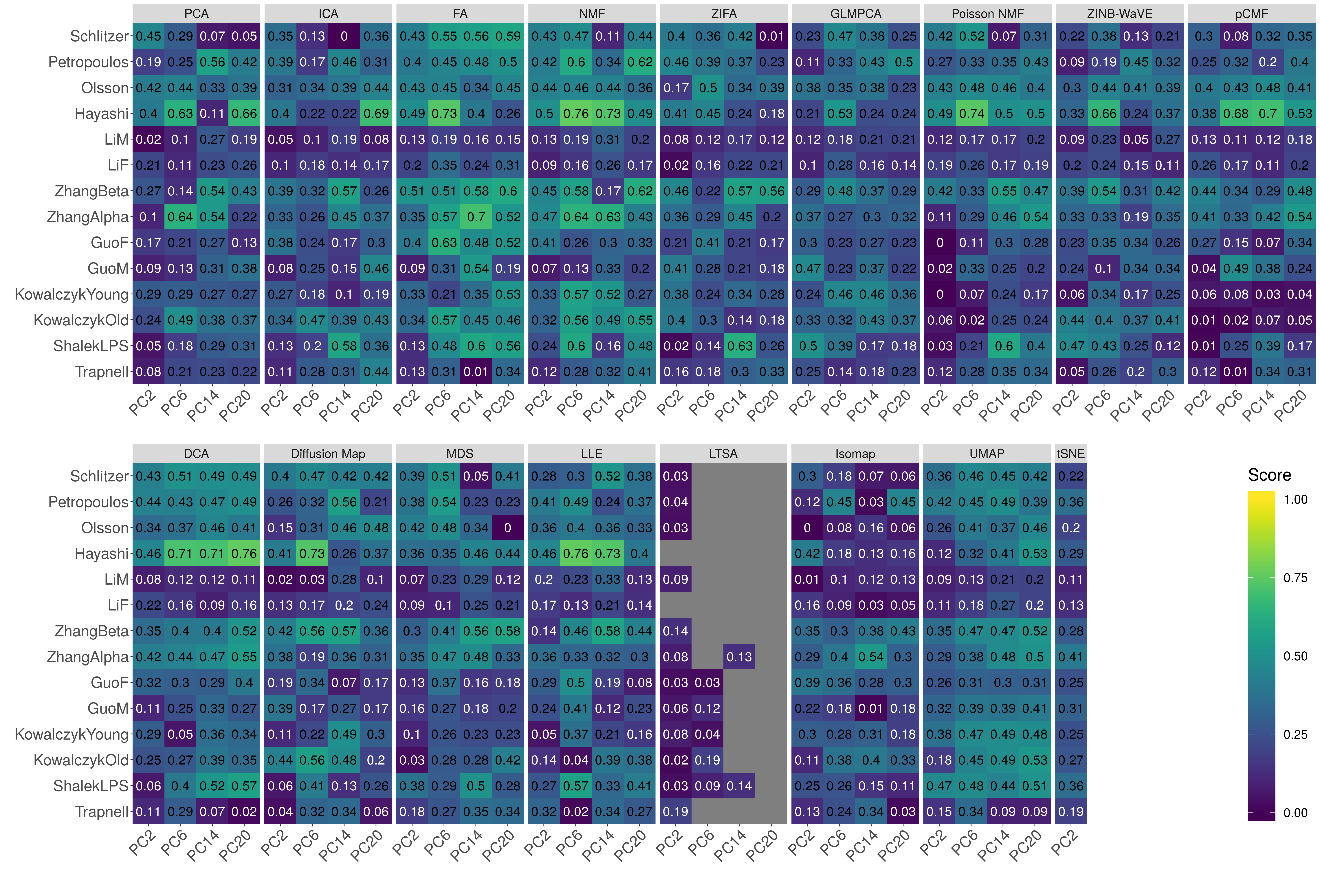

**Supplementary Figure 38. DR method performance evaluated by Kendal correlation for downstream trajectory inference analysis.** For each method, we used *Slingshot* to perform trajectory inference with Louvain method as the initial step. We compared 17 DR methods (columns), including factor analysis (FA), principal component analysis (PCA), independent component analysis (ICA), Diffusion Map, nonnegative matrix factorization (NMF), Poisson NMF, zero-inflated factor analysis (ZIFA), zero-inflated negative binomial based wanted variation extraction (ZINB-WaVE), probabilistic count matrix factorization (pCMF), deep count autoencoder network (DCA), generalized linear model principal component analysis (GLMPCA), multidimensional scaling (MDS), locally linear embedding (LLE), local tangent space alignment (LTSA), Isomap, uniform manifold approximation and projection (UMAP), and t-distributed stochastic neighbor embedding (tSNE). We evaluated their performance on 14 real scRNAseq data sets. We used log2 Count transformation for the subset of DR methods that use normalized data. For each data set, we compared the four different number of low-dimensional data. The four numbers we used equal to 2, 6, 14, and 20 (x-axis). Some results of LTSA are empty because it is not compatible with *Slingshot.* Note that, for tSNE, we only extracted two low-dimensional components due to the limitation of the tSNE software.

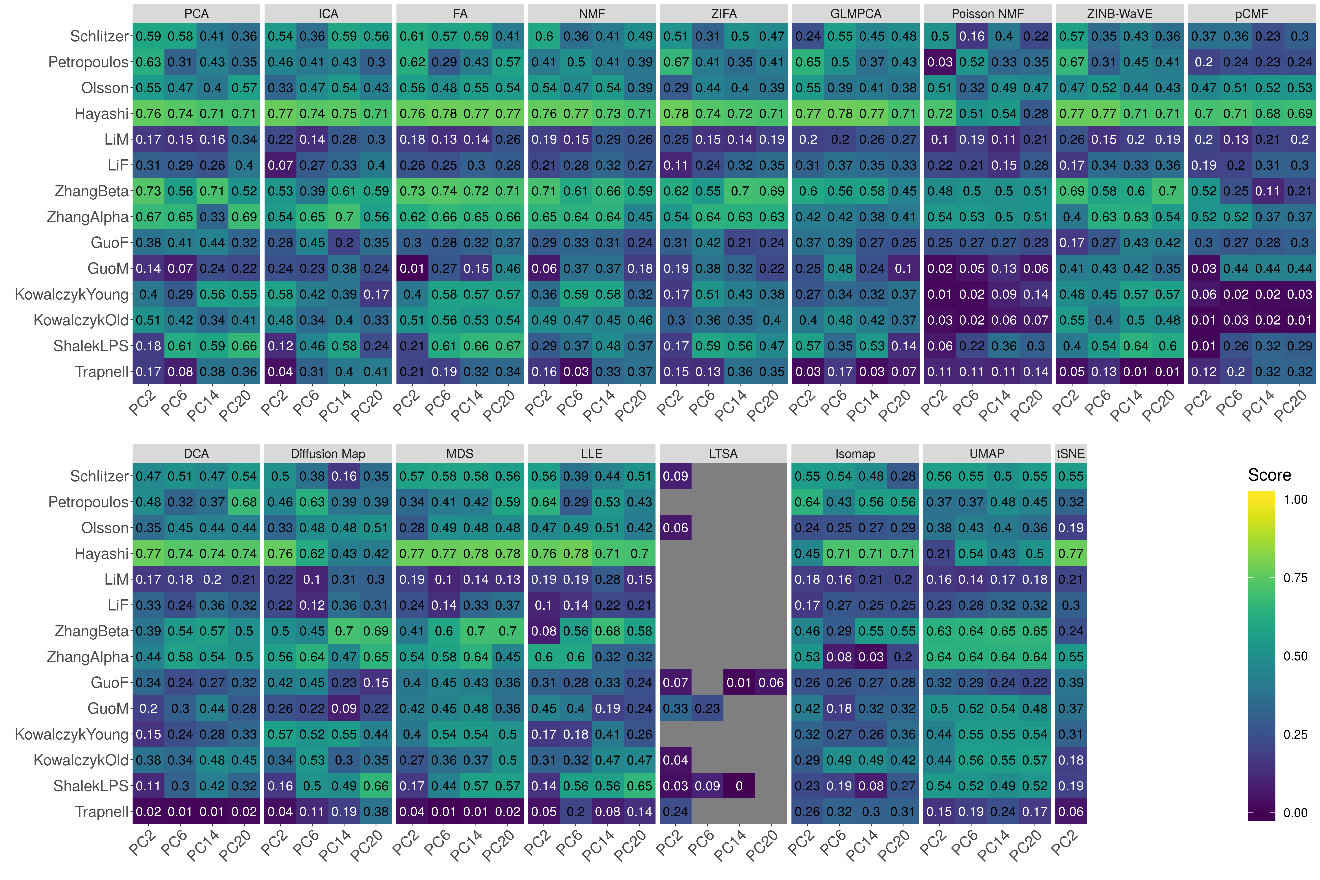

**Supplementary Figure 39. Bar plots show the average performance of different DR methods in downstream trajectory inference analysis.** The performance is evaluated by Kendal correlation averaged across 14 data sets. For each method, we used *Slingshot* to perform trajectory inference with either *k*-means (**A**), hierarchical clustering (**B**) or Louvain method (**C**) as the initial step. We compared 17 DR methods, including factor analysis (FA; light green), principal component analysis (PCA; light blue), independent component analysis (ICA; blue), Diffusion Map (pink), nonnegative matrix factorization (NMF; green), Poisson NMF(light orange), zero-inflated factor analysis (ZIFA; light pink), zero-inflated negative binomial based wanted variation extraction (ZINB-WaVE; orange), probabilistic count matrix factorization (pCMF; light purple), deep count autoencoder network (DCA; yellow), generalized linear model principal component analysis (GLMPCA; red), multidimensional scaling (MDS; cyan), locally linear embedding (LLE; blue green), local tangent space alignment (LTSA; teal blue), Isomap (grey), uniform manifold approximation and projection (UMAP; brown), and t-distributed stochastic neighbor embedding (tSNE; dark red). We used log2 Count transformation for the subset of DR methods that use normalized data. For each data set, we compared the four different number of low-dimensional data. The four numbers we used equal to 2, 6, 14, and 20 (x-axis). Note that, for tSNE, we only extracted two low-dimensional components due to the limitation of the tSNE software.

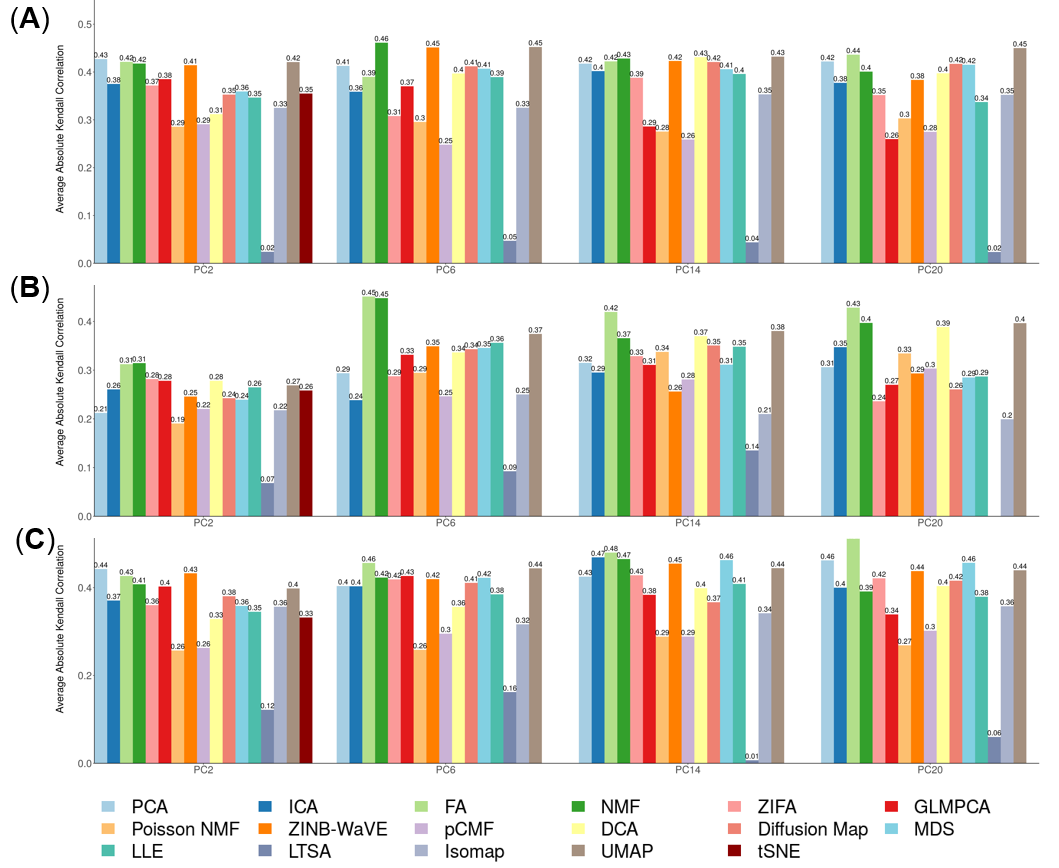

**Supplementary Figure 40. DR method performance evaluated by Kendal correlation for downstream trajectory inference analysis.** For each method, we used *Monocle3* to perform trajectory inference. We compared 17 DR methods (columns), including factor analysis (FA), principal component analysis (PCA), independent component analysis (ICA), Diffusion Map, nonnegative matrix factorization (NMF), Poisson NMF, zero-inflated factor analysis (ZIFA), zero-inflated negative binomial based wanted variation extraction (ZINB-WaVE), probabilistic count matrix factorization (pCMF), deep count autoencoder network (DCA), generalized linear model principal component analysis (GLMPCA), multidimensional scaling (MDS), locally linear embedding (LLE), local tangent space alignment (LTSA), Isomap, uniform manifold approximation and projection (UMAP), and t-distributed stochastic neighbor embedding (tSNE). We evaluated their performance on 14 real scRNAseq data sets. We used log2 Count transformation for the subset of DR methods that use normalized data. For each data set, we compared the four different number of low-dimensional data. The four numbers we used equal to 2, 6, 14, and 20 (x-axis). Some results of LTSA are empty because it is not compatible with Monocle3. Note that, for tSNE, we only extracted two low-dimensional components due to the limitation of the tSNE software.

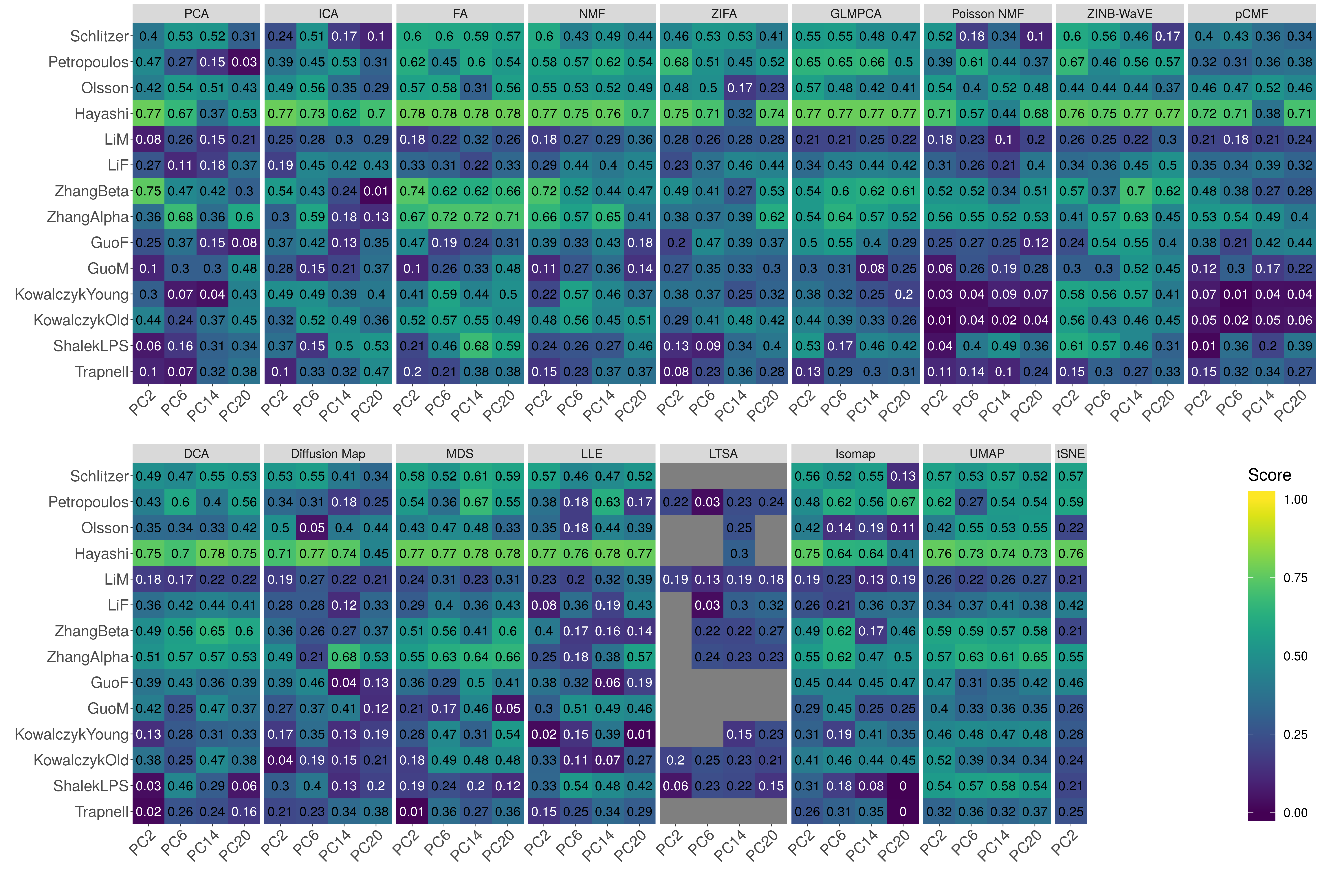

**Supplementary Figure 41. Bar plots show the average performance of different DR methods in downstream trajectory inference analysis.** The performance is evaluated by Kendal correlation averaged across 14 data sets. For each method, we used *Monocle3* to perform trajectory inference. We compared 17 DR methods, including factor analysis (FA; light green), principal component analysis (PCA; light blue), independent component analysis (ICA; blue), Diffusion Map (pink), nonnegative matrix factorization (NMF; green), Poisson NMF(light orange), zero-inflated factor analysis (ZIFA; light pink), zero-inflated negative binomial based wanted variation extraction (ZINB-WaVE; orange), probabilistic count matrix factorization (pCMF; light purple), deep count autoencoder network (DCA; yellow), generalized linear model principal component analysis (GLMPCA; red), multidimensional scaling (MDS; cyan), locally linear embedding (LLE; blue green), local tangent space alignment (LTSA; teal blue), Isomap (grey), uniform manifold approximation and projection (UMAP; brown), and t-distributed stochastic neighbor embedding (tSNE; dark red). We used log2 Count transformation for the subset of DR methods that use normalized data. For each data set, we compared the four different number of low-dimensional data. The four numbers we used equal to 2, 6, 14, and 20 (x-axis). Note that, for tSNE, we only extracted two low-dimensional components due to the limitation of the tSNE software.

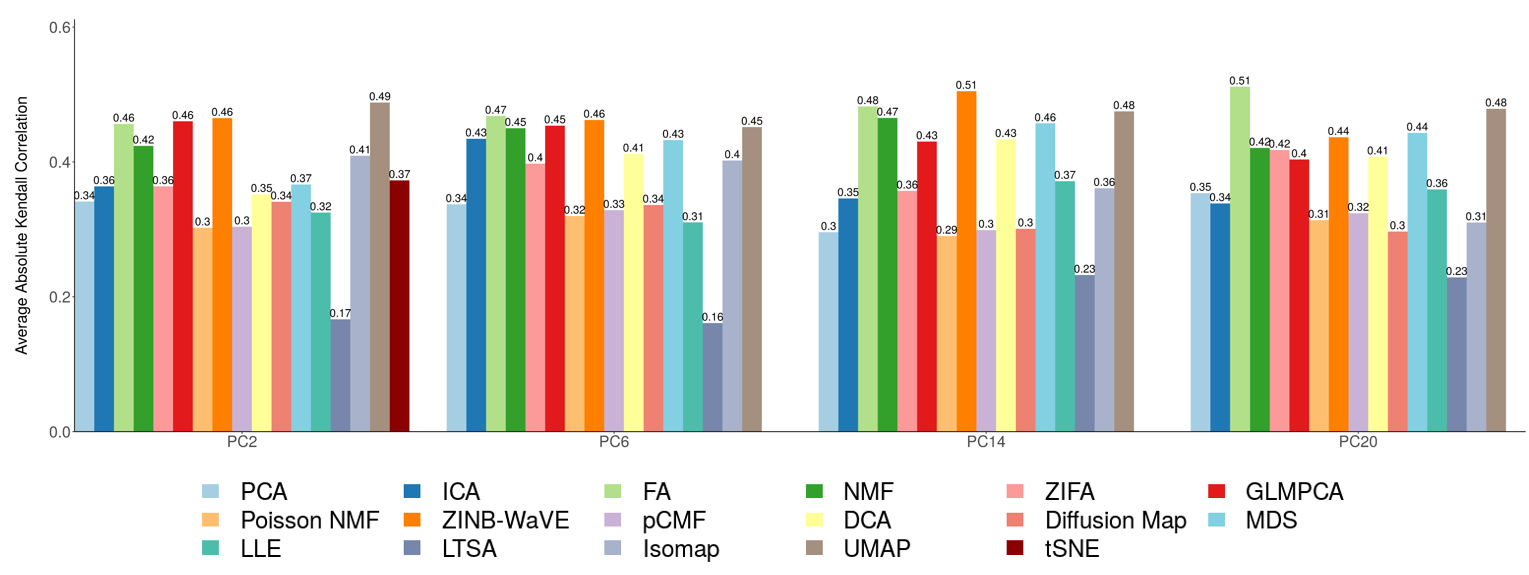

**Supplementary Figure 42. Trajectory visualization of different DR methods on ZhangBeta data set.** We compared 17 DR methods (columns), including factor analysis (FA), principal component analysis (PCA), independent component analysis (ICA), Diffusion Map, nonnegative matrix factorization (NMF), Poisson NMF, zero-inflated factor analysis (ZIFA), zero-inflated negative binomial based wanted variation extraction (ZINB-WaVE), probabilistic count matrix factorization (pCMF), deep count autoencoder network (DCA), generalized linear model principal component analysis (GLMPCA), multidimensional scaling (MDS), locally linear embedding (LLE), local tangent space alignment (LTSA), Isomap, uniform manifold approximation and projection (UMAP), and t-distributed stochastic neighbor embedding (tSNE). We used log2 Count transformation for the subset of DR methods that use normalized data. For each data set, we first reduced the dimension to 2, then applied UMAP to visualize the trajectory.

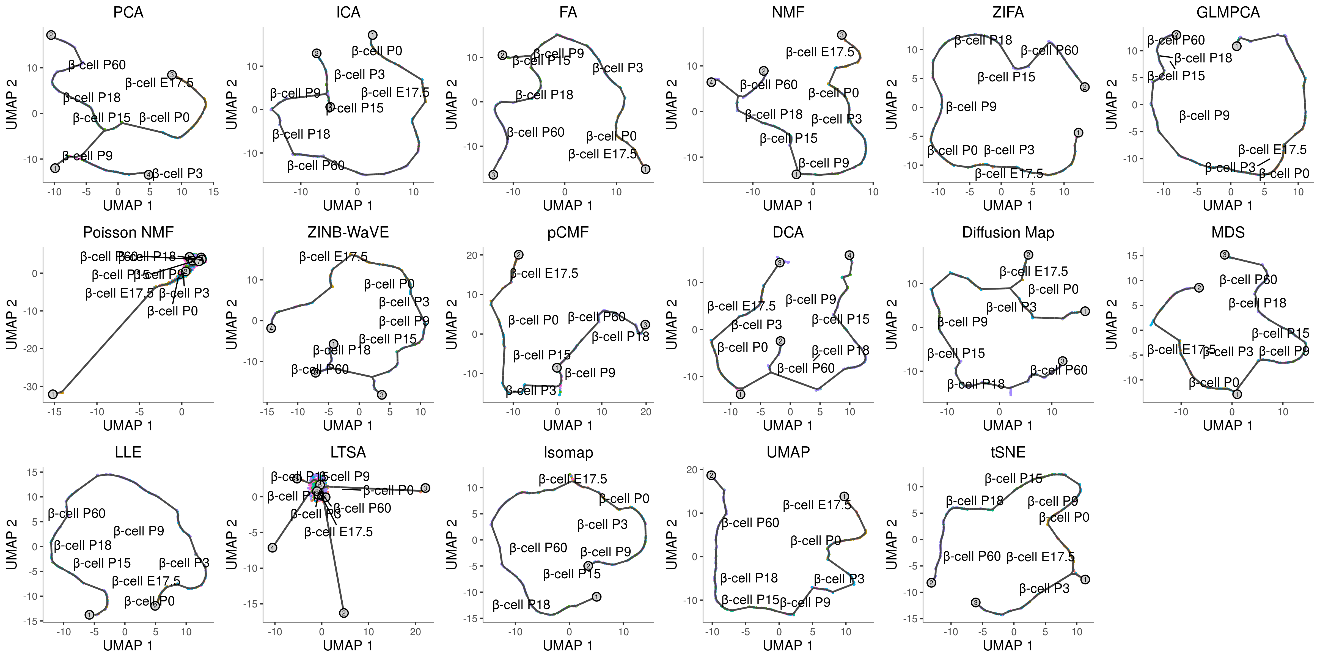

**Supplementary Figure 43. DR method performance with log-2 CPM transformed data for downstream trajectory inference analysis.** Performance is evaluated by Kendal correlation coefficient. For each method, we used *Slingshot* to perform trajectory inference on log2 CPM transformed data with *k*-means as the initial step. We compared 11 DR methods (columns), including factor analysis (FA), principal component analysis (PCA), independent component analysis (ICA), nonnegative matrix factorization (NMF), Diffusion Map, multidimensional scaling (MDS), locally linear embedding (LLE), local tangent space alignment (LTSA), Isomap, uniform manifold approximation and projection (UMAP), and t-distributed stochastic neighbor embedding (tSNE). We evaluated their performance on 14 real scRNAseq data sets (rows). For each data set, we compared the four different number of low-dimensional data. The four numbers we used equal to 2, 6, 14, and 20 (x-axis). Some results for LTSA or NMF are not shown (grey fills) because error occurred when we applied *Slingshot* on LTSA or NMF extracted low-dimensional components there. Note that, for tSNE, we only extracted two low-dimensional components due to the limitation of the tSNE software.

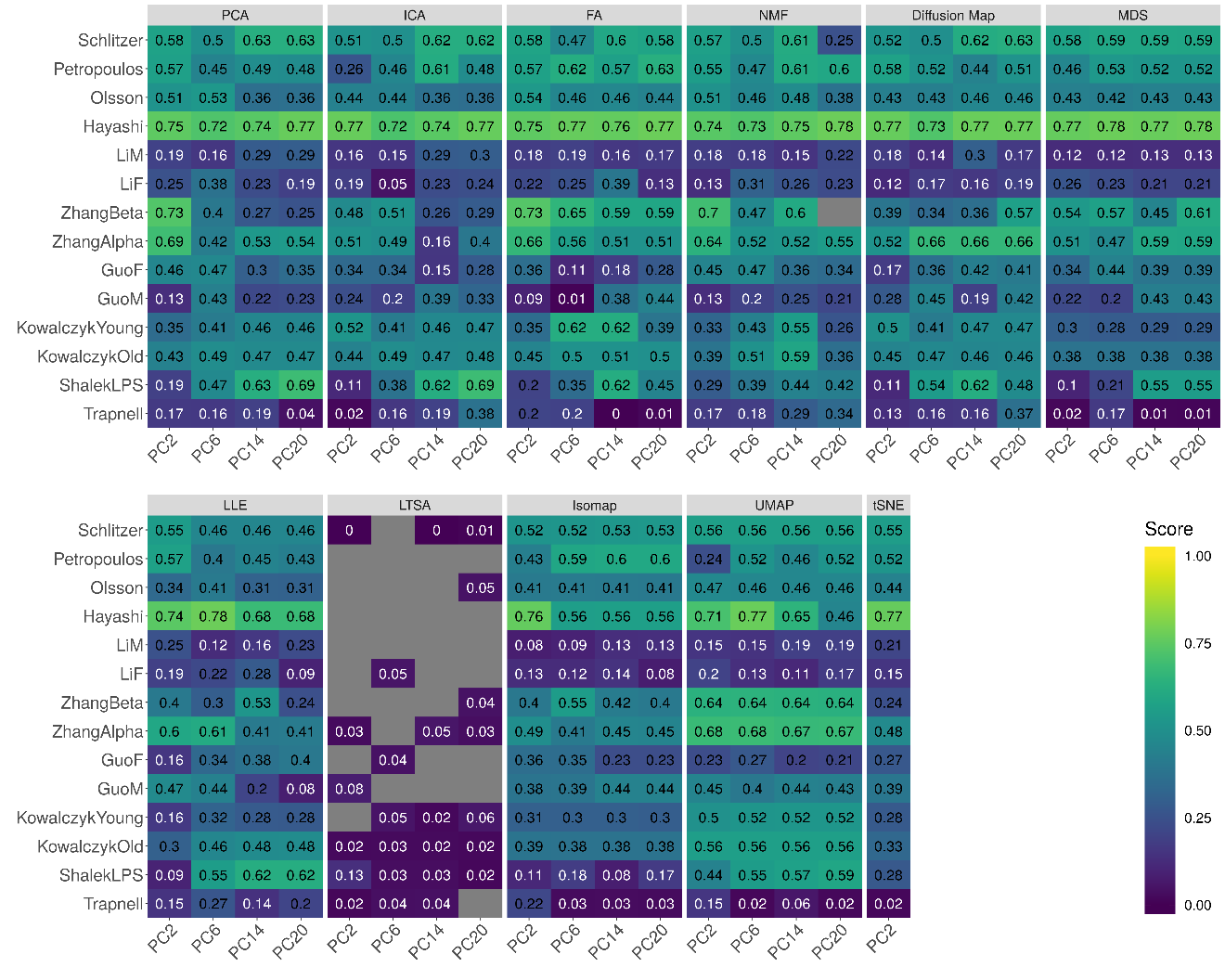

**Supplementary Figure 44. DR method performance with log-2 CPM transformed data for downstream trajectory inference analysis.** Performance is evaluated by Kendal correlation coefficient. For each method, we used *Slingshot* to perform trajectory inference on log2 CPM transformed data with hierarchical clustering as the initial step. We compared 11 DR methods (columns), including factor analysis (FA), principal component analysis (PCA), independent component analysis (ICA), Diffusion Map, nonnegative matrix factorization (NMF), multidimensional scaling (MDS), locally linear embedding (LLE), local tangent space alignment (LTSA), Isomap, uniform manifold approximation and projection (UMAP), and t-distributed stochastic neighbor embedding (tSNE). We evaluated their performance on 14 real scRNAseq data sets (rows). For each data set, we compared the four different number of low-dimensional data. The four numbers we used equal to 2, 6, 14, and 20 (x-axis). Some results for LTSA are not shown (grey fills) because error occurred when we applied *Slingshot* on LTSA extracted low-dimensional components there. Note that, for tSNE, we only extracted two low-dimensional components due to the limitation of the tSNE software.

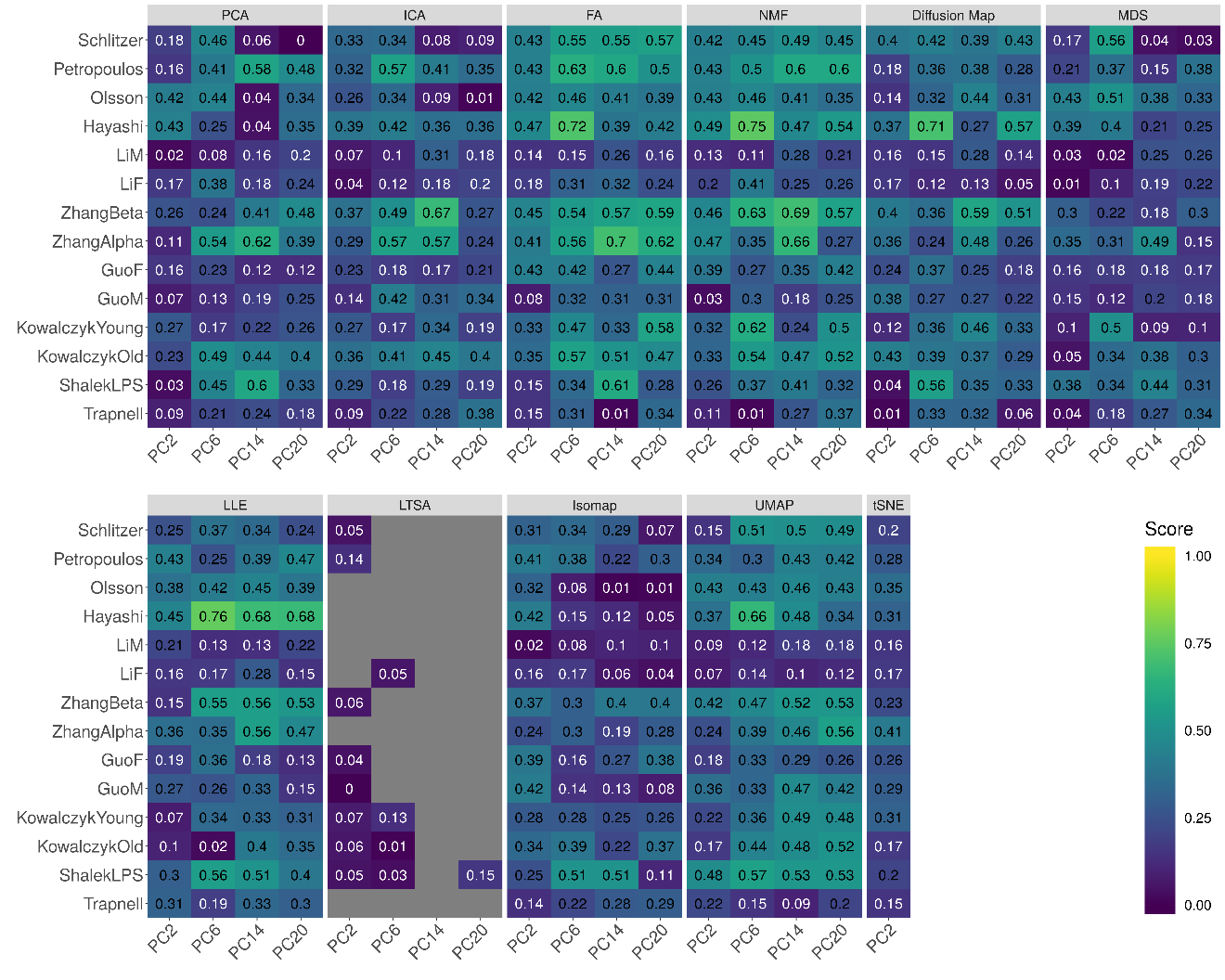

**Supplementary Figure 45. DR method performance with log-2 CPM transformed data for downstream trajectory inference analysis.** Performance is evaluated by Kendal correlation coefficient. For each method, we used *Slingshot* to perform trajectory inference on log2 CPM transformed data with Louvain method as the initial step. We compared 11 DR methods (columns), including factor analysis (FA), principal component analysis (PCA), independent component analysis (ICA), Diffusion Map, nonnegative matrix factorization (NMF), multidimensional scaling (MDS), locally linear embedding (LLE), local tangent space alignment (LTSA), Isomap, uniform manifold approximation and projection (UMAP), and t-distributed stochastic neighbor embedding (tSNE). We evaluated their performance on 14 real scRNAseq data sets (rows). For each data set, we compared the four different number of low-dimensional data. The four numbers we used equal to 2, 6, 14, and 20 (x-axis). Some results for LTSA are not shown (grey fills) because error occurred when we applied *Slingshot* on LTSA extracted low-dimensional components there. Note that, for tSNE, we only extracted two low-dimensional components due to the limitation of the tSNE software.

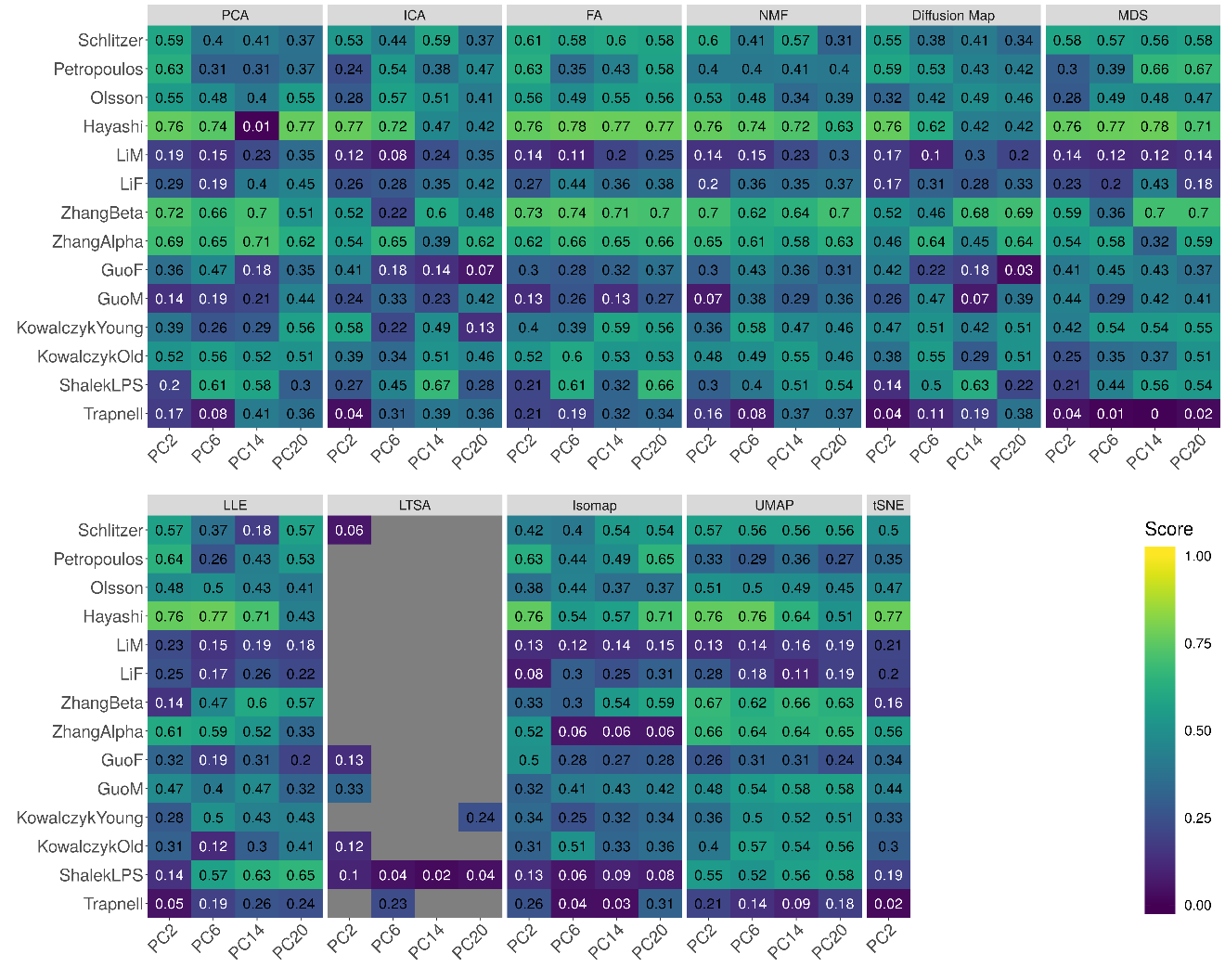

**Supplementary Figure 46. Bar plots show the average performance of a subset of DR methods that use log2 CPM transformed data in downstream trajectory inference analysis.** The performance is evaluated by Kendal correlation averaged across 14 data sets. For each method, we used *Slingshot* to perform trajectory inference with either *k*-means (**A**), hierarchical clustering (**B**) or Louvain method (**C**) as the initial step. We compared 11 DR methods, including factor analysis (FA; light green), principal component analysis (PCA; light blue), independent component analysis (ICA; blue), Diffusion Map (pink), nonnegative matrix factorization (NMF; green), multidimensional scaling (MDS; cyan), locally linear embedding (LLE; blue green), local tangent space alignment (LTSA; teal blue), Isomap (grey), uniform manifold approximation and projection (UMAP; brown), and t-distributed stochastic neighbor embedding (tSNE; dark red). For each data set, we compared the four different number of low-dimensional data. The four numbers we used equal to 2, 6, 14, and 20 (x-axis). Note that, for tSNE, we only extracted two low-dimensional components due to the limitation of the tSNE software.

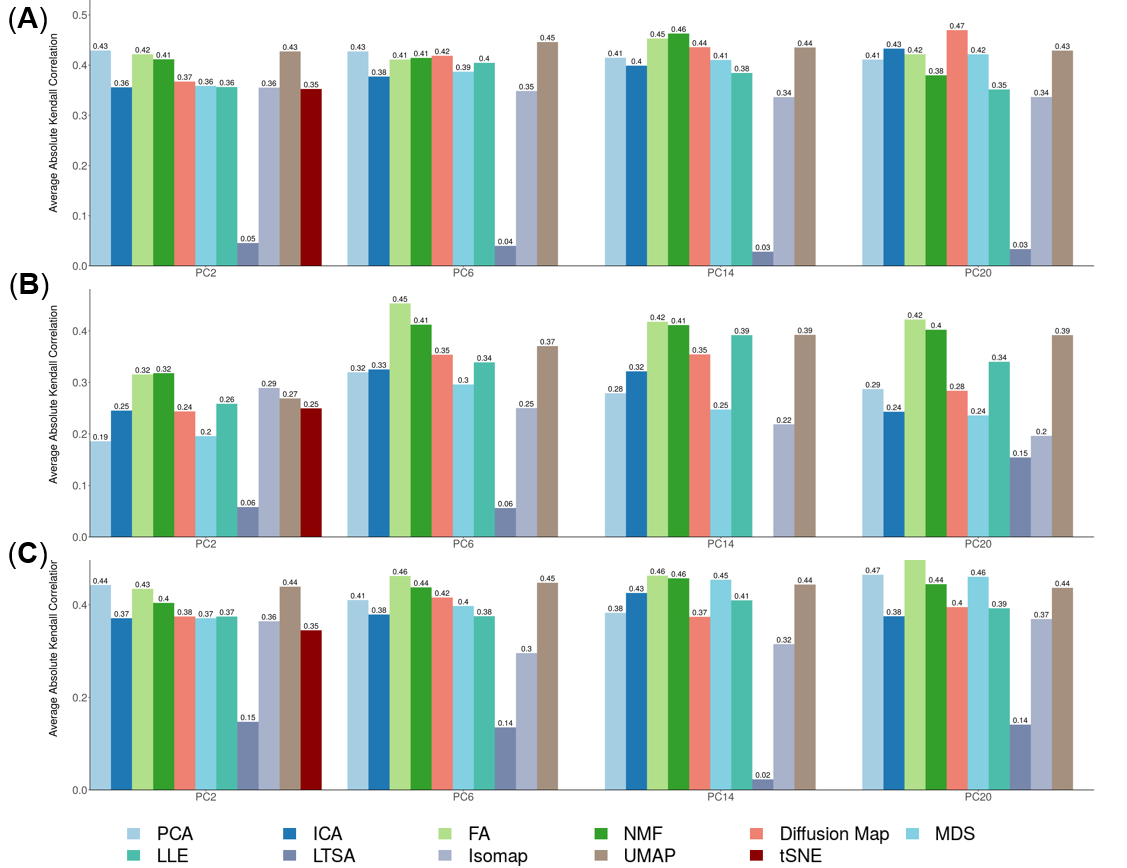

**Supplementary Figure 47. DR method performance with z-score transformed data for downstream trajectory inference analysis.** Performance is evaluated by Kendal correlation coefficient. For each method, we used *Slingshot* to perform trajectory inference on z-score transformed data with *k*-means as the initial step. We compared 10 DR methods (columns), including factor analysis (FA), principal component analysis (PCA), independent component analysis (ICA), Diffusion Map, multidimensional scaling (MDS), locally linear embedding (LLE), local tangent space alignment (LTSA), Isomap, uniform manifold approximation and projection (UMAP), and t-distributed stochastic neighbor embedding (tSNE). We evaluated their performance on 14 real scRNAseq data sets (rows). For each data set, we compared the four different number of low-dimensional data. The four numbers we used equal to 2, 6, 14, and 20. No results for NMF are shown in Figure because z-score transformation yields negative values. Some results for LTSA or ICA are not shown (grey fills) because error occurred when we applied *Slingshot* on LTSA or ICA extracted low-dimensional components there. Note that, for tSNE, we only extracted two low-dimensional components due to the limitation of the tSNE software.

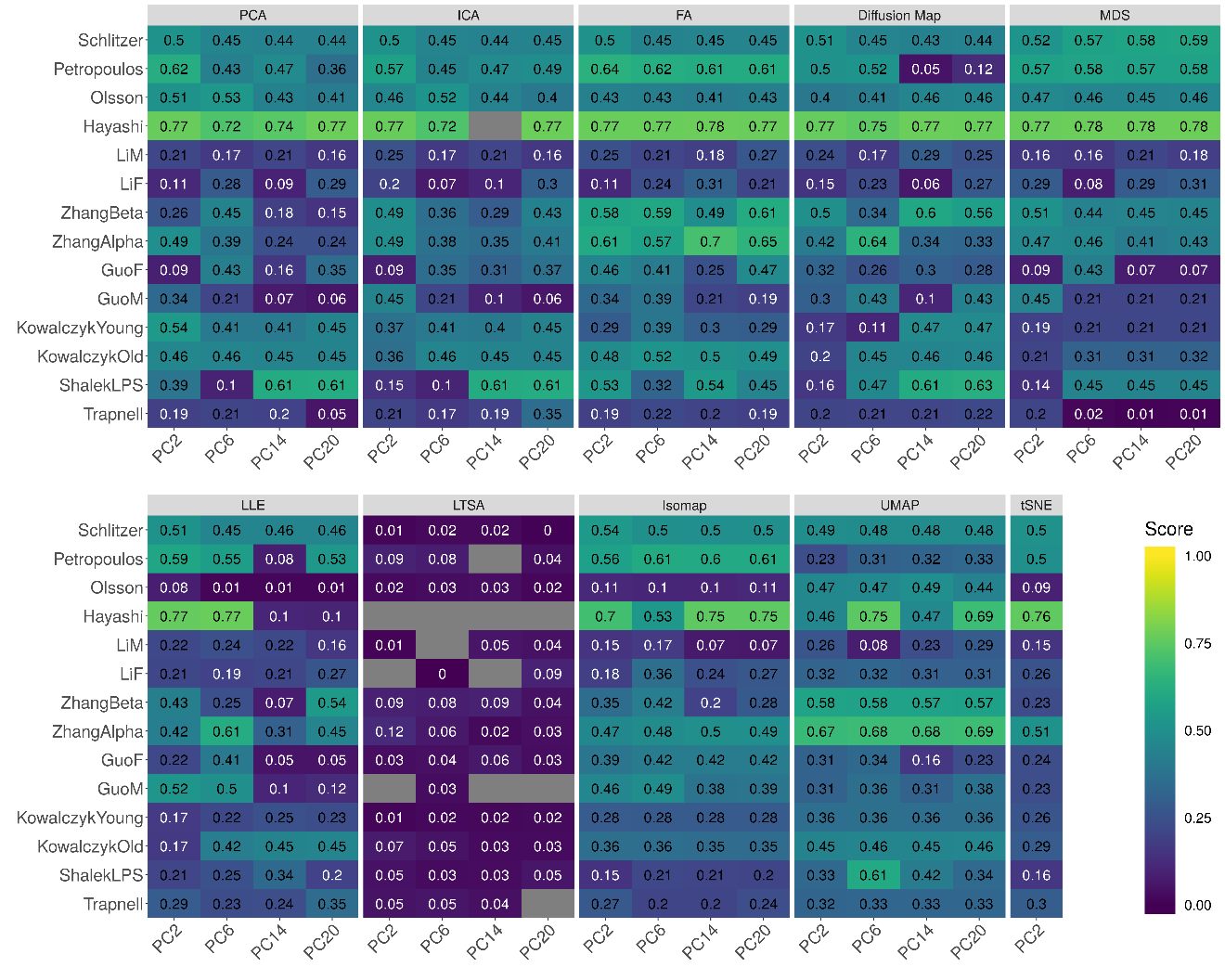

**Supplementary Figure 48. DR method performance with z-score transformed data for downstream trajectory inference analysis.** Performance is evaluated by Kendal correlation coefficient. For each method, we used *Slingshot* to perform trajectory inference on z-score transformed data with hierarchical clustering as the initial step. We compared 10 DR methods (columns), including factor analysis (FA), principal component analysis (PCA), independent component analysis (ICA), Diffusion Map, multidimensional scaling (MDS), locally linear embedding (LLE), local tangent space alignment (LTSA), Isomap, uniform manifold approximation and projection (UMAP), and t-distributed stochastic neighbor embedding (tSNE). We evaluated their performance on 14 real scRNAseq data sets (rows). For each data set, we compared the four different number of low-dimensional data. The four numbers we used equal to 2, 6, 14, and 20. No results for NMF are shown in Figure because z-score transformation yields negative values. Some results for LTSA are not shown (grey fills) because error occurred when we applied *Slingshot* on LTSA extracted low-dimensional components there. Note that, for tSNE, we only extracted two low-dimensional components due to the limitation of the tSNE software.

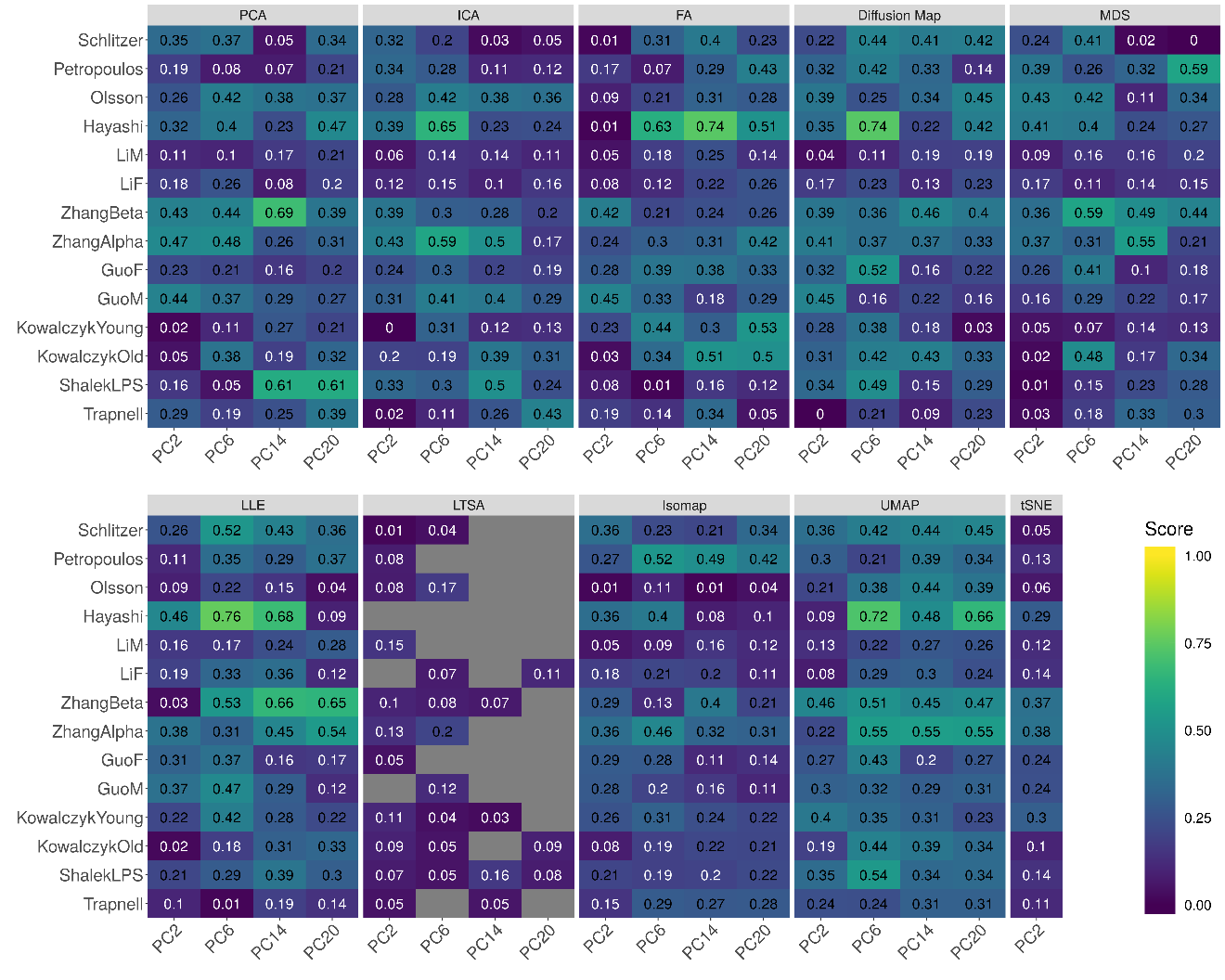

**Supplementary Figure 49. DR method performance with z-score transformed data for downstream trajectory inference analysis.** Performance is evaluated by Kendal correlation coefficient. For each method, we used *Slingshot* to perform trajectory inference on z-score transformed data with Louvain method as the initial step. We compared 10 DR methods (columns), including factor analysis (FA), principal component analysis (PCA), independent component analysis (ICA), Diffusion Map, multidimensional scaling (MDS), locally linear embedding (LLE), local tangent space alignment (LTSA), Isomap, uniform manifold approximation and projection (UMAP), and t-distributed stochastic neighbor embedding (tSNE). We evaluated their performance on 14 real scRNAseq data sets (rows). For each data set, we compared the four different number of low-dimensional data. The four numbers we used equal to 2, 6, 14, and 20. No results for NMF are shown in Figure because z-score transformation yields negative values. Some results for LTSA are not shown (grey fills) because error occurred when we applied *Slingshot* on LTSA extracted low-dimensional components there. Note that, for tSNE, we only extracted two low-dimensional components due to the limitation of the tSNE software.

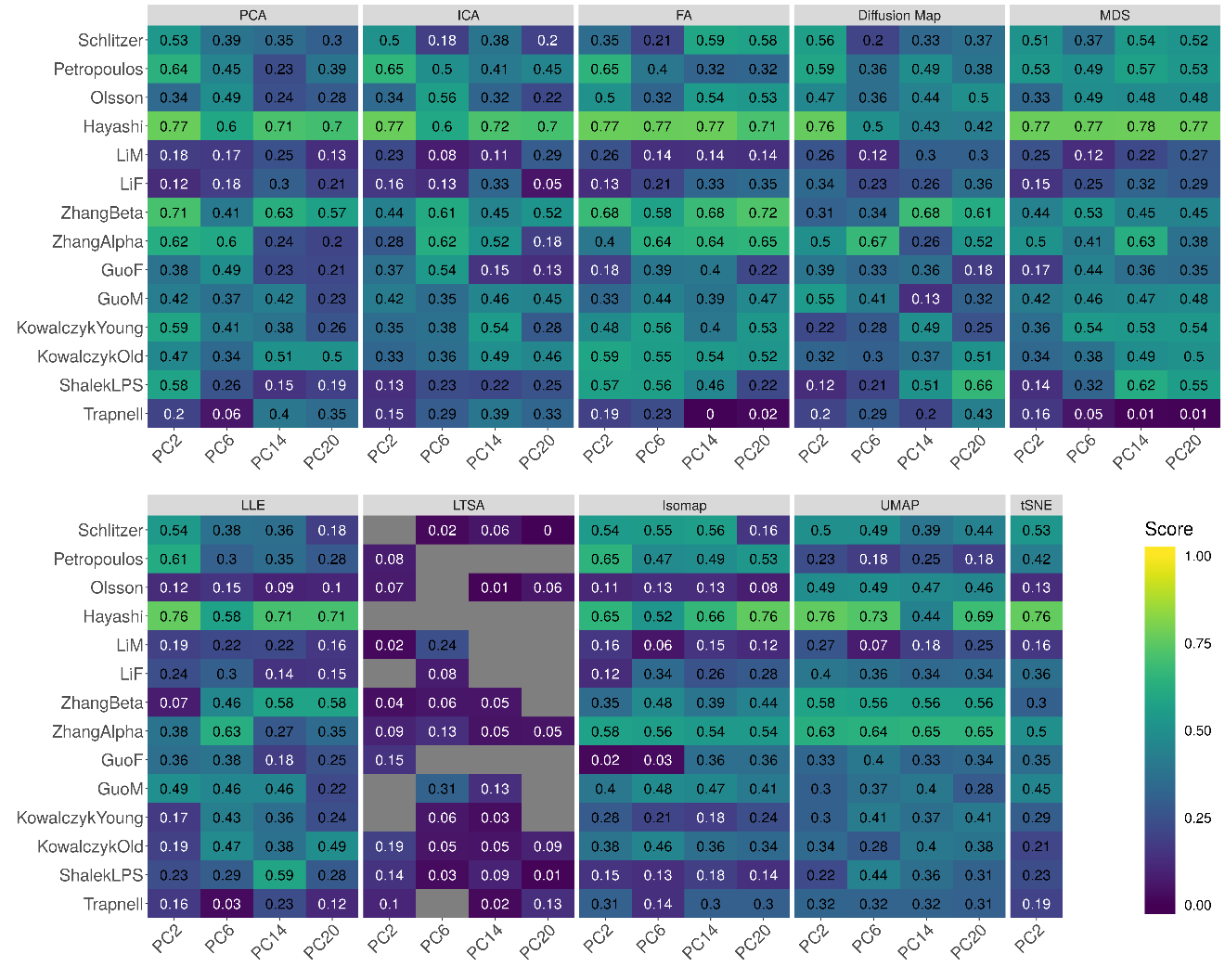

**Supplementary Figure 50. Bar plots show the average performance of a subset of DR methods that use z-score transformed data in downstream trajectory inference analysis.** The performance is evaluated by Kendal correlation averaged across 14 data sets. For each method, we used *Slingshot* to perform trajectory inference with either *k*-means (**A**), hierarchical clustering (**B**) or Louvain method (**C**) as the initial step. We compared 10 DR methods, including factor analysis (FA; light green), principal component analysis (PCA; light blue), independent component analysis (ICA; blue), Diffusion Map (pink), multidimensional scaling (MDS; cyan), locally linear embedding (LLE; blue green), local tangent space alignment (LTSA; teal blue), Isomap (grey), uniform manifold approximation and projection (UMAP; brown), and t-distributed stochastic neighbor embedding (tSNE; dark red). For each data set, we compared the four different number of low-dimensional data. The four numbers we used equal to 2, 6, 14, and 20 (x-axis). Note that, for tSNE, we only extracted two low-dimensional components due to the limitation of the tSNE software.

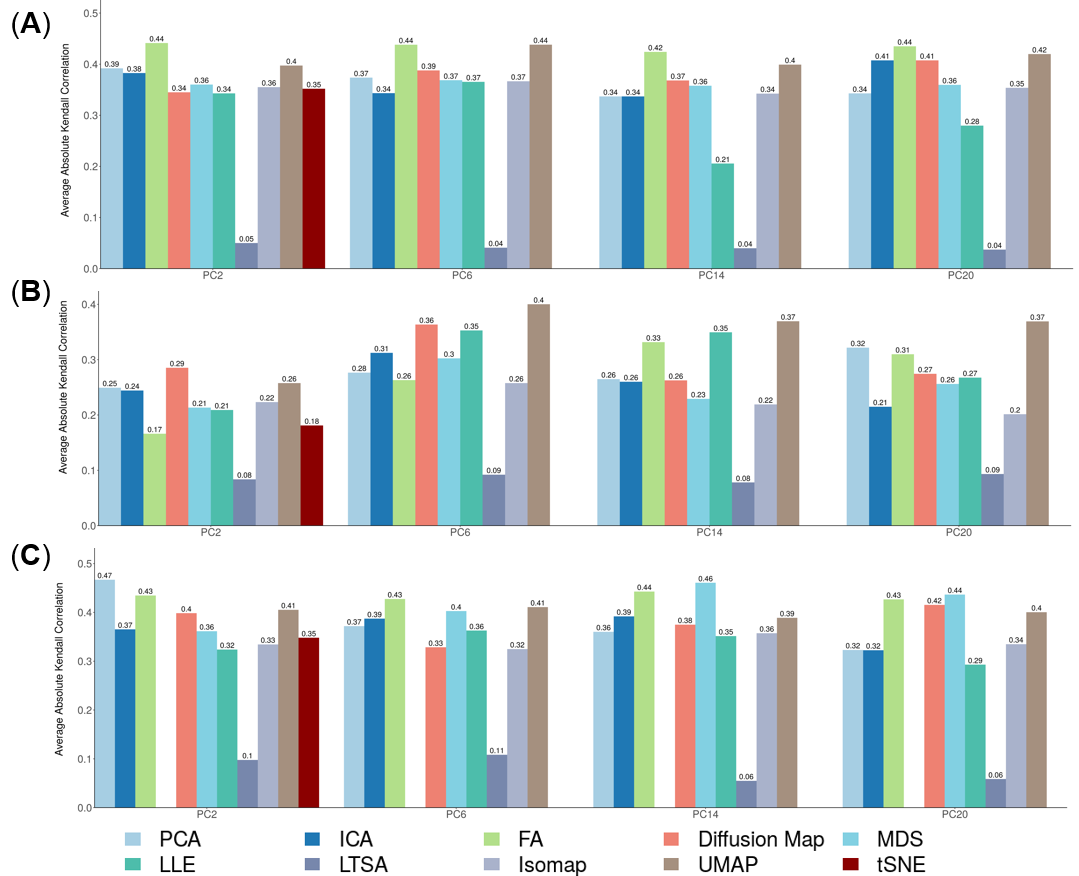

**Supplementary Figure 51. DR method performance with log-2 CPM transformed data for downstream trajectory inference analysis.** Performance is evaluated by Kendal correlation coefficient. For each method, we used *Monocle3* to perform trajectory inference on log2 CPM transformed data. We compared 11 DR methods (columns), including factor analysis (FA), principal component analysis (PCA), independent component analysis (ICA), Diffusion Map, nonnegative matrix factorization (NMF), multidimensional scaling (MDS), locally linear embedding (LLE), local tangent space alignment (LTSA), Isomap, and uniform manifold approximation and projection (UMAP), and t-distributed stochastic neighbor embedding (tSNE). We evaluated their performance on 14 real scRNAseq data sets (rows). For each data set, we compared the four different number of low-dimensional data. The four numbers we used equal to 2, 6, 14, and 20 (x-axis). Some results for LTSA are not shown (grey fills) because error occurred when we applied *Slingshot* on LTSA extracted low-dimensional components there. Note that, for tSNE, we only extracted two low-dimensional components due to the limitation of the tSNE software.

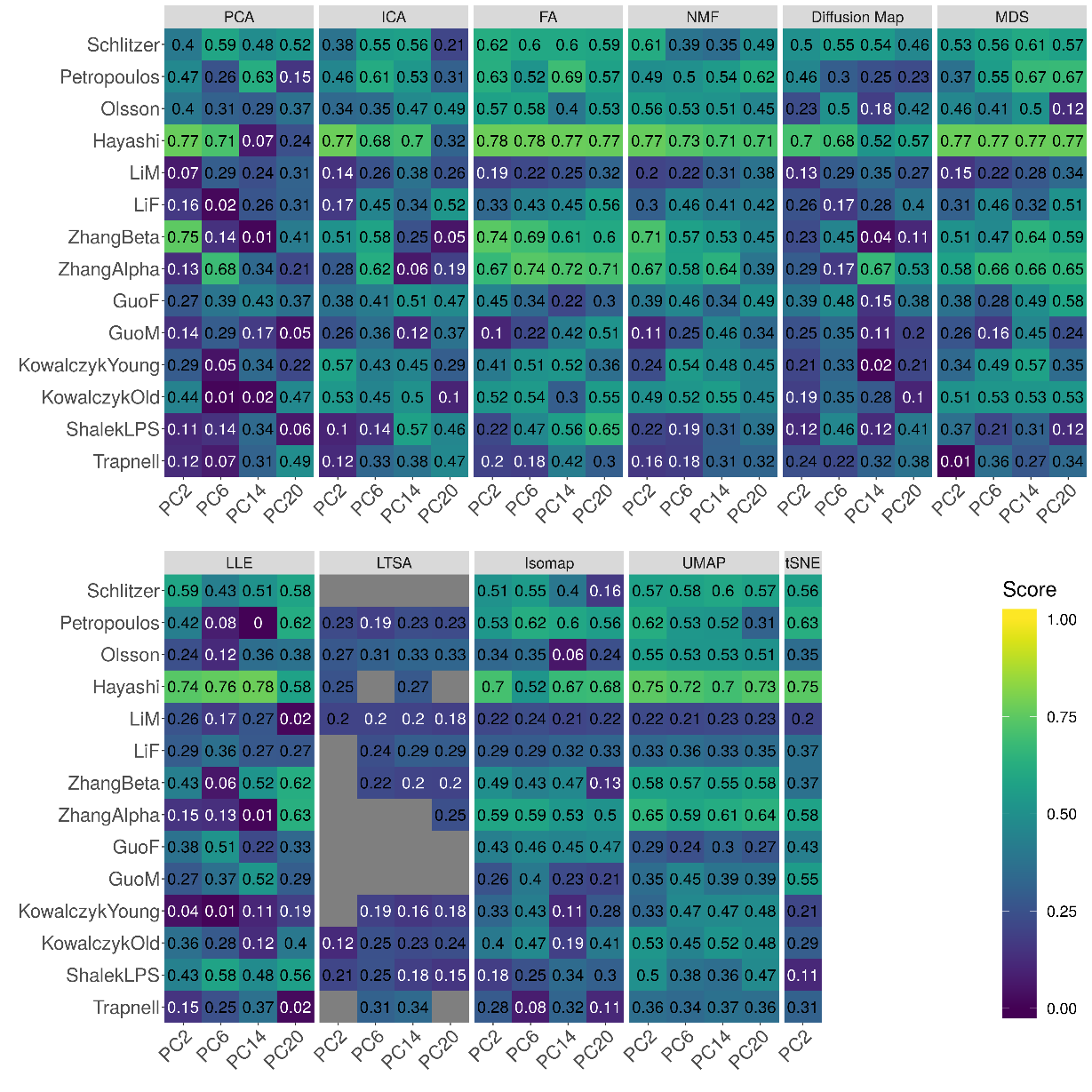

**Supplementary Figure 52. DR method performance with z-score transformed data for downstream trajectory inference analysis.** Performance is evaluated by Kendal correlation coefficient. For each method, we used *Monocle3* to perform trajectory inference on z-score transformed data. We compared 10 DR methods (columns), including factor analysis (FA), principal component analysis (PCA), independent component analysis (ICA), Diffusion Map, multidimensional scaling (MDS), locally linear embedding (LLE), local tangent space alignment (LTSA), Isomap, and uniform manifold approximation and projection (UMAP), and t-distributed stochastic neighbor embedding (tSNE). We evaluated their performance on 14 real scRNAseq data sets (rows). For each data set, we compared the four different number of low-dimensional data. The four numbers we used equal to 2, 6, 14, and 20 (x-axis). No results for NMF are shown in Figure because z-score transformation yields negative values. Some results for LTSA are not shown (grey fills) because error occurred when we applied *Slingshot* on LTSA extracted low-dimensional components there. Note that, for tSNE, we only extracted two low-dimensional components due to the limitation of the tSNE software.

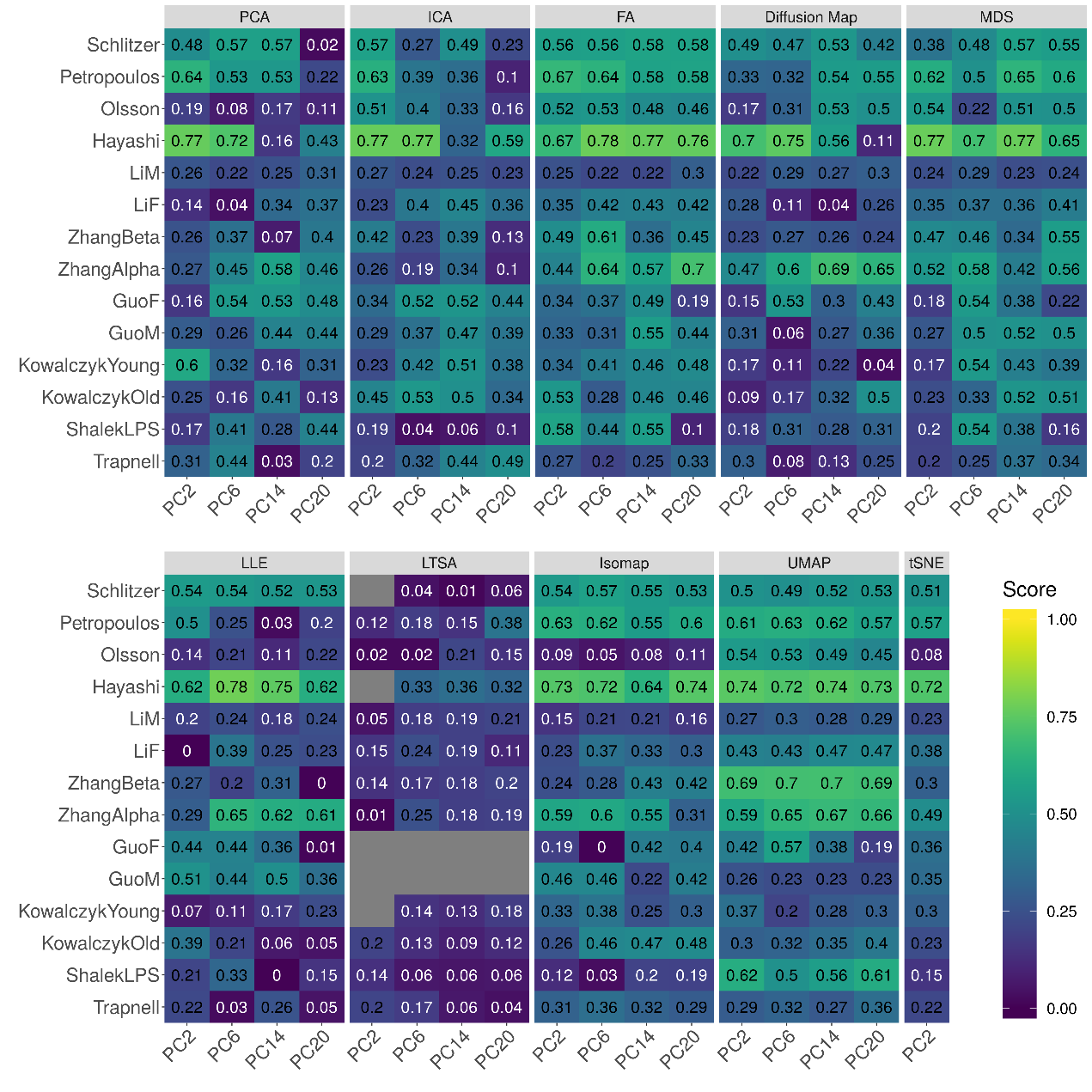

**Supplementary Figure 53. Bar plots show the average performance of a subset of DR methods in downstream trajectory inference analysis.** The performance is evaluated by Kendal correlation averaged across 14 data sets. For each method, we used *Monocle3* to perform trajectory inference on either log-2 transformation (**A**) or z-score transformation (**B**) data. We compared 11 DR methods, including factor analysis (FA; light green), principal component analysis (PCA; light blue), independent component analysis (ICA; blue), Diffusion Map (pink), nonnegative matrix factorization (NMF; green), multidimensional scaling (MDS; cyan), locally linear embedding (LLE; blue green), local tangent space alignment (LTSA; teal blue), Isomap (grey), uniform manifold approximation and projection (UMAP; brown), and t-distributed stochastic neighbor embedding (tSNE; dark red). For each data set, we compared the four different number of low-dimensional data. The four numbers we used equal to 2, 6, 14, and 20 (x-axis). Note that, for tSNE, we only extracted two low-dimensional components due to the limitation of the tSNE software.

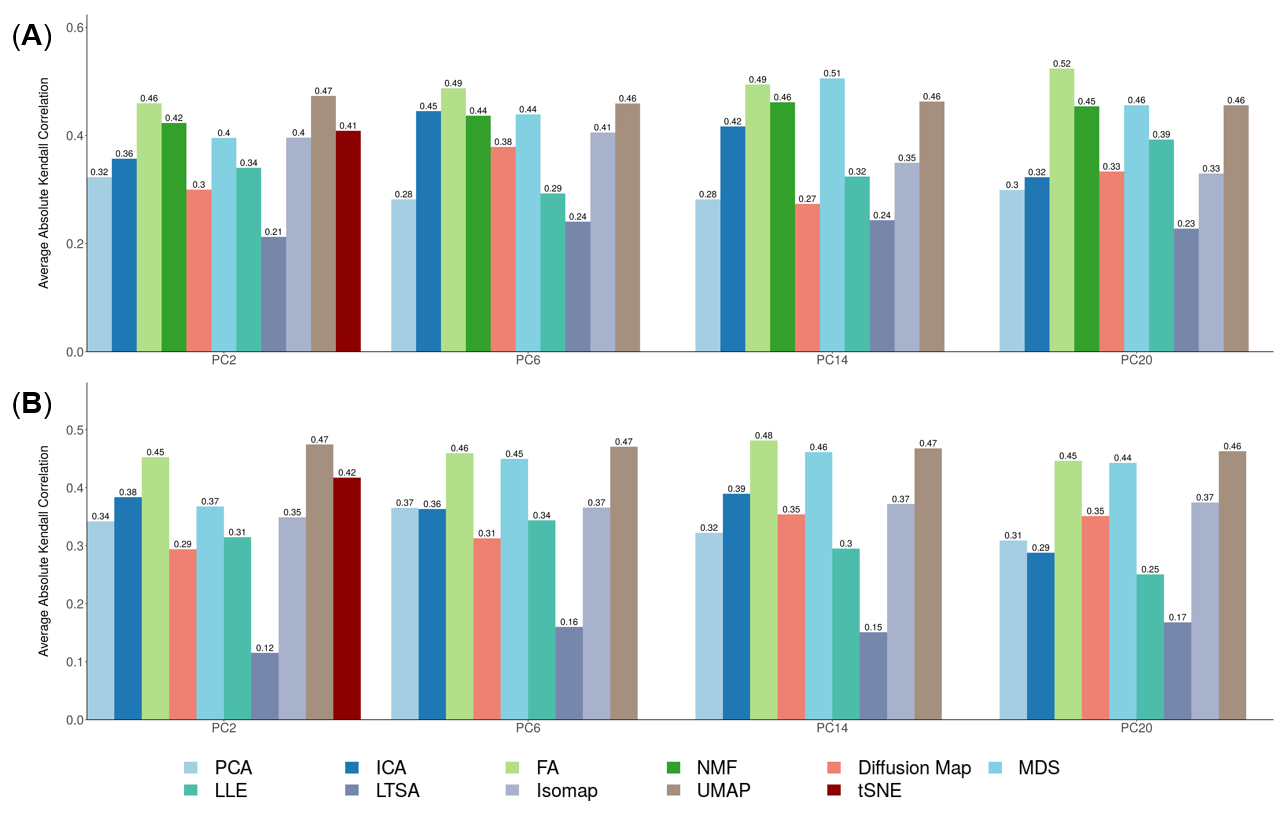

**Supplementary Figure 54. The stability and robustness of DR methods evaluated based on data split of** **the Hayashi data.** We compared 17 DR methods (columns), including factor analysis (FA), principal component analysis (PCA), independent component analysis (ICA), Diffusion Map, nonnegative matrix factorization (NMF), Poisson NMF, zero-inflated factor analysis (ZIFA), zero-inflated negative binomial based wanted variation extraction (ZINB-WaVE), probabilistic count matrix factorization (pCMF), deep count autoencoder network (DCA), generalized linear model principal component analysis (GLMPCA), multidimensional scaling (MDS), locally linear embedding (LLE), local tangent space alignment (LTSA), Isomap, uniform manifold approximation and projection (UMAP), and t-distributed stochastic neighbor embedding (tSNE). We split the Hayashi data randomly into two subsets with equal number of cells in each cell type (one colored red; the other colored green). We applied each DR method to each subset separately to obtain lineage results. Performance is measured by Kendall correlation after *Slingshot*. We performed 10 replicates for each data and show the performance results across these replicates using boxplot. For each data set, we compared the four different number of low-dimensional data. The four numbers we used equal to 2, 6, 14, and 20 (x-axis). Note that, for tSNE, we only extracted two low-dimensional components due to the limitation of the tSNE software.

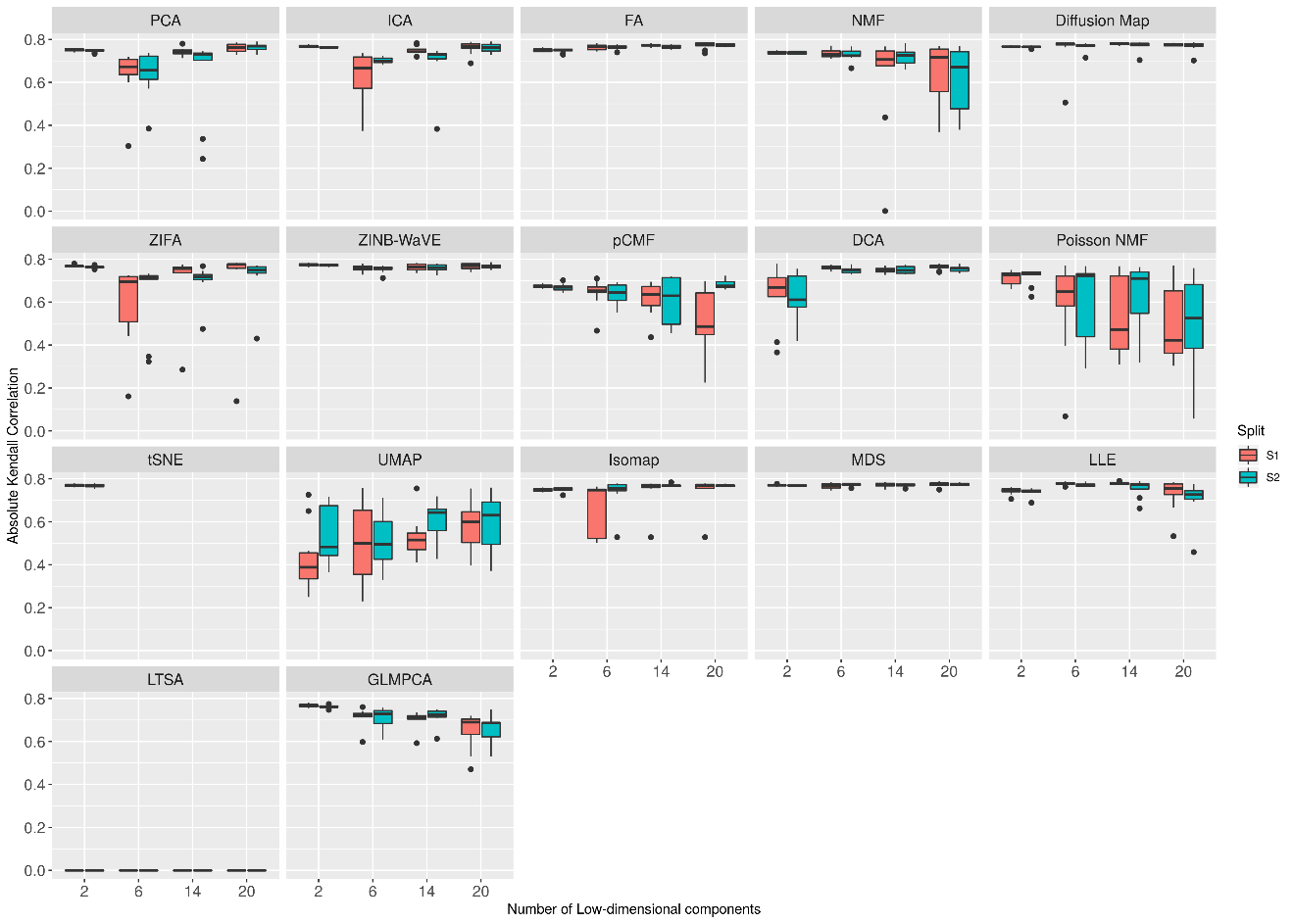

**Supplementary Figure 55. Bar plots show the performance of different DR methods with *k*-means clustering algorithm on raw Guo data set.** The performance is measured by NMI (**A**) or ARI (**B**). We compared 18 DR methods, including factor analysis (FA; light green), principal component analysis (PCA; light blue), independent component analysis (ICA; blue), Diffusion Map (pink), nonnegative matrix factorization (NMF; green), Poisson NMF(light orange), zero-inflated factor analysis (ZIFA; light pink), zero-inflated negative binomial based wanted variation extraction (ZINB-WaVE; orange), probabilistic count matrix factorization (pCMF; light purple), deep count autoencoder network (DCA; yellow), scScope (purple), generalized linear model principal component analysis (GLMPCA; red), multidimensional scaling (MDS; cyan), locally linear embedding (LLE; blue green), local tangent space alignment (LTSA; teal blue), Isomap (grey), uniform manifold approximation and projection (UMAP; brown) , and t-distributed stochastic neighbor embedding (tSNE; dark red). We used log2 count transformation for the subset of DR methods that use normalized data. For each data set, we compared the four different number of low-dimensional data. The four numbers we used equal to 0.5%, 1%, 2%, and 3% of the total number of cells. Note that, for tSNE, we only extracted two low-dimensional components due to the limitation of the tSNE software.

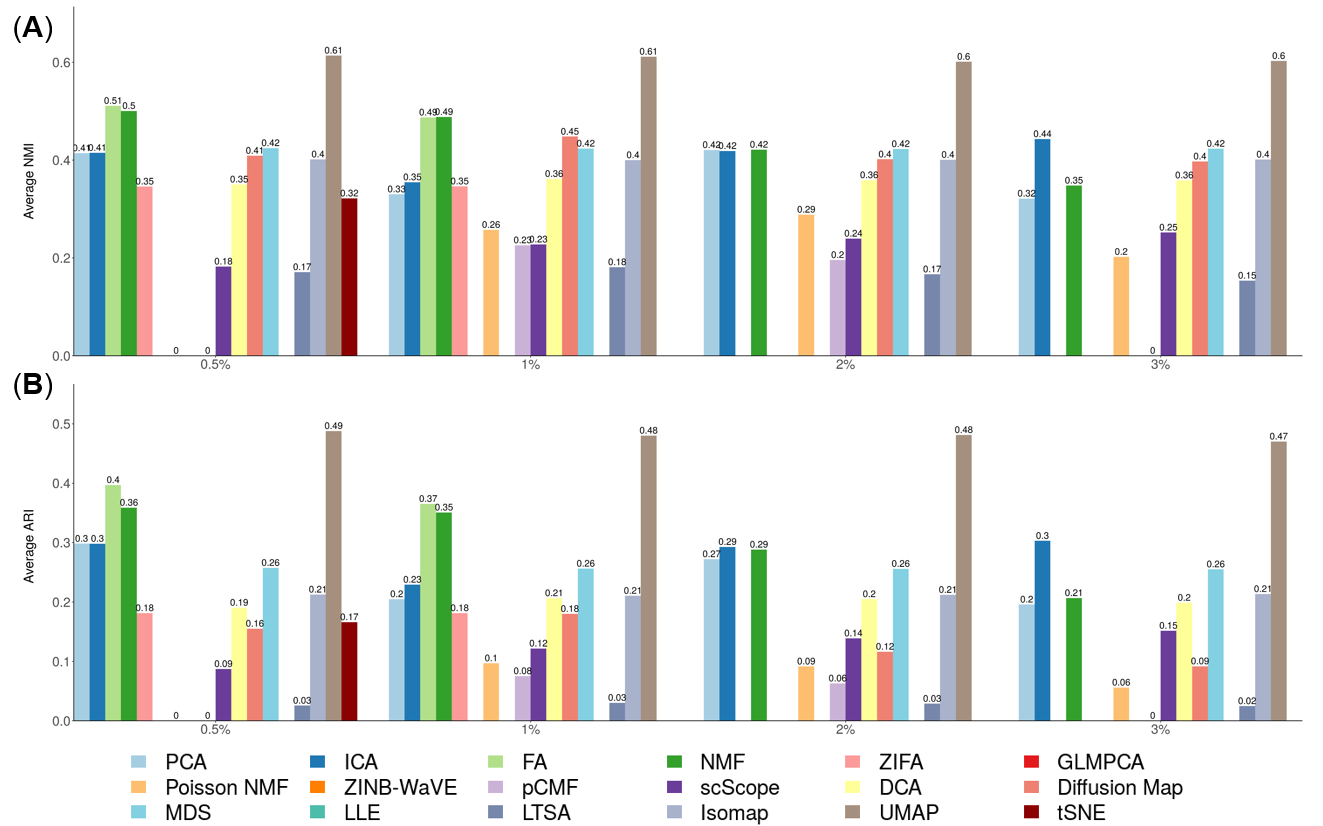

**Supplementary Figure 56. Bar plots show the performance of different DR methods with *k*-means clustering algorithm on Guo data set with subsampling procedure.** The performance is measured by NMI (**A**) or ARI (**B**). We compared 18 DR methods, including factor analysis (FA; light green), principal component analysis (PCA; light blue), independent component analysis (ICA; blue), Diffusion Map (pink), nonnegative matrix factorization (NMF; green), Poisson NMF(light orange), zero-inflated factor analysis (ZIFA; light pink), zero-inflated negative binomial based wanted variation extraction (ZINB-WaVE; orange), probabilistic count matrix factorization (pCMF; light purple), deep count autoencoder network (DCA; yellow), scScope (purple), generalized linear model principal component analysis (GLMPCA; red), multidimensional scaling (MDS; cyan), locally linear embedding (LLE; blue green), local tangent space alignment (LTSA; teal blue), Isomap (grey), uniform manifold approximation and projection (UMAP; brown) , and t-distributed stochastic neighbor embedding (tSNE). We used log2 count transformation for the subset of DR methods that use normalized data. For each data set, we compared the four different number of low-dimensional data. The four numbers we used equal to 0.5%, 1%, 2%, and 3% of the total number of cells. Note that, for tSNE, we only extracted two low-dimensional components due to the limitation of the tSNE software.

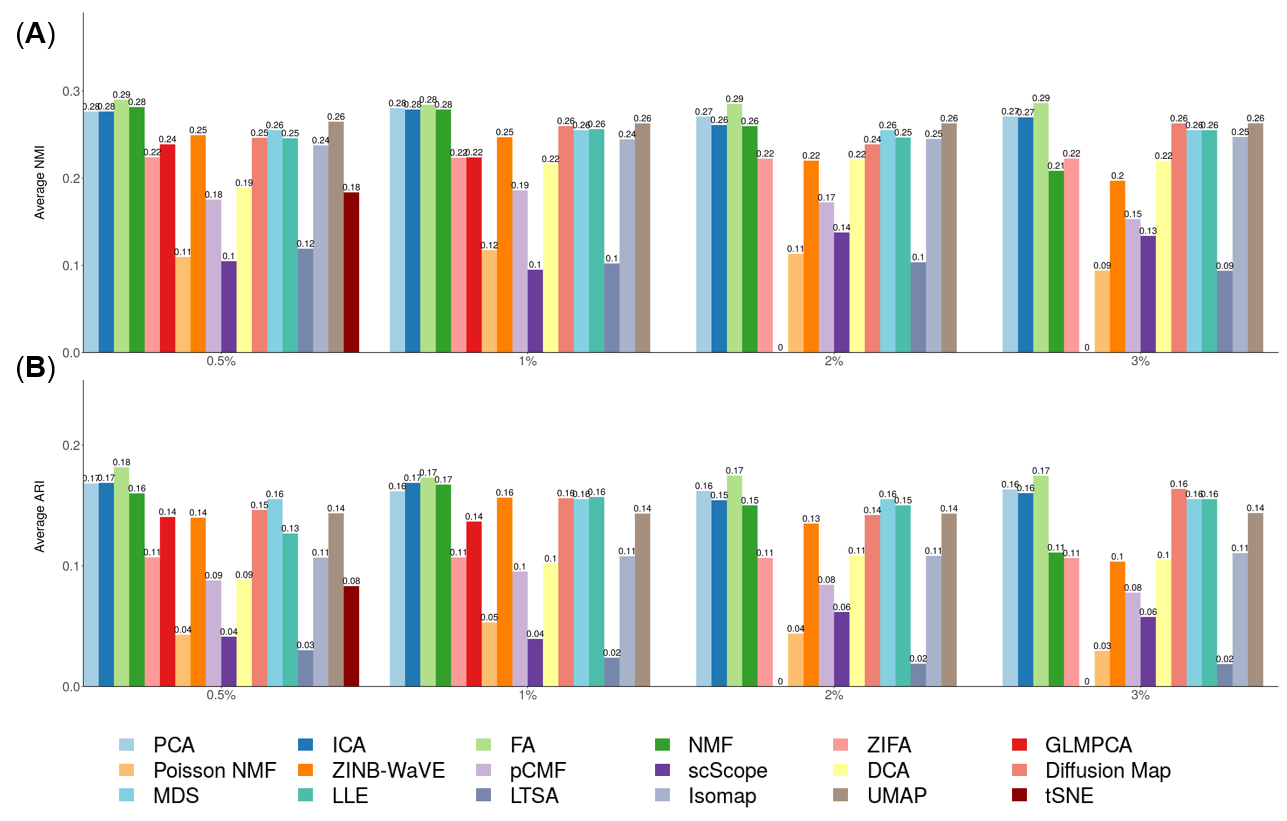

**Supplementary Figure 57. Bar plots show the performance of different DR methods with *Slingshot* on Cao data set with subsampling procedure.** The performance is measured by NMI (**A**) or ARI (**B**). We compared 17 DR methods, including factor analysis (FA; light green), principal component analysis (PCA; light blue), independent component analysis (ICA; blue), Diffusion Map (pink), nonnegative matrix factorization (NMF; green), Poisson NMF(light orange), zero-inflated factor analysis (ZIFA; light pink), zero-inflated negative binomial based wanted variation extraction (ZINB-WaVE; orange), probabilistic count matrix factorization (pCMF; light purple), deep count autoencoder network (DCA; yellow), generalized linear model principal component analysis (GLMPCA; red), multidimensional scaling (MDS; cyan), locally linear embedding (LLE; blue green), local tangent space alignment (LTSA; teal blue), Isomap (grey), and uniform manifold approximation and projection (UMAP; brown), and t-distributed stochastic neighbor embedding (tSNE; dark red). We used log2 count transformation for the subset of DR methods that use normalized data. For each data set, we compared the four different number of low-dimensional data. The four numbers we used equal to 0.5%, 1%, 2%, and 3% of the total number of cells. Note that, for tSNE, we only extracted two low-dimensional components due to the limitation of the tSNE software.

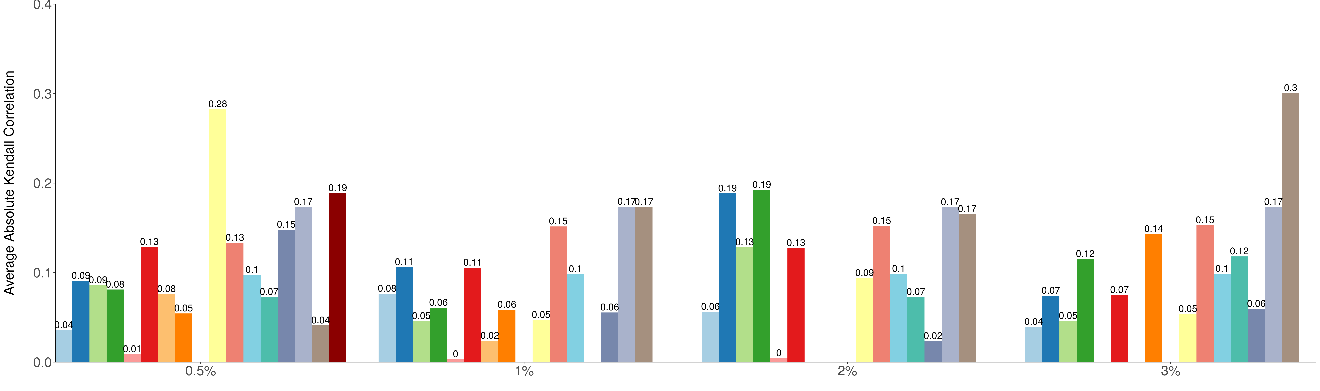

**Supplementary Table 1**. **List of 15 scRNAseq data sets used for benchmarking DR methods through cell clustering.** The table contains data set ID (1^st^ column), data set name (2^nd^ column), species (3^rd^ column), experimental platform (4^th^ column), year of publication (5^th^ column), how the true clusters were determined (6^th^ column), number of genes (7^th^ column), number of cells (8^th^ column), number of cell types (9^th^ column) and unique molecular identifiers (UMI; 10^th^ column). We colored different categories in each column by different colors.

| ID | Data set | Species | Protocol | Year | Determination | # Genes | # Cells | #Cluster | UMI |
| --- | --- | --- | --- | --- | --- | --- | --- | --- | --- |
| GSE111108 | FreytagGold | human | 10xGenomics Chromium | 2018 | FACS | 58,302 | 925 | 3 | Yes |
| GSE115189 | GSE115189Silver | human | 10xGenomics Chromium | 2018 | clustering | 58,302 | 2,590 | 11 | Yes |
| SRP073767 | Zhengmix4eq | human | 10xGenomics GemCode | 2017 | FACS | 15,568 | 3,994 | 4 | Yes |
|  | Zhengmix4uneq | human | 10xGenomics GemCode | 2017 | FACS | 16,443 | 6,498 | 4 | Yes |
|  | PBMC3k | human | 10xGenomics GemCode | 2017 | FACS | 58,302 | 3,205 | 11 | Yes |
|  | PBMC4k | human | 10xGenomics Chromium | 2017 | FACS | 58,302 | 4,292 | 11 | Yes |
| GEO67835 | Darmanis | human | Smart-Seq2 | 2017 | clustering | 22,085 | 420 | 4 | No |
| GSE73727 | Pancreatic | human | Smart-Seq2 | 2016 | FACS | 139,918 | 60 | 6 | No |
| GSE84133 | Baron | human | inDrop | 2016 | clustering | 14,878 | 1,886 | 13 | Yes |
| GSE75748 | ChuBatch1 | human | SMARTer | 2016 | FACS | 19,097 | 350 | 5 | No |
|  | ChuBatch2 | human | SMARTer | 2016 | FACS | 19,097 | 425 | 6 | No |
| GSE60361 | Zeisel | mouse | Smart-Seq | 2015 | clustering | 19,972 | 1,800 | 7 | No |
| GSE60749 | Kumar | mouse | Smart-Seq | 2014 | culture conditions | 45,159 | 246 | 3 | No |
| SRP073808 | Koh | human | SMARTer | 2016 | FACS | 48,981 | 531 | 9 | No |
| GSE99254 | Guo | human | Smart-Seq2 | 2018 | FACS | 18,178 | 12,346 | 25 | No |

**Supplementary Table 2**. **List of 15 scRNAseq data sets used for benchmarking DR methods through trajectory inference.** The table contains data set ID (1^st^ column), data set name (2^nd^ column), species (3^rd^ column), experimental platform (4^th^ column), year of publication (5^th^ column), how the true lineage were determined (6^th^ column), number of genes (7^th^ column), number of cells (8^th^ column), unique molecular identifiers (UMI; 9^th^ column) and lineage type (10^th^ column). We colored different categories in each column by different colors.

| ID | Data set | Species | Platform | Year | Determination | # Genes | # Cells | UMI | Lineage Type |
| --- | --- | --- | --- | --- | --- | --- | --- | --- | --- |
| GSE60783 | Schlitzer | mouse | Fluidigm | 2015 | FACS | 4480 | 238 | No | Linear |
| E-MTAB-3929 | Petropoulos | human | Smart-Seq2 | 2016 | timeseries | 8,772 | 1,289 | No | Linear |
| GSE86146 | LiM | human | Smart-Seq2 | 2017 | timeseries | 4,777 | 649 | No | Linear |
|  | LiF | human | Smart-Seq2 | 2017 | timeseries | 5,319 | 666 | No | Linear |
| GSE87375 | ZhangBeta | mouse | Smart-Seq2 | 2017 | timeseries | 6,138 | 562 | No | Linear |
|  | ZhangAlpha | mouse | Smart-Seq2 | 2017 | timeseries | 6,138 | 322 | No | Linear |
| GSE63818 | GuoF | human | Tang et. al. | 2015 | timeseries | 8,772 | 100 | No | Linear |
|  | GuoM | human | Tang et. al. | 2015 | timeseries | 8,772 | 166 | No | Linear |
| GSE59114 | KowalczykYoung | mouse | Smart-Seq | 2015 | FACS | 2,227 | 493 | No | Linear |
|  | KowalczykOld | mouse | Smart-Seq | 2015 | FACS | 2,815 | 873 | No | Linear |
| GSE98664 | Hayashi | mouse | RamDA-seq | 2018 | timeseries | 23,658 | 414 | No | Linear |
| GSE48968 | ShalekLPS | mouse | Smart-Seq | 2014 | timeseries | 4,158 | 504 | No | Linear |
| GSE52529 | Trapnell | human | SMARTer | 2014 | timeseries | 8,772 | 290 | No | Linear |
| GSE70245 | Olsson | mouse | SMARTer | 2016 | FACS | 3,594 | 316 | No | Linear |
| GSE119945 | Cao | mouse | Sci-RNA-seq3 | 2019 | timeseries | 26,183 | 2,058,652 | Yes | Linear |

**Download links for 15 data sets used in cell clustering:**

**FreytagGold**: The data set consists of three human lung adenocarcinoma cell lines, HCC827, H1975, and H2228, which were cultured separately and mixed together for sequencing [1]. The dataset was sequenced by 10x Genomics Chromium Controller. Cell subpopulation information was established by the applying demuxlet (version 0.0.1) on single nucleotide variants (SNVs) to explore the genetic difference and then identify near absolute truth labels of the cells. Raw counts were downloaded from <https://www.ncbi.nlm.nih.gov/geo/query/acc.cgi?acc=GSE111108>.

**GSE115189Silver**: The data set consists of five fresh human peripheral blood mononuclear cells (PBMCs) [1]. The dataset was sequenced by 10x Genomics Chromium Controller. Cell subpopulation information was established by the applying cell labeling approach from 10x Genomics. The cell type labeling obtained through this approach has high consistency with literature. Raw counts were downloaded from <https://www.ncbi.nlm.nih.gov/geo/query/acc.cgi?acc=GSE115189>.

**Zhengmix4eq**: The data set consists of four pre-sorted cell types (B-cells, naive cytotoxic T-cells, CD14 monocytes, regulatory T-cells) that were combined with equal proportions, with 1,000 cells per cell type [2]. The data were sequenced by 10x Genomics, and the Cell Ranger Single-Cell Software Suite was used to perform sample demultiplexing, barcode processing and single-cell 3’ gene counting. The cell type for each cell were known through the pre-sorting process and is considered as the underlying truth. Raw UMI counts were downloaded from <https://support.10xgenomics.com/single-cell-gene-expression/datasets>.

**Zhengmix4uneq**: The data set consists of four pre-sorted cell types (B-cells, naive cytotoxic T-cells, CD14 monocytes, regulatory T-cells) that were combined with unequal proportions, with 1,000, 500, 2,000, and 3,000 cells for the four cell types, respectively [2]. The data were sequenced by 10x Genomics, and the Cell Ranger single cell software suite was used to perform sample demultiplexing, barcode processing and single-cell 3’ gene counting. The cell type for each cell were known through the pre-sorting process and is considered as the underlying truth. Raw UMI counts were downloaded from <https://support.10xgenomics.com/single-cell-gene-expression/datasets>.

**PBMC4K**: The data set consists of 11 cell types obtained by in fluorescence-activated cell sorting (FACS). The data were generated with an earlier version of the microfluidics instrument, the 10x Genomics GemCode Controller. Raw counts were downloaded from <https://support.10xgenomics.com/single-cell-geneexpression/datasets/1.2.0/pbmc4k>.

**PBMC3k**: The data set consists of 11 cell types obtained by in fluorescence-activated cell sorting (FACS). The data set consists of 11 cell types obtained by in fluorescence-activated cell sorting (FACS). These 11 cell types include CD14+ Monocyte (446 cells), CD19+ B (406 cells), CD34+ (17 cells), CD4+/CD25 T Reg (198 cells), CD4+/CD45RA+/CD25- Naive T (242 cells), CD4+/CD45RO+ Memory (465 cells), CD4+ T Helper2 (76 cells), CD56+ NK (428 cells), CD8+/CD45RA+ Naive Cytotoxic (510 cells), CD8+ Cytotoxic T(398 cells), Dendritic (19 cells). The data were generated with the latest instrument, the 10x Genomics Chromium Controller. Raw counts were downloaded from <https://support.10xgenomics.com/single-cell-gene-expression/datasets/2.1.0/pbmc4k>.

**Darmanis**: This data set originally contains 10 distinct cell groups. Clusters 1-8 consist of adult brain cells, whereas cluster 9 and 10 consist of fetal brain cells. We excluded two mixed groups of cells that is probably due to contamination of each cell with pieces of myelin debris or noisy gene expression profile [3], remaining 8 major brain cell types: astrocytes (62 cells), oligodendrocytes (38 cells), oligodendrocyte precursor cells (OPCs; 18 cells), neurons (131 cells), microglia (16 cells), fetal quiescent (110 cells), fetal replicating (25 cells) and endothelial (20 cells). First, we used an unbiased approach to sort all 466 individual cells into distinct groups defined by the entirety of their molecular signatures. Raw count data were downloaded from <https://www.ncbi.nlm.nih.gov/geo/query/acc.cgi?acc=GSE67835>.

**Koh**: This data set comprises of 9 different cell types that include H7 hESCs, H7-derived anterior primitive streak populations, H7-derived mid primitive streak populations, H7-derived lateral mesoderm, H7-derived FACS-purified GARP+ cardiac mesoderm, H7-derived FACS-purified DLL1+ paraxial mesoderm populations, H7-derived day 3 early somite progenitor populations, H7-derived dermomyotome populations, and H7-derived FACS-purified PDGFRα+ sclerotome populations. RNA was extracted from either whole cell populations or alternatively, cell subsets purified by fluorescence activated cell sorting (FACS). Raw data can be downloaded from <https://www.ncbi.nlm.nih.gov/sra?term=SRP073808>. In our comparison, we directly downloaded the processed data from conquer [4].

**Baron**: This data set characterized pancreatic cells from human donors and was sequenced by the inDrop platform. Raw count data were downloaded from <https://www.ncbi.nlm.nih.gov/geo/query/acc.cgi?acc=GSE84133>.

**ChuBatch1**: This is the batch 1 part of Chu et al. data [5]. Chu et at al contains four lineage-specific progenitor cells derived from human pluripotent stem cells (hPSCs) and were sequenced by Fluidigm C1 platform. Raw data was downloaded from <https://www.ncbi.nlm.nih.gov/geo/query/acc.cgi?acc=GSE75748>.

**ChuBatch2**: This is the batch 2 part of Chu et al. data [5]. Chu et at al contains four lineage-specific progenitor cells derived from human pluripotent stem cells (hPSCs) and were sequenced by Fluidigm C1 platform. Raw data was downloaded from <https://www.ncbi.nlm.nih.gov/geo/query/acc.cgi?acc=GSE75748>.

**Pancreatic**: This data set contains human islet cells from one human donor [6]. This dataset was sequenced by Smart-seq2 and raw data were download from <https://www.ncbi.nlm.nih.gov/geo/query/acc.cgi?acc=GSE73727>. We excluded undefined cells in scRNAseq samples and focused on the remaining 60 cells in our evaluation.

**Zeisel**: This data set contains cells of the somatosensory cortex and hippocampal CA1 region from mouse brains and was sequenced by the Fluidigm C1 platform. Raw counts were downloaded from <http://linnarssonlab.org/cortex/>.

**Kumar**: This data set contains three populations of mouse embryonic stem cells (mESCs): (1) v6.5 mESCs cultured in serum+LIF media (183 individuals); (2) v6.5 mESCs cultured in 2i+LIF media (94 individuals); (3) Dgcr8-/-mESCs (constructed in a v6.5 background), which lack mature miRNAs due to knockout of a miRNA processing factor in the cells, are cultured in serum+LIF (84 individuals). Raw count data can be downloaded from <https://www.ncbi.nlm.nih.gov/geo/query/acc.cgi?acc=GSE60749>. In our experiments, we directly downloaded the processed data from the conquer website [4].

**Guo**: This data set contains 25 populations of 12346 T cells from the tumour, adjacent normal tissues and peripheral blood from 14 treatment-naïve NSCLC patients [7]. Following the original paper, we removed CD4 iNKT, CD4 MAIT, DN diverse, DN iNKT, DN MAIT, DP diverse, and other cell subpopulations, and retained 9055 cells with 16 cell subpopulations, including CD4_C1-CCR7 (531 cells), CD4_C2-ANXA1 (665 cells), CD4_C3-GNLY (433 cells), CD4_C4-CD69 (1084 cells), CD4_C5-EOMES (161 cells), CD4_C6-GZMA (668 cells), CD4_C7-CXCL13 (342 cells), CD4_C8-FOXP3 (427 cells), CD4_C9-CTLA4 (939 cells), CD8_C1-LEF1 (303 cells), CD8_C2-CD28 (206 cells), CD8_C3-CX3CR1 (1192 cells), CD8_C4-GZMK (674 cells), CD8_C5-ZNF683 (832 cells), CD8_C6-LAYN (493 cells), and CD8_C7-SLC4A10 (105 cells). Raw count data can be downloaded from https://www.ncbi.nlm.nih.gov/geo/query/acc.cgi?acc=GSE99254.

**Download link for 15 data sets used in trajectory inference:**

In our comparison, all scRNAseq data sets with trajectory information were downloaded from <https://zenodo.org/record/1443566#.XNV25Y5KhaR> [8]. The detailed information of each data sets is as follows:

**Schlitzer**: This data set contains three cell types at three different developmental stages: common dendritic cell progenitor (CDP; 89 cells); macrophage and DC precursor (MDP; 57 cells); and Pre-DC (92 cell), from the first ontogeny of dendritic cell (DC) development in the bone marrow. Raw data can be downloaded from <https://www.ncbi.nlm.nih.gov/geo/query/acc.cgi?acc=GSE60783>.

**Petropoulos**: This data set is from a human embryo development study. The data includes the sequenced transcriptomes of 1,529 individual cells from 88 human preimplantation embryos. The data includes five developmental stages: E3 (81 cells), E4 (190 cells), E5 (377 cells), E6 (415 cells), and E7 (466 cells).The read count expression matrices for all cells is available on ArrayExpress: <https://www.omicsdi.org/dataset/arrayexpress-repository/E-MTAB-3929>.

**Trapnell:** This data set contains human skeletal muscle myoblasts cells collected over a time-course of serum-induced differentiation. At four different time points, cells were captured and processed with the Fluidigm C1 platform followed by Illumina sequencing. The underlying lineage in the data is known and was determined based on the collection time for the cells. The raw data is available at <https://www.ncbi.nlm.nih.gov/geo/query/acc.cgi?acc=GSE52529>.

**Olsson:** This data set was performed on stem/multipotent progenitors (LSK; lin−Sca1+c-Kit+; 92 cells), common myeloid progenitors (CMP; 92 cells), and granulocyte monocyte progenitors (GMP; 57 cells) that included granulocytic precursors. To isolate LSK, CMP and GMP, lineage stained cells were stained with: Streptavidin APC-Cy7 (Becton, Dickinson and Company), CD16/32-PerCp-ef710 (clone 93, eBioscience), CD117-APC (clone 2B8, Becton, Dickinson and Company), Sca-1-Pe-Cy7 (clone D7, Becton, Dickinson and Company) and CD34-BV421 (clone RAM34, Becton, Dickinson and Company). GMP and CMP gates were set using CD34 FMO. The data set can be downloaded from <https://www.ncbi.nlm.nih.gov/geo/query/acc.cgi?acc=GSE70245>.

**LiM:** This data set was collected from male fetal gonads with intact morphology and reasonable cell viability from embryos spanning a time period of 4 weeks (4W) to 26W. The stages of human embryos in this study were calculated from the estimated fertilization time, rather than last menstruation bleeding time. The clinicians who obtained the samples made this determination. The number of replicates for each developmental stage was no more than three. In total, 12 male embryos were collected: 21W samples had three biological replicates; 10W, 19W samples each had two biological replicates; and 4W, 9W, 12W, 20W and 25W each had one biological replicate. Raw data can be downloaded from <https://www.ncbi.nlm.nih.gov/geo/query/acc.cgi?acc=GSE86146>.

**LiF:** This data set was collected from female fetal gonads with intact morphology and reasonable cell viability from embryos spanning 4W to 26W. The stages of human embryos in this study were calculated from the estimated fertilization time, rather than last menstruation bleeding time. The clinicians who obtained the samples made this determination. The number of replicates for each developmental stage was no more than three. In total, 17 female embryos were in this study: 5W, 18W, 20W, 23W and 24W samples had two biological replicates; and 7W, 8W, 10W, 11W, 12W 14W and 26W had one biological replicate. Raw data can be downloaded from <https://www.ncbi.nlm.nih.gov/geo/query/acc.cgi?acc=GSE86146>.

**ZhangBeta:** This data set contains mouse pancreatic beta cells collected from seven developmental stages. The overall goal of this study was to define the roadmaps for pancreatic β-cell development. Specifically, we performed single-cell RNAseq at various developmental stages of E17.5, P0, P3, P9, P15, P18 and P60 of β-cells. Cells are sorted through FACS and followed by single-cell RNAseq using SMART-seq2. The lineage was determined based on the cell collection time. The raw data is available at <https://www.ncbi.nlm.nih.gov/geo/query/acc.cgi?acc=GSE87375>.

**ZhangAlpha:** This data set contains mouse pancreatic alpha cells collected from six developmental stages. The overall goal of this study was to define the roadmaps for pancreatic α-cell development. Specifically, we performed single-cell RNAseq at various developmental stages of E17.5, P0, P9, P15, P18 and P60 of α- cells. Cells are sorted through FACS and followed by single-cell RNAseq using SMART-seq2. The lineage was determined based on the cell collection time. The raw data is available at <https://www.ncbi.nlm.nih.gov/geo/query/acc.cgi?acc=GSE87375>.

**GuoF:** This data set contains human female primordial germ cells collected at five different time points. Cells are sorted through magnetic-activated cell sorting (MACS) or FACS and followed by scRNAseq using a modified single-cell cDNA amplification method [9] . The lineage was determined based on the cell collection time. The raw data is available at <https://www.ncbi.nlm.nih.gov/geo/query/acc.cgi?acc=GSE63818>.

**GuoM:** This data set contains human male primordial germ cells collected at five different time points. Cells are sorted through magnetic-activated cell sorting (MACS) or FACS and followed by single-cell RNAseq using a modified single-cell cDNA amplification method [9]. The lineage was determined based on the cell collection time. The raw data is available at <https://www.ncbi.nlm.nih.gov/geo/query/acc.cgi?acc=GSE63818>

**KowalczykYoung:** This data set contains mouse long-term hematopoietic stem cells, short-term hematopoietic stem cells and multi-potent progenitors from young mice (2~3 months of age). Cells are sorted through FACS and followed by single-cell RNAseq using Smart-seq. The lineage was determined based on the FACS. The raw data is available at <https://www.ncbi.nlm.nih.gov/geo/query/acc.cgi?acc=GSE59114>.

**KowalczykOld:** This data set contains mouse long-term hematopoietic stem cells, short-term hematopoietic stem cells and multi-potent progenitors from old mice (>20 months). Cells are sorted through FACS and followed by scRNAseq using Smart-seq. The lineage was determined based on FACS. The raw data is available at <https://www.ncbi.nlm.nih.gov/geo/query/acc.cgi?acc=GSE59114>.

**Hayashi:** The data set is from a cell differentiation time series study with RamDA-seq samples. The data contains 414 RamDA-seq samples of mouse ES cells that were collected at 0 (89 cells), 12 (67 cells), 24 (89 cells), 48 (79 cells), and 72 (90 cells) hours after the induction of cell differentiation into PrE cells. The samples were sequenced by Smart-Seq technology. The counts data can be downloaded from <https://www.ncbi.nlm.nih.gov/geo/query/acc.cgi?acc=GSE98664>.

**ShalekLPS:** This data set contains mouse dendritic-cells along a time-course of LPS-stimulation. At five different time points, cells were captured and processed with the Fluidigm C1 platform followed by scRNAseq using Smart-seq. The lineage was determined based on the cell collection time. The raw data is available at <https://www.ncbi.nlm.nih.gov/geo/query/acc.cgi?acc=GSE48968>.

**Cao:** This data set contains over 2 million cells derived from 61 mouse embryos staged between 9.5 and 13.5 days of gestation, including 0.8× at E9.5 (200,000 cells per embryo; 152,000 profiled across all replicates), 0.3× at E10.5 (1.1 million cells per embryo; 378,000 profiled), 0.2× at E11.5 (2.6 million cells per embryo; 616,000 profiled), 0.08× at E12.5 (6 million cells per embryo; 475,000 profiled) and 0.03× at E13.5 (13 million cells per embryo; 437,000 profiled) [10]. In our experiments, we randomly selected 10,000 cells from each stage to perform the trajectory analysis. The raw data is available at https://www.ncbi.nlm.nih.gov/geo/query/acc.cgi?acc=GSE119945.
